# Myeloid Gi signaling acts as a weight-independent immunometabolic switch controlling systemic insulin sensitivity

**DOI:** 10.64898/2026.03.28.713834

**Authors:** Seema Kuldeep, Harender Yadav, Seema Riyaz, Soumita Bhaumik, Shruti Agarwal, Aishi S Satapathy, Simran Singh, Ashish Kumar, Sudipta Paul, Monika Patel, Mriganka Sarkar, Raashidha Farhath, Sonal Amit, Rashmi Parihar, Hamim Zafar, Prem N Yadav, Suresh Kumar, Sai P Pydi

## Abstract

Metabolic dysfunction does not necessarily correlate with adiposity. Metabolically healthy obese individuals and insulin-resistant lean individuals represent a fundamental paradox that implicates immune cell intrinsic mechanisms in the pathogenesis of type 2 diabetes. Here, we identify myeloid Gi signaling as a previously unrecognized determinant of whole-body glucose homeostasis. Single-cell transcriptomic analysis of adipose tissue macrophages from obese mice and humans reveals marked alteration in Gnai isoform, suggesting that myeloid Gi signaling is functionally engaged during metabolic disease. Using complementary myeloid-specific rodent models of Gi inhibition (pertussis toxin) and chemogenetic Gi activation (DREADD), we demonstrate that inhibition of Gi signaling improves glucose tolerance and enhances insulin sensitivity under both regular chow and high-fat diet conditions, independent of body weight and energy expenditure. Whereas acute Gi activation in lean mice modestly enhances glucose disposal, the same intervention during diet-induced obesity markedly impairs systemic glucose homeostasis, revealing context-dependent pathway function. Mechanistically, Gi inhibition amplifies macrophage cAMP–CREB signaling to drive IL-6 production, engaging STAT3– and AMPK-dependent pathways in adipose tissue and skeletal muscle to support insulin action. Conversely, Gi activation engages a previously uncharacterized Gβγ–mTOR/AKT–JNK cascade, driving IL-1β secretion that directly impairs insulin signaling in adipocytes and myotubes. Pharmacological IL-6 receptor blockade abolishes the metabolic benefits of Gi inhibition, whereas IL-1 receptor antagonism fully rescues Gi activation–induced metabolic dysfunction, establishing these cytokines as obligate downstream effectors. This signaling architecture is conserved in human macrophages, and ATAC-seq profiling reveals chromatin remodeling at cAMP–CREB and IL-6 regulatory pathway loci, consistent with the observed transcriptional reprogramming.

Together, these findings establish myeloid Gi signaling as a weight-independent immunometabolic switch that couples opposing cytokine programs to systemic insulin sensitivity and identify this pathway as a therapeutic target in obesity-associated metabolic disease.

Graphical Abstract

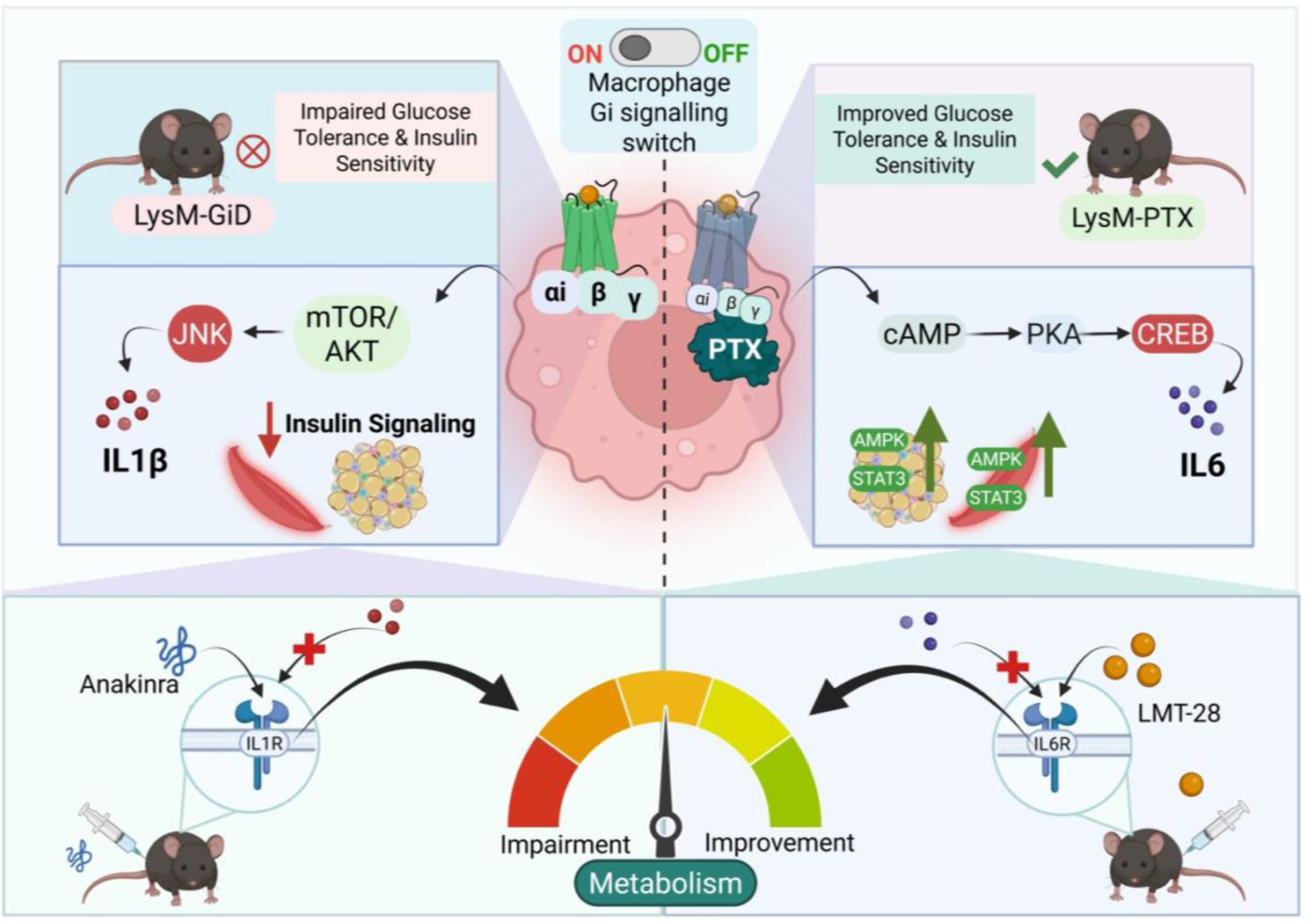

## Introduction

Chronic low-grade inflammation is now recognized as a fundamental pathogenic driver of neurodegenerative, cardiovascular, and metabolic diseases, including obesity and type 2 diabetes (T2D) [1–3]. Nevertheless, the relationship between inflammation and metabolic dysfunction is highly heterogeneous. Epidemiological studies have shown that a substantial proportion of obese individuals remain metabolically healthy, maintaining insulin sensitivity and normal glycemia despite excess adiposity, whereas some lean individuals develop insulin resistance and T2D in the absence of obesity [4–9]. These divergent phenotypes challenge the traditional paradigm that inflammation invariably drives metabolic disease and highlight critical gaps in our understanding of how inflammatory processes are regulated in metabolically active tissues.

Adipose tissue macrophages (ATMs) are central to this regulatory landscape. In healthy adipose tissue, macrophages constitute approximately 5–10% of the stromal vascular fraction and maintain tissue homeostasis through anti-inflammatory programs and metabolic buffering [9–11]. During obesity, macrophage abundance increases nearly eightfold, reaching up to 40% of adipose tissue cells [12–14]. Nevertheless, the existence of metabolically healthy obese individuals, who maintain near-normal insulin sensitivity despite comparable macrophage infiltration, demonstrates that quantitative expansion of ATMs alone does not determine metabolic outcomes [15]. Notably, macrophages have been shown to retain a pro-inflammatory transcriptional state even following sustained weight loss [16], raising the possibility that cell-intrinsic signaling programs, once established, may exert lasting and adiposity-independent control over systemic metabolism. This dissociation has focused attention on the upstream signaling networks that govern macrophage inflammatory programming across metabolic contexts. While considerable attention has been devoted to downstream cytokine effectors and polarization states in metabolic disease [17–22], the cell-intrinsic mechanisms that determine whether macrophages adopt adaptive or pathological inflammatory outputs remain poorly understood.

G protein-coupled receptors (GPCRs) are transmembrane proteins that play a central role in maintaining cellular homeostasis by sensing chemokines, metabolites, lipids, and hormones. Upon ligand binding, GPCRs couple to one of four major classes of heterotrimeric G proteins, Gs, Gq, G12/13, and Gi/o, which engage downstream pathways that control diverse aspects of immune cell behaviour. In macrophages, Gi-coupled GPCRs are key receptors that coordinate signals involved in chemotaxis, cytoskeletal changes, cytokine release, and tissue positioning [23–27]. Despite their central role in macrophage biology, the function of Gi signaling pathways has never been systematically characterized in the context of metabolic inflammation. Existing knowledge derives almost exclusively from infection models and classical immunology, where Gi governs macrophage migration toward pathogens and damaged tissue. Whether Gi-dependent signaling shapes macrophage cytokine programs and systemic insulin sensitivity under conditions of chronic nutrient excess has not been addressed. The GPCR landscape in macrophages is highly complex, with dozens of receptors converging onto a small number of G-protein pathways [28–30]. Studying individual receptors in isolation provides only a fragmented understanding of macrophage signaling. We reasoned that interrogating Gi signaling as a unifying downstream node, rather than any single receptor, would reveal fundamental principles governing how macrophages integrate metabolic and inflammatory cues during obesity. Consistent with this reasoning, analysis of publicly available single-cell transcriptomic datasets revealed upregulation of Gnai isoforms in adipose tissue macrophages from obese mice and humans, providing direct empirical rationale for mechanistic investigation of this pathway in metabolic disease.

To dissect the role of Gi signaling, we used two complementary genetic strategies that enabled bidirectional modulation of Gi pathways in a myeloid-specific manner. Gi activation was achieved using Designer Receptors Exclusively Activated by Designer Drugs (Gi-DREADDs), which allow selective pathway activation by the highly selective synthetic ligand deschloroclozapine (DCZ) [31–33]. Gi inhibition was accomplished by expressing the catalytic S1 subunit of pertussis toxin (PTX), which blocks Gi/o signaling through ADP-ribosylation [34, 35]. Together, these complementary models provide a rigorous and interpretable framework for interrogating how the activity state of myeloid Gi signaling shapes macrophage inflammatory programming and systemic glucose homeostasis.

## Results

### Macrophage Gαi expression is upregulated during metabolic stress

It is well established that Gi-coupled GPCRs regulate immune cell function across diverse disease contexts. We therefore asked whether expression of inhibitory G proteins in macrophages is altered during metabolic stress. To address this, we analyzed publicly available single-cell transcriptomic datasets from mice and humans focusing on adipose tissue-resident macrophages under metabolically healthy and obese conditions (**Figure 1A**). For the murine analysis, we integrated multiple adipose tissue single-cell RNA-seq datasets (GSE128518, GSE160729, GSE176171, GSE183288) using the scDREAMER framework to enable robust cross-study alignment and batch correction [36]. Following integration, we subsetted the immune compartment and performed cell-type annotation based on canonical marker genes to resolve distinct immune populations, including macrophages (**Figure S1A-C, S1E-H**). For the human datasets, we leveraged a previously processed and annotated single-cell atlas of adipose tissue [37], and directly interrogated GNAI gene expression within the macrophage populations without additional preprocessing or re-integration. Consistent with previous reports, macrophage abundance was markedly increased in adipose tissue from HFD fed mice and obese individuals compared to their respective controls [12] (**Figure S1C,D**). Within these expanded macrophage populations, we examined the expression of Gnai family members. Analysis of *Gnai* gene expression in macrophages revealed that *Gnai2* and *Gnai3* were markedly increased in macrophages from HFD-fed mice, whereas *Gnai1* showed minimal change (**Figure 1B**). In contrast to mice dataset, analysis of human macrophage datasets revealed elevated expression of *GNAI1* and *GNAI3* in obese individuals whereas *GNAI2* expression was higher in healthy subjects (**Figure 1C**). Feature plots further confirmed increased expression of *Gnai* genes in macrophages under metabolically stressed conditions in both species (**Figure S1I**), indicating that macrophage Gαi signaling is broadly enhanced during obesity despite species-specific differences in isoform dominance. To experimentally validate these observations, we isolated peritoneal macrophages from mice fed RC or HFD and quantified *Gnai* expression by quantitative PCR. *Gnai2* and *Gnai3* genes were significantly upregulated in macrophages from HFD-fed mice compared with controls (**Figure 1D**). Consistent with a macrophage-enriched expression pattern, analysis of bone marrow–derived macrophages (BMDMs) and peripheral blood mononuclear cells (PBMCs) revealed that *Gnai2* was the predominant isoform expressed relative to *Gnai1* and *Gnai3,* both in mouse (**Figure S1J**) and in human samples (**Figure S1K**). We next asked whether macrophage *Gnai* expression associates with metabolic parameters in vivo. Analysis of ex vivo cultured peritoneal macrophages revealed positive correlations between *Gnai1*, *Gnai2,* and *Gnai3* expression and blood glucose levels (**Figure 1E**). These observations suggest that individual Gαi isoforms may differentially reflect chronic adiposity and acute glycaemic status, pointing to distinct roles for inhibitory G protein signaling in metabolic diseases.

**Figure 1.**
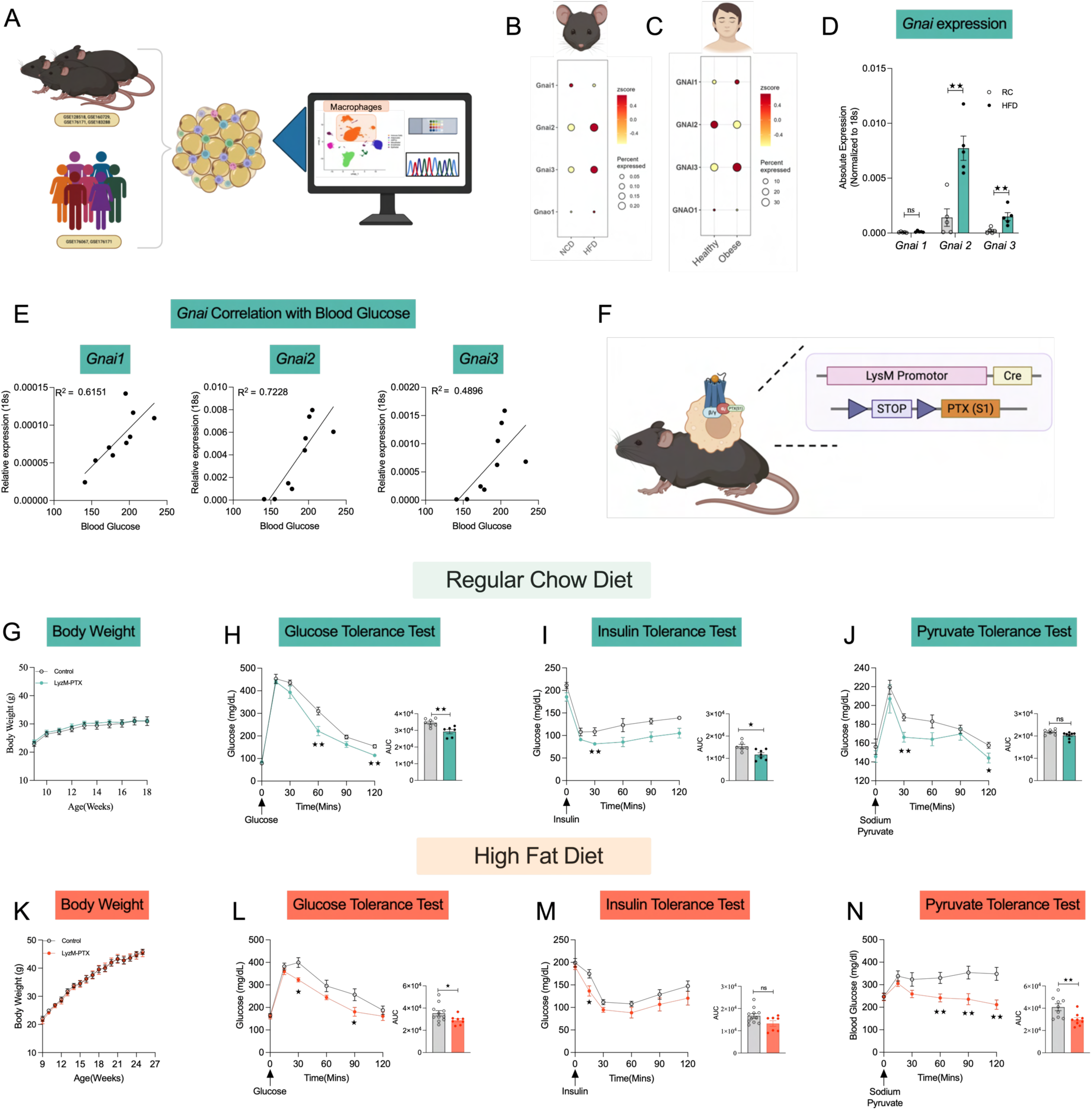
Myeloid Gαi expression increases during metabolic stress and modulates systemic glucose homeostasis. (A) Schematic overview of the experimental workflow integrating murine and human datasets to analyze Gαi signaling in macrophages under metabolic stress. (B, C) Dot plots showing the expression and proportion of immune cells expressing *Gnai* family members across immune cell populations in mice fed regular chow (RC) or high-fat diet (HFD) (B) and in lean and obese human cohorts (C). (D) qPCR analysis of *Gnai1*, *Gnai2*, and *Gnai3* expression in peritoneal macrophages isolated from RC– or HFD-fed mice. Gene expression was normalized to 18s rRNA. (E) Correlation analysis of *Gnai1*, *Gnai2*, and *Gnai3* expression with circulating blood glucose levels. (F) Schematic representation of the myeloid-specific Gαi inhibition model generated by LysM-driven expression of the pertussis toxin S1 subunit (LysM-PTX). (G, K) Body weight of control and LysM-PTX mice maintained on RC (G) or HFD (K) conditions (n = 6-11 per group). (H–J) Glucose tolerance test (GTT; glucose 2g/kg, i.p.), insulin tolerance test (ITT; insulin 0.75U/kg, i.p.), and pyruvate tolerance test (PTT; sodium pyruvate 2g/kg, i.p.) were performed in control and LysM-PTX mice maintained on RC diet. (L–N) GTT, ITT, and PTT were performed in control mice, and LysM-PTX mice were maintained on an HFD. Data are presented as mean ± s.e.m.; statistical significance was determined using an unpaired two-tailed Student’s t-test.

### Inhibition of myeloid Gi signaling improves whole-body glucose homeostasis in healthy and obese states

To directly assess the functional contribution of myeloid Gi signaling to systemic glucose metabolism, we generated mice expressing pertussis toxin subunit S1 under the control of the LysM promoter, thereby selectively inhibiting Gαi signaling in myeloid cells (LysM-PTX mice; **Figure 1F**). When maintained on RC, LysM-PTX mice displayed body weight trajectories similar to those of littermate controls (**Figure 1G**). However, LysM-PTX mice displayed significantly improved glucose tolerance(**Figure 1H**), enhanced insulin sensitivity (**Figure 1I**), and improved pyruvate tolerance(**Figure 1J**), as assessed by glucose tolerance testing (GTT), insulin tolerance testing (ITT), and pyruvate tolerance testing (PTT) respectively. We next asked whether these metabolic benefits persist under conditions of diet-induced obesity. Following prolonged HFD feeding, control and LysM-PTX mice gained weight at similar rate (**Figure 1K**). Despite comparable weight gain, LysM-PTX mice remained protected from HFD-induced metabolic dysfunction, exhibiting improved glucose tolerance (**Figure 1L**), increased insulin sensitivity (**Figure 1M**), and enhanced pyruvate tolerance **Figure 1N**), as revealed by GTT, ITT, and PTT.

To determine whether inhibition of myeloid Gi signaling alters basal metabolic or endocrine parameters, we quantified circulating glucose, insulin, glycerol, triglycerides, and non-esterified fatty acids (NEFAs) under both fed and fasted conditions. Under RC, no significant differences were observed between genotypes in basal glucose (**Figure S2A**), insulin (**Figure S2B**), glycerol (**Figure S2C**), triglyceride (**Figure S2D**), and NEFA (**Figure S2E**) levels in both fasting and fed state. Similarly, under HFD conditions, basal metabolic parameters remained comparable between control and LysM-PTX mice(**Figure S2F-H**), with the notable exception of reduced circulating NEFA levels in LysM-PTX animals (**Figure S2I**). Next, we examined energy expenditure and respiratory exchange ratio under both RC and HFD conditions, however, no difference was observed between LysM-PTX and the control group in both RC and HFD groups kept at 23^0^ and 30^0^ C (**Figure S3**). Together, these findings demonstrate that inhibition of myeloid Gi signaling improves systemic glucose homeostasis under both lean and obese conditions. Importantly, these effects occur independently of changes in body weight or basal insulin secretion, identifying myeloid Gi pathways as key modulators of whole-body glucose homeostasis during metabolic challenge.

### Diet-dependent effects of myeloid Gi signaling activation on glucose homeostasis

Given that inhibition of myeloid Gi signaling improved systemic glucose metabolism, we next asked whether acute or chronic activation of Gi signaling in myeloid cells would exert opposing effects on whole-body glucose homeostasis. To address this, we generated myeloid-specific Gi-DREADD mice (LysM-GiD) by crossing LysM-Cre mice with Rosa26-LSL-Gi-DREADD mice, enabling selective activation of Gi signaling in myeloid cells upon administration of the DREADD agonist, Deschloroclozapine (DCZ). First to confirm expression of the Gi-DREADD receptor in myeloid cells, we isolated BMDMs from LysM-GiD and control mice and assessed receptor expression at both the mRNA and protein levels. Gi-DREADD transcripts were readily detected in LysM-GiD macrophages but were absent in control cells **(Figure S4A**). Consistently, immunoblotting using the HA tag confirmed selective receptor expression in LysM-GiD BMDMs **(Figure S4B**). Analysis of metabolic tissues expression further showed that Gi-DREADD expression was present only in LysM-GiD mice **(Figure S4C**). To determine whether the receptor was functional, BMDMs were stimulated with forskolin to elevate cAMP levels and subsequently treated with DCZ to activate Gi-DREADD. DCZ stimulation significantly reduced forskolin-induced cAMP accumulation in LysM-GiD macrophages but had no effect in control cells **(Figure S4D**). This response was abolished by pretreatment with PTX, confirming that the receptor signals through Gi **(Figure S4E**). Together, these results demonstrate selective and functional expression of Gi-DREADD in myeloid cells of LysM-GiD mice.

Next, we evaluated metabolic parameters under basal conditions. LysM-GiD mice and control littermates displayed comparable body weights (**Figure S5A**) and blood glucose levels (**Figure S5B**). Acute activation of macrophage Gi signaling by DCZ administration did not alter basal glucose levels or overall glucose homeostasis (**Figure S5C**). However, DCZ-treated LysM-GiD mice displayed significantly improved glucose clearance during GTT (**Figure S5D**), without detectable changes in pyruvate (**Figure S5E**) or insulin tolerance test (**Figure S5F**). These findings indicate that, in the RC condition, acute macrophage Gi signaling can enhance peripheral glucose disposal without broadly perturbing metabolic homeostasis.

We next examined the effects of acute Gi activation in the context of diet-induced obesity. LysM-GiD and control mice were subjected to HFD feeding for 12 weeks (**Figure 2A**). Body weight gain and blood glucose were comparable between genotypes throughout the feeding period; no gross differences were observed between groups (**Figure 2B,C**). To exclude constitutive Gi-DREADD activity and off-target effects of LysM-Cre, glucose tolerance was assessed in the absence of DCZ administration. LysM-GiD and control mice performed comparably on GTT under basal conditions (**Figure 2D**), confirming ligand-dependent receptor activity. Having validated the model, we next performed an acute DCZ challenge to selectively activate myeloid Gi signaling. Following DCZ administration, LysM-GiD mice exhibited elevated blood glucose levels over time (**Figure 2E**) and a marked impairment in glucose clearance during GTT (**Figure 2F**), indicative of defective peripheral glucose disposal. Consistent with this, ITT revealed significantly reduced insulin sensitivity in LysM-GiD mice (**Figure 2G**), whereas no significant differences were observed in pyruvate tolerance testing between groups (**Figure 2H).** Acute DCZ treatment did not alter circulating insulin levels in either control or LysM-GiD mice (**Figure 2I**). In parallel, lipid profiling revealed a trend toward increased circulating NEFAs in LysM-GiD mice 15 minutes following DCZ administration (**Figure S6A**). Together, these findings indicate that acute activation of myeloid Gi signaling impairs systemic glucose homeostasis without detectable changes in circulating insulin levels.

**Figure 2.**
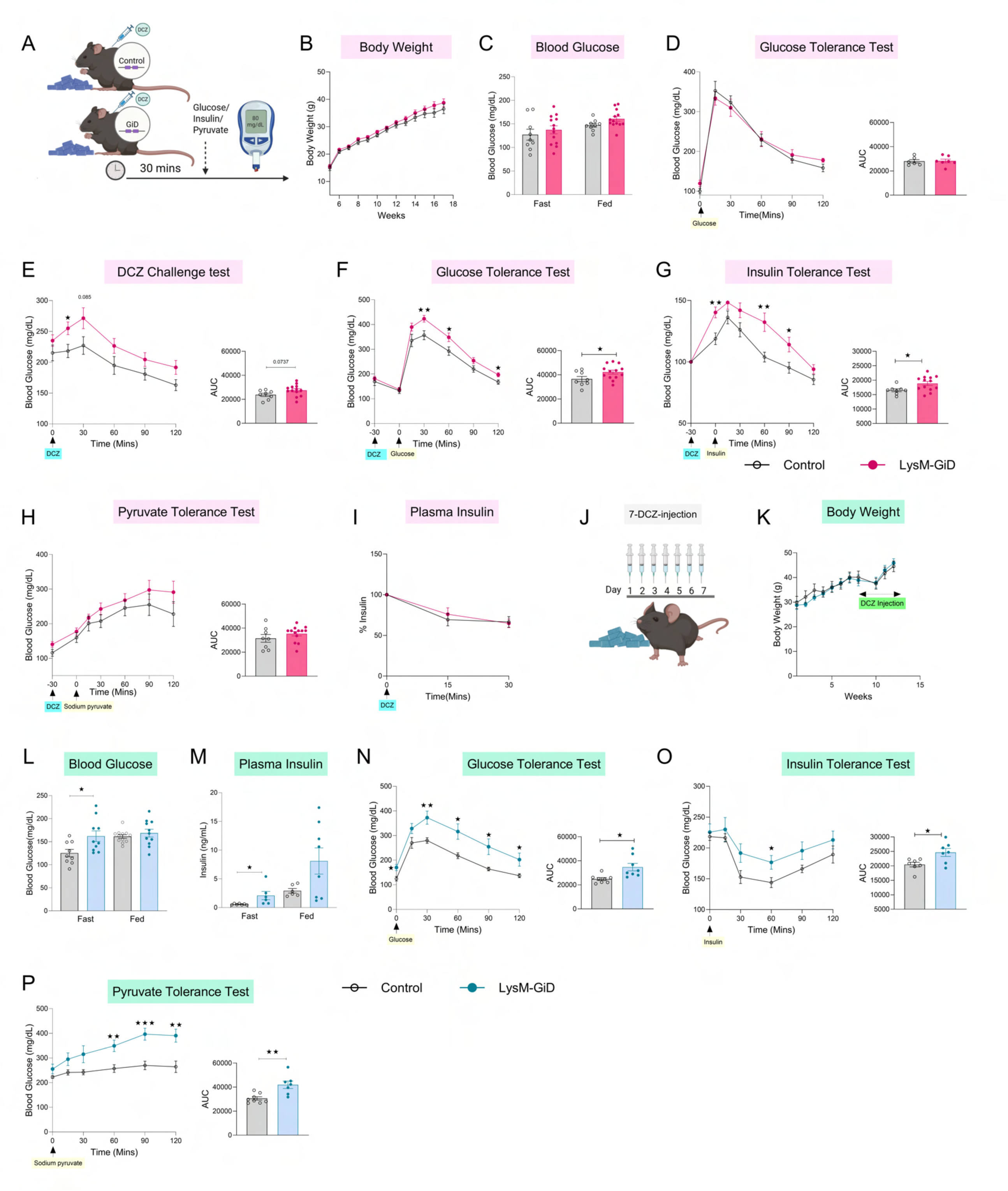
Acute and chronic activation of GiD signaling in myeloid cells regulates glucose homeostasis. (A) Schematic representation of acute GiD activation following intraperitoneal administration of DCZ (30 µg/kg) 30 min before metabolic assays. (B) Body weights of control and LysM-GiD mice (n = 8–13 per group). (C) Blood glucose is measured at the fed and fasting states. (D) Glucose tolerance test performed without GiD activation following an overnight fast (1 g glucose/kg; n = 7 per group. (E) Blood glucose levels were measured after acute DCZ injection (30 µg/kg, i.p.) following a 4-h fast (n = 8–13 per group). (F) Effect of acute DCZ treatment (30 µg/kg, i.p.) on glucose tolerance (GTT; 1 g glucose/kg body weight) after an overnight fast (n = 8–13 per group). (G) Effect of acute DCZ treatment on insulin tolerance (ITT; 1.5 U insulin/kg body weight) after a 6-h fast (n = 8–13 per group). (H) Effect of acute DCZ treatment on pyruvate tolerance (PTT; 1 g sodium pyruvate/kg body weight) after a 6-h fast (n = 8–13 per group). (I) Plasma insulin levels were measured after acute DCZ treatment following a 6-h fast (n = 8 per group). (J) Schematic representation of chronic GiD activation achieved by daily intraperitoneal injections of DCZ (30 µg/kg) for 7 days. (K) Body weights of control and LysM-GiD mice following chronic DCZ treatment (n = 11–12 per group). (L) Blood glucose levels in fed and fasted states following chronic GiD activation (n = 9–12 per group). (M) Plasma insulin levels in fed and fasted states following chronic GiD activation (n = 6–8 per group). (N) Effect of chronic GiD activation on glucose tolerance (GTT; 1 g glucose/kg body weight, i.p.) after an overnight fast (n = 8 per group). (O) Effect of chronic GiD activation on insulin tolerance (ITT; 1.5 U insulin/kg body weight, i.p.) after a 6-h fast (n = 7–8 per group). (P) Effect of chronic GiD activation on pyruvate tolerance (PTT; 1 g sodium pyruvate/kg body weight, i.p.) after a 6-h fast (n = 8 per group). Data are presented as mean ± s.e.m., C-M: two-tailed Student’s t-test.

We next evaluated the consequences of sustained Gi activation. To this end, LysM-GiD and control mice were maintained on an HFD for 12 weeks, followed by daily intraperitoneal administration of DCZ (30 μg/kg) to chronically activate Gi signaling (**Figure 2J**). Chronic Gi activation did not alter body weight (**Figure 2K**), but it resulted in persistent metabolic dysfunction as LysM-GiD mice had elevated fasting blood glucose levels (**Figure 2L**) and increased circulating insulin concentrations (**Figure 2M**), indicating progressive disruption of glucose homeostasis. LysM-GiD mice also displayed impaired glucose clearance during GTT (**Figure 2N**) and reduced insulin sensitivity during ITT (**Figure 2O**). In contrast to the acute setting, chronic Gi activation also led to impaired pyruvate tolerance (**Figure 2P**). No significant differences in fasting and fed plasma NEFA (**Figure S6B**) or glycerol (**Figure S6C**) levels were observed between groups in chronic DCZ treatment. These results show that chronic macrophage Gi signaling induces sustained impairments in glucose metabolism, insulin action, and pancreatic islet function, without altering adipocyte morphology in iWAT and eWAT (**Figure S6D**). Next, we asked whether chronic activation of myeloid Gi signaling alters energy homeostasis. To address this, we measured total energy expenditure and respiratory exchange ratio during both light and dark cycles. No significant differences were observed between LysM-GiD and control mice under these conditions (**Figure S7**).

### Acute and chronic activation of myeloid Gi-signaling induces an IL-1–driven inflammatory program that impairs insulin signaling in target tissues

Given that both acute and chronic activation of myeloid Gi signaling impaired glucose homeostasis, we next asked whether myeloid Gi activation alters systemic inflammatory tone, thereby contributing to metabolic dysfunction. To test this, we performed multiplex cytokine profiling of plasma from LysM-GiD and control mice following acute DCZ treatment (1 hour). Notably, LysM-GiD mice exhibited significantly elevated levels of pro-inflammatory cytokines, including IL-1β, CXCL1, and MCP-3 (**Figure 3A**), however, no significant difference was observed in MIP-1α, MIP-2α, CCL-11, CXCL-10 (**Figure S8A-D**).

**Figure 3.**
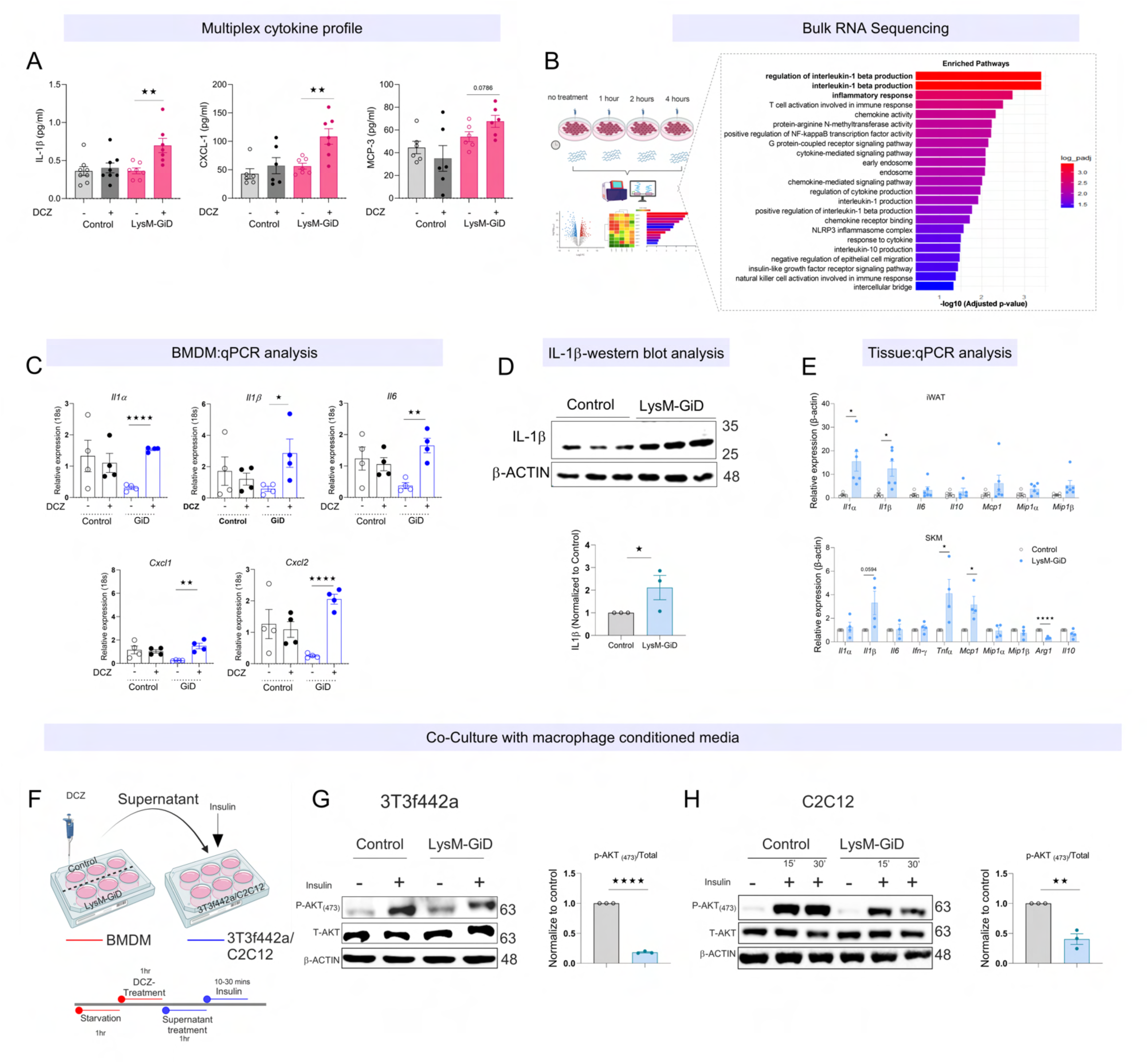
Acute and chronic activation of macrophage Gi-signaling induces an IL-1–driven inflammatory program that impairs insulin signaling in target tissues. (A) Multiplex cytokine analysis of plasma collected from control and LysM-GiD mice 30 min after vehicle or DCZ injection (30 µg/kg, i.p.). (B) Bulk RNA sequencing of bone BMDMs isolated from LysM-GiD mice and treated with DCZ (100 nM) for 1, 2, or 4 h (n = 3 per time point). Untreated BMDMs served as controls. Pathway enrichment analysis for the 1 h DCZ treatment showing significantly upregulated and downregulated pathways. (C) qPCR analysis of inflammatory cytokine gene expression in BMDMs treated with vehicle or DCZ (100 nM) for 1 h from control and LysM-GiD mice (n = 4 per group). (D) Representative Western blot and quantification of IL-1β protein levels in BMDMs treated with DCZ (100 nM) for 1 h (n = 6 per group). (E) qPCR analysis of cytokine expression in inguinal white adipose tissue (iWAT; n = 5–6 per group) and skeletal muscle (SKM; n = 4 per group) from control and LysM-GiD mice fed a high-fat diet (HFD) following 7 days of chronic DCZ treatment (30 µg/kg, i.p.). (F) Schematic representation of the macrophage–adipocyte/myocyte co-culture experimental design. BMDMs were treated with DCZ (30 nM) for 1 h, and the conditioned media were applied to differentiated adipocytes (3T3-F442a) or myotubes (C2C12) for 1 h, followed by insulin stimulation (10 min for 3T3-F442a and 15 and 30 min for C2C12). (G) Western blot analysis and quantification of insulin signaling (p-AKT/total AKT) in 3T3-F442a adipocytes treated with conditioned media from control or LysM-GiD BMDMs (n = 3 per group). (H) Western blot analysis and quantification of insulin signaling (p-AKT/total AKT) in C2C12 myotubes pre-incubated with conditioned media from control or LysM-GiD BMDMs, with or without insulin stimulation (n = 3 per group). Data are presented as mean ± s.e.m., A, C-D, E, G-H: two-tailed Student’s t-test.

To delineate the molecular consequences of Gi signaling in macrophages, we performed bulk RNA sequencing on BMDMs from LysM-GiD mice following DCZ stimulation at defined time points (0 h, untreated; 1 h; 2 h; and 4 h) (**Figure 3B**). Differential expression analysis revealed robust transcriptional changes at 1 and 2 hours after Gi activation, whereas no significant differences were detected at 4 hours (**Figure S9A-C, Figure S10A,B, Figure S11A**). Gene set enrichment analysis identified strong induction of inflammatory pathways, with significant enrichment of gene sets associated with regulation of interleukin-1β production, interleukin-1β production, and inflammatory responses (**Figure 3B**). We validated these findings by quantitative PCR using a targeted panel of inflammatory and anti-inflammatory genes (**Figure 3C** and **Figure S10 B-F**). Consistent with the RNA-seq data, *Il-1α*, *Il-1β*, *Il-6*, *Cxcl1*, and *Cxcl2* were rapidly upregulated following 1 hour of DCZ treatment in LysM-GiD BMDMs. To determine whether these transcriptional changes translated to protein-level responses in cells, we assessed IL-1β expression in DCZ-treated LysM-GiD and control BMDMs. As expected, Gi activation resulted in a marked increase in IL-1β protein abundance (**Figure 3D**).

To determine whether myeloid Gi activation in vivo recapitulates the pro-inflammatory response observed in macrophages in vitro, we acutely activated Gi signaling by administering DCZ and harvested insulin-sensitive tissues two hours later, including adipose depots and skeletal muscle. Acute Gi activation led to a selective increase in inflammatory gene expression across tissues. In particular, *Il-1α* and *Il-1β* expression was significantly elevated in iWAT, whereas skeletal muscle exhibited marked upregulation of *Tnf-α*, *Mcp1*, and *Il-1β* (**Figure 3E**).

To test whether macrophage-derived factors are sufficient to impair metabolic cell function, we performed indirect co-culture experiments using conditioned media from DCZ-treated LysM-GiD or control macrophages isolated from HFD mice (1-hour stimulation). Conditioned media were transferred onto differentiated 3T3-F442A adipocytes and C2C12 myotubes (**Figure 3F**). Exposure to supernatants from Gi-activated macrophages resulted in impaired insulin signaling in both adipocytes (**Figure 3G**) and myotubes (**Figure 3H**), as evidenced by reduced insulin responsiveness compared with controls. Together, these findings indicate that acute activation of Gi signaling in macrophages promotes the release of inflammatory mediators that act in a paracrine manner to impair insulin signaling and glucose metabolism in adipose tissue and skeletal muscle.

### Gi signaling drives IL-1β production in macrophages through a Gβγ–mTOR–JNK axis

To identify the intracellular mechanisms coupling Gi activation to inflammatory cytokine production, we interrogated candidate downstream signaling pathways in macrophages. BMDMs from LysM-GiD and control mice were stimulated with DCZ (100nM) for 30 minutes, followed by analysis of key signaling intermediates. Activation of Gi signaling increased phosphorylation of AKT at both Thr308 and Ser473, together with enhanced phosphorylation of S6K, indicating activation of mTOR signaling (**Figure 4A**). However, no phosphorylation was observed in P85 subunit of P13K **(Figure S12A).** Given that mTOR has been reported to activate JNK which regulates the production of inflammatory cytokines, including IL-1β [38, 39], we next examined whether JNK is activated downstream of Gi signaling. As expected, we observed a marked increase in phospho-JNK levels, placing JNK downstream of Gi-dependent signaling in macrophages (**Figure 4A**).

**Figure 4.**
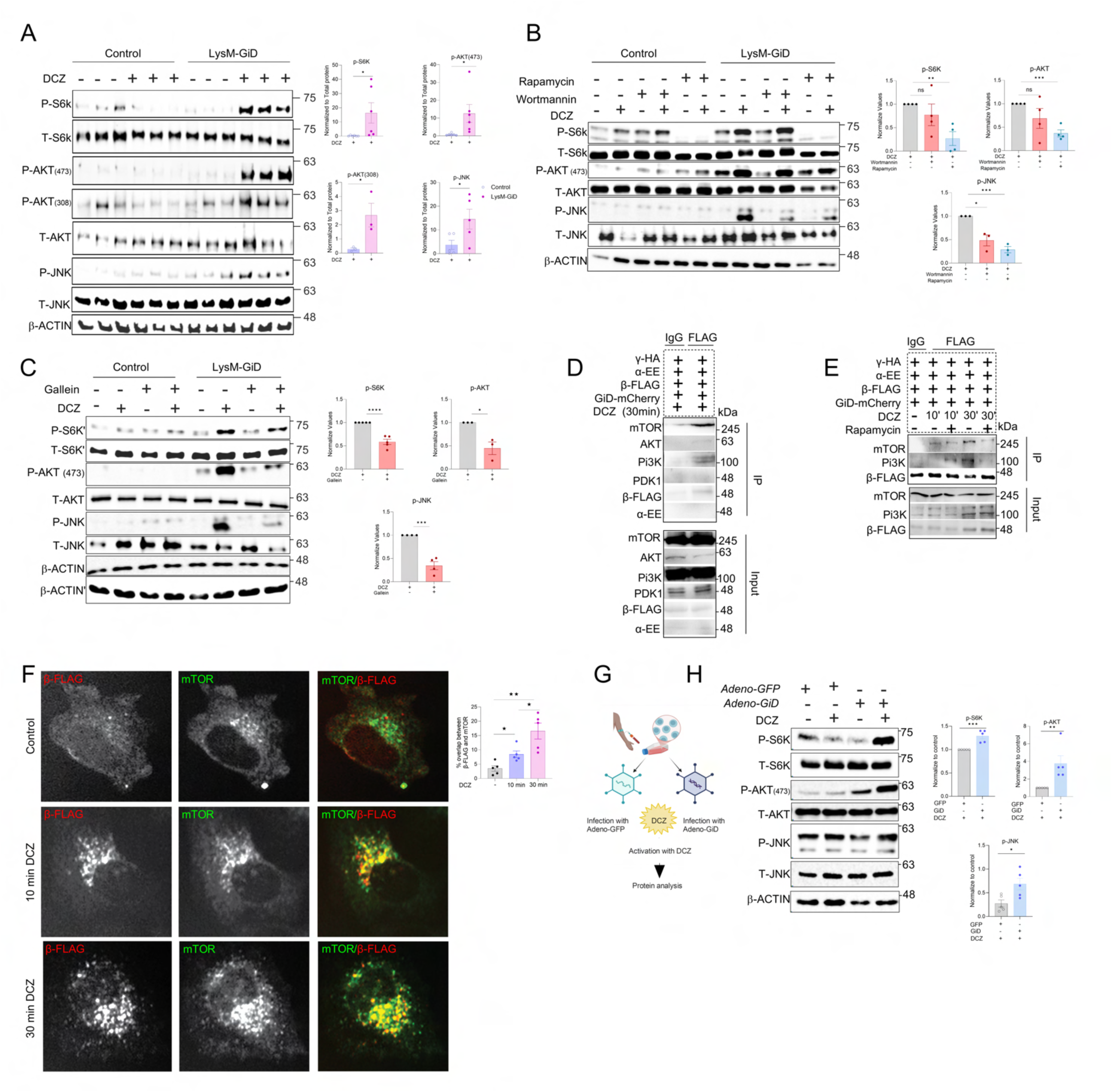
GiD signaling activates the AKT–mTOR pathway in macrophages. (A) Immunoblot analysis of signaling downstream of GiD activation in control and LysM-GiD BMDMs following DCZ stimulation (30 min). Phosphorylation of S6K, AKT (Ser473 and Thr308), and JNK was assessed by western blot; total proteins and β-actin served as loading controls. (B) Pharmacological inhibition of PI3K–mTOR signaling in control and LysM-GiD BMDMs. Cells were treated with wortmannin (100 nM, 1 h) or rapamycin (100 nM, 12 h) before DCZ stimulation (30 min). Quantification values were normalized to total protein levels and expressed relative to the GiD + DCZ condition, which was set to 1. Fold changes upon inhibitor treatment (wortmannin or rapamycin) were calculated relative to this reference (C) Effect of Gβγ inhibition on GiD-mediated signaling. Control and LysM-GiD BMDMs were treated with the Gβγ inhibitor gallein (100 µM, 1 h) before DCZ stimulation and analyzed by western blot. Quantification values were normalized to total protein levels and expressed relative to the GiD + DCZ condition, which was set to 1. Fold changes upon Gallein treatment were calculated relative to this reference (D) Co-immunoprecipitation (Co-IP) analysis demonstrated interaction of the β subunit with mTOR pathway components, including PI3K (p110), PDK1, AKT, and mTOR, as well as the Gα subunit. FLAG antibody was used to immunoprecipitate the β subunit complex, with IgG as a control. (E) Co-IP analysis of β subunit interactions with downstream signaling proteins in the presence of rapamycin following DCZ stimulation (10 and 30 min). (F, G) High-content microscopy showing colocalization of the β subunit with mTOR following GiD activation and corresponding quantification. (H) Experimental schematic of human PBMCs transduced with adenovirus-GFP (control) or adenovirus-GiD, followed by DCZ stimulation and protein isolation for signaling analysis. (I) Immunoblot analysis of PBMCs expressing adeno-GFP or adeno-GiD following DCZ stimulation. Quantification of phosphorylated S6K, AKT (Ser473), and JNK is shown. Data are presented as mean ± s.e.m. Statistical significance was determined using a two-tailed Student’s t-test.

To functionally validate this signaling cascade, we employed a series of pharmacological inhibitors targeting distinct nodes of the pathway. Inhibition of Gi signaling with pertussis toxin (PTX, 100 nM) **(Figure S12B)** and blockade of Gβγ using gallein (100 μM) (**Figure 4C, Figure S12D**) both resulted in a marked reduction in phosphorylation of S6K, AKT, and JNK, confirming the dependence of downstream signaling on Gi and Gβγ activity. To delineate the contribution of downstream signaling nodes, we next inhibited mTOR using rapamycin (100 nM) and PI3K using wortmannin (100 nM). PI3K inhibition did not significantly alter S6K or AKT phosphorylation; however, JNK activation was substantially reduced. In contrast, mTOR inhibition attenuated phosphorylation of AKT and JNK, but not PI3K (**Figure 4B, Figure S12C**). Together, these results define a signaling hierarchy in which Gi activation engages the mTOR/ AKT–JNK axis to promote inflammatory responses in macrophages.

To define the molecular basis of this signaling architecture, we examined whether Gi activation promotes stimulus-dependent assembly of a Gβγ-associated signaling complex. To delineate the biochemical composition of the Gi-activated signaling complex and validate key effector intermediates, we performed co-immunoprecipitation (Co-IP) assays in HEK293T cells transiently co-expressing HA-GiD, Gαi-EE, Gβ-FLAG, and Gγ-HA. Building on prior evidence of a physical Gβγ–PI3K association [40], Gβ was immunoprecipitated using anti-FLAG antibody following 30-minute DCZ stimulation, and co-precipitating proteins were detected by immunoblotting. Gβ co-precipitated with PI3K p110, AKT, and mTOR (**Figure 4D**), indicating that Gi activation recruits these effectors into a common Gβ-associated complex. Consistent with the specificity of this interaction, no co-precipitation was observed with Gαi-EE. To define the pathway dependency of these associations, Co-IP was performed in the presence of wortmannin. Wortmannin did not disrupt Gβ co-precipitation with AKT or mTOR (**Figure S12E**), indicating that AKT and mTOR are recruited to Gβγ independently of PI3K catalytic activity. To establish the temporal dynamics of complex formation, time-course Co-IP experiments were performed at defined intervals following DCZ stimulation, in the presence or absence of rapamycin. At 10 minutes, Gβ co-precipitation with mTOR was readily detectable, whereas PI3K association was absent; rapamycin pretreatment abolished these early interactions (**Figure 4E**). By 30 minutes, mTOR and PI3K co-precipitation with Gβ was no longer detected (**Figure 4E**). High-content imaging at both time points independently confirmed dynamic Gβ–mTOR co-localization, with peak co-localization at 30 minutes (**Figure 4F**). Taken together, these biochemical and imaging data establish that Gi activation drives rapid, stimulus-dependent recruitment of mTOR to Gβγ in a temporally ordered and rapamycin-sensitive manner. Providing direct molecular evidence for the Gβγ–mTOR–JNK axis as the effector conduit through which macrophage Gi signaling promotes inflammatory cytokine production.

Finally, to determine whether this signaling axis is conserved in human macrophages, we isolated PBMC-derived macrophages and transduced them with adenoviruses encoding Gi-DREADD or GFP control (**Figure 4G**). DCZ stimulation elicited comparable activation of the mTOR/AKT–JNK pathway in human macrophages (**Figure 4H**), indicating that this Gi-dependent inflammatory signaling cascade is conserved across species.

### Inhibition of IL-1β signaling reverses metabolic defects induced by myeloid Gi activation

To determine whether Gi activation–induced metabolic impairment is mediated through IL-1β signaling, we pharmacologically blocked IL-1 receptor activity using the antagonist anakinra [41]. In the acute setting, both LysM-GiD and control mice received two doses of anakinra, administered 12–14 hours and again 2 hours prior to metabolic testing (**Figure 5A**). Under these conditions, DCZ challenge testing revealed no differences in blood glucose excursions between LysM-GiD and control cohorts (**Figure 5B**). Consistently, glucose tolerance (**Figure 5C**) and insulin tolerance (**Figure 5D**) tests showed that anakinra treatment completely abolished the glucose intolerance and insulin resistance observed in DCZ-treated LysM-GiD mice, indicating that inhibition of IL-1β signaling normalizes systemic glucose homeostasis. We next assessed the contribution of IL-1β signaling during sustained activation of myeloid Gi signaling. LysM-GiD mice were administered DCZ every three days, while anakinra was delivered on days −6, −3, −1, and −2 hrs relative to metabolic testing (**Figure 5E**). Strikingly, anakinra-treated LysM-GiD mice showed marked improvements in glucose tolerance (**Figure 5F**) and insulin sensitivity (**Figure 5G**) compared with vehicle-treated LysM-GiD controls. Together, these findings demonstrate that IL-1β signaling is required for the metabolic dysfunction induced by both acute and chronic activation of myeloid Gi signaling in vivo, establishing IL-1β as a key downstream mediator linking macrophage Gi activation to impaired systemic glucose homeostasis.

**Figure 5.**
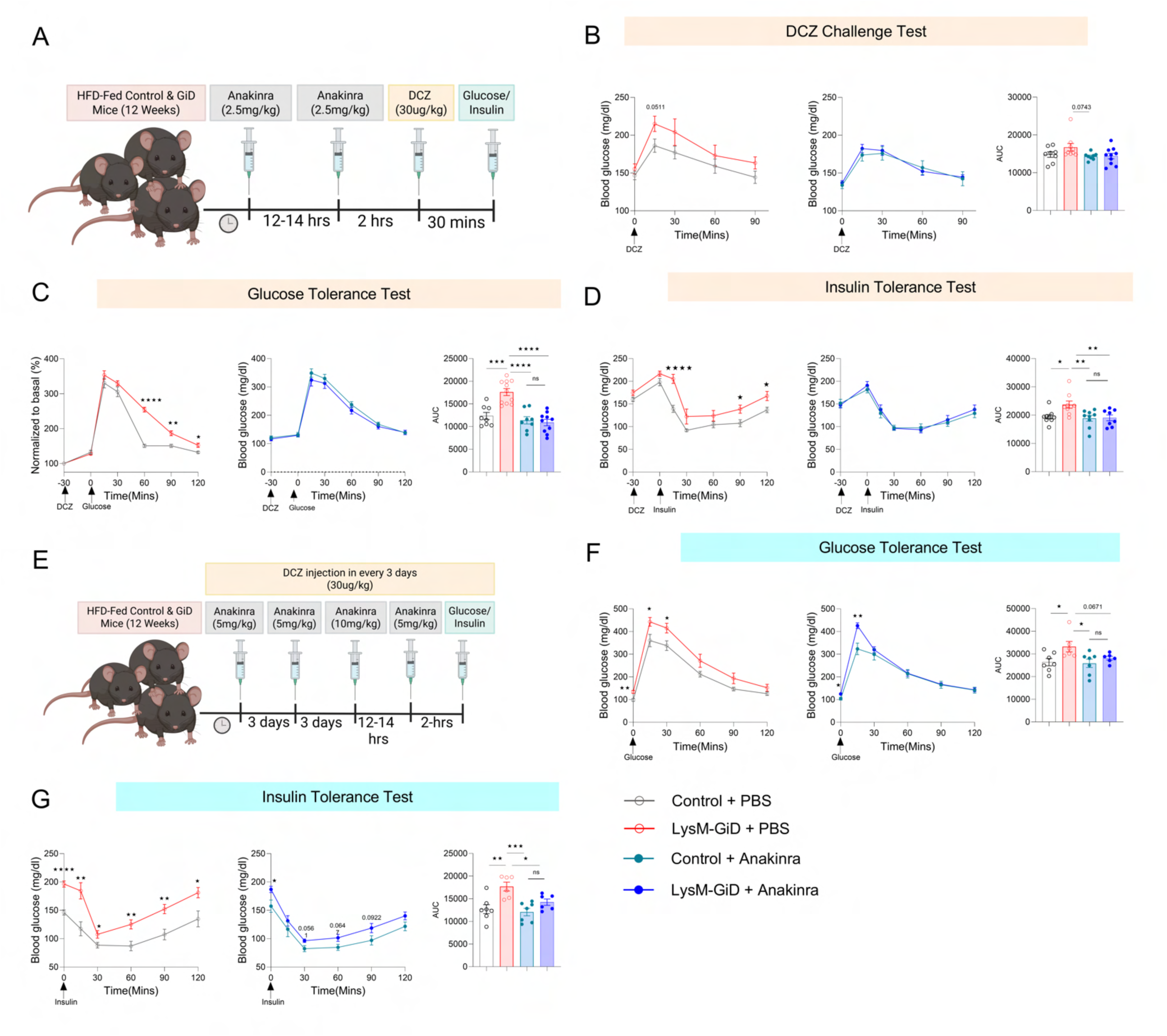
IL-1R inhibition modulates the metabolic effects of acute and chronic Gi-signaling activation in LysM-GiD mice. (A) Schematic representation of acute IL-1 receptor (IL-1R) inhibition using anakinra (2.5 mg/kg, i.p.; two doses) in control and LysM-GiD mice, combined with acute activation of Gi signaling by deschloroclozapine (DCZ; 30 µg/kg, i.p.). (B) DCZ challenge test performed in vehicle-and anakinra-treated control and LysM-GiD mice (n = 8–10 per group). (C) Effect of IL-1R inhibition on glucose tolerance during a glucose tolerance test (GTT; glucose 1 g/kg, i.p.) following acute GiD activation with DCZ (30 µg/kg, i.p.). (D) Effect of IL-1R inhibition on insulin sensitivity assessed by an insulin tolerance test (ITT; insulin 1.5 U/kg, i.p.) following acute GiD activation. (E) Schematic representation of the chronic IL-1R inhibition protocol, in which mice received four doses of anakinra (5, 5, 10, and 5 mg/kg, i.p.) along with DCZ injections (30 µg/kg, i.p.) every three days, followed by metabolic assessments. (F) Effect of anakinra on glucose tolerance following chronic activation of Gi signaling, assessed by GTT after overnight fasting (glucose 1 g/kg, i.p.) (n = 6–7 per group). (G) Effect of anakinra on insulin sensitivity following chronic activation of Gi signaling, assessed by ITT after 6 h of fasting (insulin 1.5 U/kg, i.p.) (n = 6–7 per group). Data are presented as mean ± s.e.m., B-D, F-G: two-tailed Student’s t-test.

### Myeloid Gi inhibition enhances cAMP–CREB–IL-6 signaling to improve systemic glucose homeostasis

Given that activation of myeloid Gi signaling impaired systemic glucose homeostasis through IL-1β, whereas inhibition of Gi signaling improved glucose metabolism, we next sought to define the molecular mechanisms underlying this protective phenotype. As Gαi proteins are known to suppress adenylyl cyclase activity, we first examined whether blockade of myeloid Gi signaling alters intracellular cAMP responses. To this end, we isolated peritoneal macrophages and BMDM from LysM-PTX and control mice and stimulated them with forskolin (FRS), a direct activator of adenylyl cyclase. cAMP accumulation was significantly elevated in peritoneal macrophages from LysM-PTX mice compared with controls (**Figure 6A**), confirming effective inhibition of Gαi-mediated signaling.

**Figure 6.**
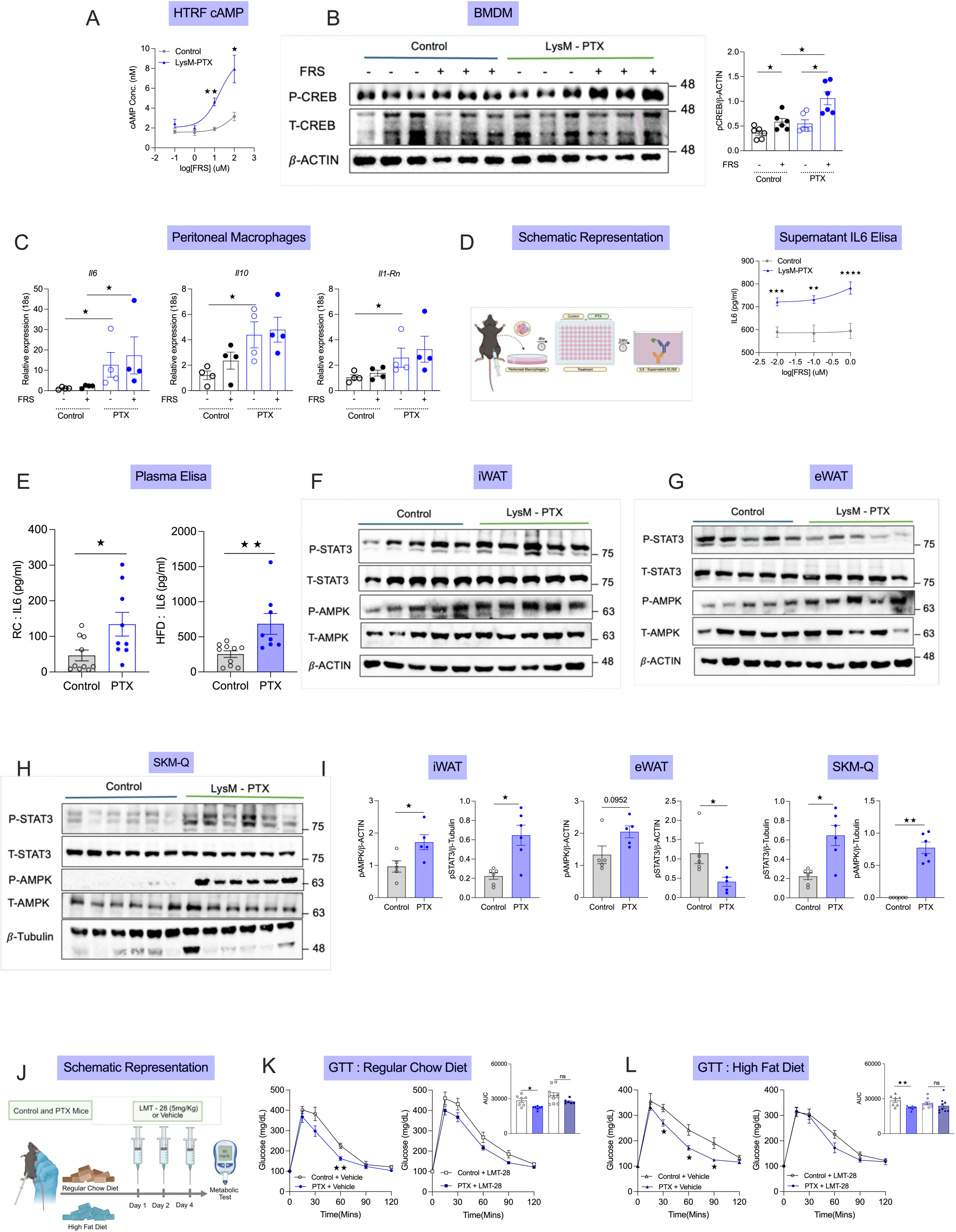
Gi signaling regulates macrophage inflammatory responses and systemic metabolic regulation. (A) HTRF cAMP assay measuring intracellular cAMP levels in control and LysM-PTX macrophages following stimulation with increasing concentrations of forskolin(FRS), demonstrating functional inhibition of Gi signaling. (B) Immunoblot analysis of CREB phosphorylation in bone marrow-derived macrophages treated with FRS (10uM) in control and LysM-PTX conditions. Quantification of phosphorylated CREB normalized to total CREB is shown. (C) Relative expression of *Il6*, *Il10*, and *Il1rn* in macrophages under basal and forskolin-stimulated conditions, isolated from control and LysM-PTX mice following FRS stimulation. (D) Experimental schematic illustrating macrophage stimulation and subsequent analysis of IL6 secretion upon FRS stimulation in macrophage culture supernatants. (E) ELISA quantification of plasma IL-6 concentration in control and LysM-PTX mice under regular chow (RC) and high-fat diet (HFD) conditions. (F–H) Immunoblot analysis of STAT3 and AMPK signaling pathways in inguinal white adipose tissue (iWAT), epididymal white adipose tissue (eWAT), and glycolytic skeletal muscle (SKM-Q) from control and LysM-PTX mice. (I) Quantification of pSTAT3 and pAMPK signaling across metabolic tissues. (J) Experimental schematic depicting pharmacological inhibition of IL-6 signaling using LMT-28 in control and LysM-PTX mice during metabolic testing. (K, L) Glucose tolerance tests were performed in control and LysM-PTX mice treated with vehicle or LMT-28 under regular chow (K) or high-fat diet (L) conditions (n = 6–10 per group). Insets show AUC quantification. Data are presented as mean ± s.e.m. Statistical significance was determined using a two-tailed Student’s t-test.

We next asked whether increased cAMP signaling translated into enhanced CREB activation [42]. Basal and FRS-stimulated phosphorylation of CREB was markedly increased in LysM-PTX BMDMs relative to controls, as assessed by immunoblotting (**Figure 6B**). These findings indicate sustained activation of the cAMP–CREB axis following macrophage Gi inhibition. We next examined whether this signaling shift altered cytokine expression. Under both basal conditions and FRS stimulation, LysM-PTX macrophages exhibited increased expression of *Il6 compared to controls. Concurrently the* anti-inflammatory genes*, Il10* and *Il1-Rn* genes, were significantly upregulated at steady state in LysM-PTX macrophages relative to controls (**Figure 6C).** In addition expression of *Arg1*, *Klf4*, and *Atf4* was elevated, consistent with a regulatory and stress-adaptive transcriptional program. In contrast, canonical pro-inflammatory markers such as *Il1β* and *Nfkb* showed no marked induction, and *Il4* expression remained unchanged (**Figure S13A-F**). Futhermore, previous studies have implicated CREB in the regulation of Il6 transcription toward an anti-inflammatory program [43]. Consistent with this, FRS treatment induced a dose-dependent increase in IL-6 secretion from LysM-PTX macrophages, compared to control macrophages (**Figure 6D**). These data indicate that inhibition of myeloid Gi signaling amplifies cAMP–CREB–dependent IL-6 production at the cellular level. To determine whether this enhanced IL-6 production is reflected systemically, we next measured plasma IL-6 concentrations in LysM-PTX and control mice under both RC and HFD conditions. Notably, IL-6 levels were significantly elevated in LysM-PTX mice in both dietary states (**Figure 6E**). In contrast, circulating levels of IL-1β and TNF-α (**Figure S13G**) were not significantly altered in HFD condition, indicating that the myeloid Gi inhibition does not induce a broad pro-inflammatory cytokine response but instead promotes a selective CREB mediated IL-6 dominant programme.

Given that IL-6 signals through STAT3 and can engage insulin-independent, AMPK-dependent metabolic pathways [44–46], we next assessed downstream signaling in insulin-sensitive tissues. In inguinal white adipose tissue, LysM-PTX mice exhibited significant increases in both STAT3 and AMPK phosphorylation, consistent with a metabolically active, insulin-sensitive adipose phenotype (**Figure 6F,I**). In epididymal white adipose tissue, STAT3 phosphorylation was significantly reduced, whereas AMPK activation showed a trend toward increase (**Figure 6G,I**), in agreement with prior studies, studies associating reduced STAT3 signaling in visceral adipose tissue with improved metabolic outcomes [47, 48]. In skeletal muscle, both STAT3 and AMPK phosphorylation were robustly and significantly elevated (**Figure 6H,I**), confirms the enhanced IL-6-driven metabolic signaling across peripheral tissues. Finally, to establish the functional relevance of IL-6 in mediating the metabolic benefits of Gi inhibition, IL-6 receptor signaling was pharmacologically blocked using LMT-28 (**Figure 6J**). Under both RC and HFD conditions, IL-6R inhibition abolished the improved glucose tolerance observed in LysM-PTX mice, restoring glycaemic responses toward control levels (**Figure 6K, L**).

### Gi inhibition remodels macrophage chromatin to promote cAMP–CREB–IL-6 signaling programs

Finally, to determine whether chronic myeloid Gi inhibition is associated with epigenomic remodeling, we assessed chromatin accessibility using ATAC-seq in peritoneal macrophages isolated from LysM-PTX and control mice kept on HFD. Gi inhibition resulted in a pronounced shift in chromatin accessibility, characterized by a strong bias toward chromatin opening. In total, 1,214 regions exhibited increased accessibility, whereas only 57 regions showed reduced accessibility (**Figure S14A**), indicating that loss of Gi signaling predominantly promotes chromatin activation rather than repression. We next asked whether these accessibility changes were spatially organized across the genome. Strikingly, gained regions were overwhelmingly enriched at promoter-proximal sites, with ∼92% mapping within ±3 kb of transcription start sites (TSS), whereas only a minor fraction localized to distal or enhancer regions (**Figure S14B**). To explore the potential functional relevance of these regions, we performed pathway enrichment analysis of genes associated with regions showing increased accessibility. This analysis identified enrichment of pathways related to cAMP–CREB signaling, IL-6/STAT3 signaling, and metabolic processes, including AMPK-associated pathways (**Figure S14C**). In parallel, motif analysis revealed enrichment of binding sites for CREB/ATF, NF-κB, and KLF family transcription factors within regions of increased accessibility (**Figure S14D**). Consistent with these observations, clustering of chromatin accessibility across representative loci revealed a clear separation between WT and PTX macrophages, with PTX samples exhibiting higher accessibility across multiple gene sets (**Figure 7A**). Functional annotation of these regions indicated enrichment of processes related to cytokine signaling and transcriptional regulation (**Figure 7D**). At the gene level, increased accessibility was observed at loci associated with cAMP–CREB signaling (Crebbp, Crtc1, Atf2, Atf4, Prkaca, Creb1), IL-6/STAT3 signaling (Stat3, Jak2, Il6st), metabolic regulation (Nampt, Stk11, Prkab2), and anti-inflammatory responses (Relb, Nfkbia, Klf2). Visualization of representative loci confirmed increased promoter accessibility at genes associated with the cAMP–CREB axis, including Crebbp, Crtc1, Atf2, and Prkaca (**Figure 7B**, **S14E**). Increased accessibility was also observed at key components of the IL-6 signaling pathway, including Il6ra, Jak2, and Stat3 (**Figure 7C**). In addition, genes associated with regulatory and anti-inflammatory programs, including Klf2, Nfkbia, and Relb, showed similar patterns of increased promoter accessibility (**Figure S14F**). Together, ATAC-seq profiling myeloid Gi inhibition drives coordinated transcriptional reprogramming through convergent cAMP–CREB and IL-6–STAT3 regulatory axes to establish the insulin-sensitizing macrophage phenotype.

**Figure 7.**
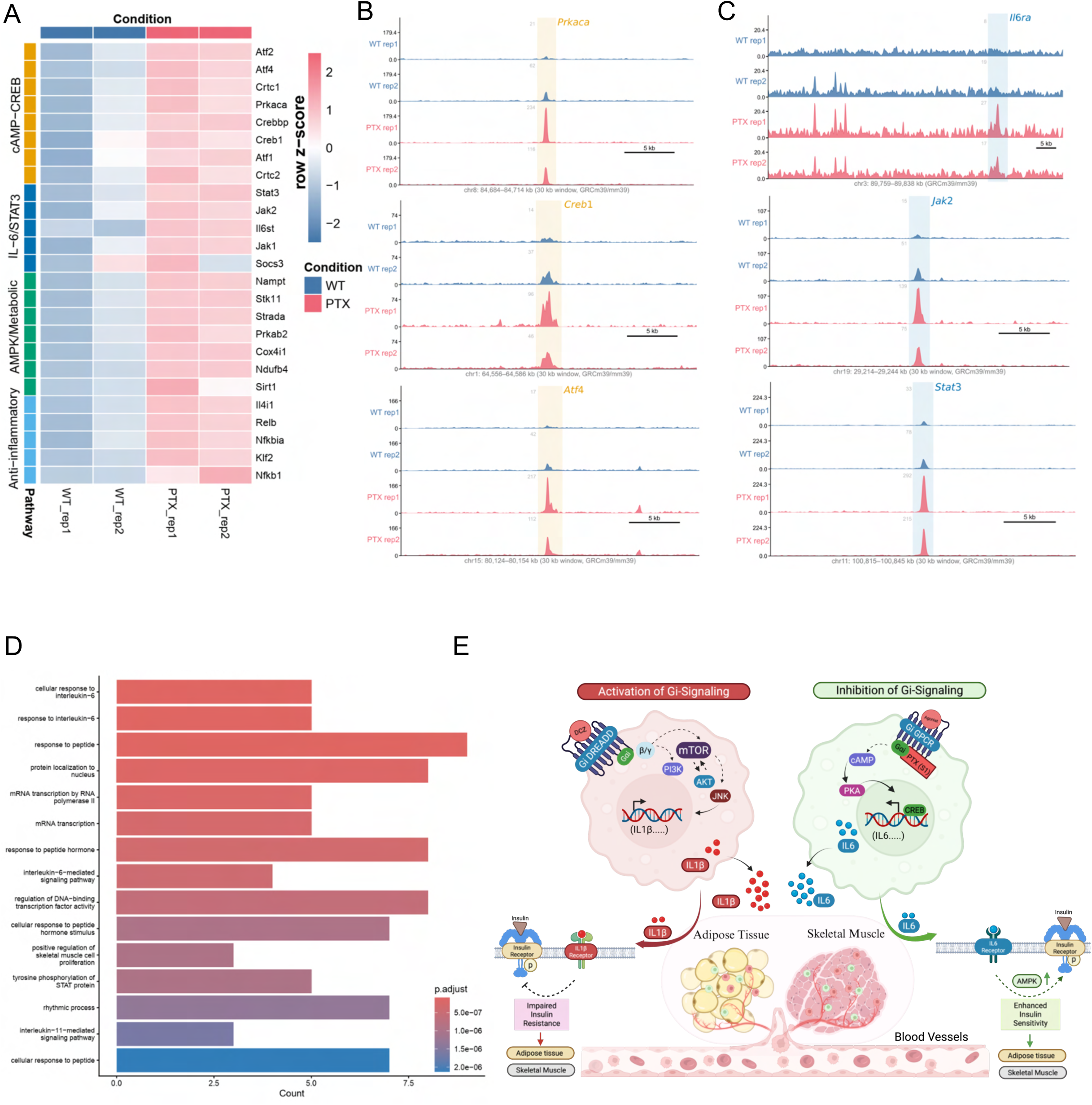
Gi signaling inhibition induces chromatin remodeling and CREB–IL-6 associated transcriptional programs in macrophages. (A) Heatmap of chromatin accessibility across selected loci associated with cAMP–CREB signaling, IL-6/STAT3 signaling, metabolic regulation, and anti-inflammatory programs in WT and LysM-PTX macrophages, showing relative accessibility differences between groups. (B) Genome browser tracks showing chromatin accessibility at representative loci associated with the cAMP–CREB signaling axis (Creb1, Atf2, Prkaca), illustrating increased accessibility in LysM-PTX macrophages compared to WT controls. (C) Genome browser tracks showing chromatin accessibility at representative loci associated with IL-6 signaling (Il6ra, Jak2, Stat3), indicating increased accessibility in LysM-PTX macrophages. (D) Functional enrichment analysis of genes associated with regions showing increased accessibility, highlighting enrichment of pathways related to cytokine signaling and transcriptional regulation. Data represent pooled samples from N = 3 mice each, with two independent biological replicates per condition. (E) Schematic model summarizing the proposed role of myeloid Gi signaling as a bidirectional regulator of systemic metabolism. Activation of Gi signaling engages a Gβγ–mTOR–JNK pathway associated with increased IL-1β production and impaired insulin responsiveness, whereas inhibition of Gi signaling is associated with enhanced cAMP–CREB–dependent IL-6/STAT3 signaling and improved insulin sensitivity. Together, these pathways link myeloid signaling state to systemic metabolic homeostasis.

Collectively, these findings establish myeloid Gi signaling as a bidirectional regulator of systemic metabolism, wherein Gi activation engages a Gβγ–mTOR–JNK–IL-1β inflammatory axis that drives insulin resistance, whereas Gi inhibition promotes cAMP–CREB–dependent IL-6/STAT3 signaling to enhance insulin sensitivity, thereby linking macrophage signaling state to whole-body metabolic homeostasis (**Figure 7E**).

## Discussion

Individuals with equivalent degrees of adiposity diverge markedly in insulin sensitivity and susceptibility to T2D [49–53], revealing a fundamental limitation of fat-centric models of metabolic disease. This highlights a critical role for non-endocrine cells, particularly immune cells, in shaping systemic metabolic outcomes. The present study addresses this gap by identifying myeloid Gi signaling as a previously unrecognized determinant of whole-body insulin sensitivity that operates independently of body weight and energy balance. Using bidirectional genetic and chemogenetic modulation of myeloid Gi signaling, we demonstrate that this pathway functions as a molecular switch. Inhibition of Gi signaling engages a cAMP–CREB–IL-6 axis that improves insulin sensitivity across both lean and obese states, whereas its activation in the context of diet-induced obesity recruits a Gβγ–mTOR–JNK cascade that drives IL-1β production and systemic insulin resistance. These findings establish macrophage Gi signaling as a context-dependent immunometabolic regulator that links nutritional state to cytokine production and metabolic outcomes.

The demonstration that myeloid-specific Gi inhibition improves glucose tolerance and insulin sensitivity without altering body weight or energy expenditure positions macrophage Gi signaling within the growing category of adiposity-independent immunometabolic regulators. This is consistent with the established principle that the qualitative inflammatory state of adipose tissue macrophages (ATMs), rather than their numerical expansion, is the principal determinant of metabolic outcome [54, 55]. ATMs can comprise up to 40% of the stromal vascular fraction in obese adipose tissue [12], yet metabolically healthy obese individuals maintain near-normal insulin sensitivity despite comparable macrophage infiltration, a dissociation that has focused attention on cell-intrinsic signaling programs as the relevant variables. The present findings extend this concept to the level of a G-protein pathway, demonstrating that its activity state can dictate the systemic metabolic phenotype independently of macrophage number. Importantly, *Gnai* isoform expression was markedly elevated in macrophages from diet-induced obese mice and positively correlated with blood glucose levels, raising the possibility that upregulation of myeloid Gi signaling is not an epiphenomenon of the obese microenvironment but a functionally active amplifier of the inflammatory programs that sustain metabolic dysfunction. However, whether the *Gnai* upregulation is driven by lipotoxic signals, pattern recognition receptor activation, or hypoxia-induced transcriptional reprogramming in the obese adipose niche remains to be established.

The context dependence of Gi activation is specifically revealing. Under regular chow conditions, acute Gi activation modestly improved glucose clearance without affecting insulin sensitivity or other metabolic parameters, indicating that limited Gi-driven inflammatory signaling may transiently enhance glucose disposal in the absence of metabolic stress. However, in the setting of diet-induced obesity, where chronic low-grade inflammation has already been established, both acute and chronic Gi activation impaired glucose tolerance and insulin action, independent of body weight (**Figure 2**). This diet-dependent pathogenicity may help reconcile conflicting reports in the field regarding the role of macrophage inflammation in metabolism, where identical signaling pathways can be adaptive or deleterious depending on nutritional and temporal context [56].

The mechanistic basis of Gi-driven metabolic deterioration centers on a Gβγ–mTOR/AKT–JNK signaling axis that ultimately drives IL-1β induction and secretion. While Gβγ-dependent PI3Kγ–AKT engagement is established in the context of chemokine-driven immune cell migration [57] [58, 59]. our findings reveal an additional, non-canonical signaling route. Specifically, we show that Gβγ can engage AKT and mTOR independently of classical PI3K activity and promote JNK-dependent IL-1β production under conditions of nutrient excess (**Figure 2**). In BMDMs, Gi activation engaged AKT and mTOR without detectable PI3K involvement, suggesting that this arm of Gβγ signaling is PI3K-independent in the macrophage context. Co-IP experiments in HEK293T cells showed PI3K engagement with mTOR, however, wortmannin failed to disrupt Gβ–mTOR co-precipitation (**Figure S12E**). The transient PI3K co-precipitation observed in HEK293T cells but not in BMDMs likely reflects cell-type-specific differences in G-protein effector stoichiometry rather than a discrepancy in pathway architecture. Importantly, this work identifies a TLR4-independent mode of myeloid JNK activation, as upstream regulators of this pathway beyond classical pattern recognition receptor biology. Given that JNK-mediated inhibitory serine phosphorylation of IRS-1 is required for obesity-induced insulin resistance in vivo [60] [61], our findings position macrophage Gi signaling upstream of this pathogenic axis. In this context, Gi signaling integrates GPCR-mediated inputs with nutrient-derived signals to amplify inflammatory outputs during diet-induced obesity. Thus, macrophage Gi signaling emerges as a key upstream regulator linking G protein–coupled receptor signaling to JNK-driven inflammation and systemic insulin resistance.

The identification of IL-1β as the primary effector of this pathway is consistent with extensive evidence linking macrophage-derived IL-1β to insulin resistance. IL-1β suppresses IRS-1 expression through IKKβ–NF-κB signaling at both transcriptional and post-transcriptional levels [62], and NLRP3 inflammasome–mediated IL-1β production has been directly linked to lipotoxicity-induced insulin resistance in adipose tissue macrophages [63]. Importantly, pharmacological IL-1 receptor antagonism with anakinra fully reversed both acute and chronic Gi-driven metabolic dysfunction in vivo, providing direct evidence that IL-1β is the necessary effector of pathological Gi signaling rather than a parallel correlate. The rescue was more complete in the acute than the chronic setting, likely reflecting the contribution of additional pro-inflammatory mediators, including CXCL1 and MIP-1α, that accumulate during sustained Gi activation and are not targeted by IL-1R inhibition. It is worth acknowledging the physiological duality of IL-1β: at low concentrations and in the acute postprandial setting, IL-1β amplifies glucose-stimulated insulin secretion through islet IL-1 receptor signaling, an effect that is abolished in Il1r1-deficient mice on a HFD [64, 65]. Our findings are consistent with this context-dependent function and position macrophage Gi signaling as an upstream regulator that determines whether IL-1β remains within an adaptive range or shifts toward sustained production that impairs insulin action. In this context, the CANTOS trial, in which IL-1β neutralization reduced cardiovascular events independently of lipid lowering [66], provides clinical support for targeting IL-1β in inflammatory metabolic disease. Whether elevated myeloid *Gnai* expression or activity can identify patients most likely to benefit from such interventions remains to be determined.

In contrast to Gi activation, inhibition of myeloid Gi signaling engaged a protective metabolic program initiated by relief of Gi-mediated adenylyl cyclase suppression, elevation of intracellular cAMP, and CREB-dependent transcriptional induction of IL-6 (**Figure 6**). Notably, plasma profiling revealed that TNF-α and IL-1β levels remained unchanged in LysM-PTX mice, indicating that this response is not a generalized inflammatory activation but a selective, CREB-driven cytokine program centered on IL-6. Moreover, elevated cAMP–CREB signaling in macrophages is associated with broad anti-inflammatory reprogramming, including induction of IL-10, IL-1 receptor antagonist, and IL-6 [67, 68]. Although chronically elevated circulating IL-6 engages drives systemic insulin resistance [69], an emerging body of evidence supports a context-dependent, cell-type–specific role for IL-6 in metabolic regulation. For example, IL-6 released during exercise enhances AMPK activation and glucose uptake in skeletal muscle [46], whereas myeloid-derived IL-6 has been shown to improve metabolic homeostasis and limit inflammatory macrophage recruitment [70, 71] suggesting that the cellular source and signaling context critically determine its metabolic effects. Consistent with this framework, IL-6 produced by macrophages following Gi inhibition engaged STAT3 and AMPK in both inguinal white adipose tissue[72, 73] and skeletal muscle[74, 75], supporting insulin action across depots. A tissue-specific divergence was apparent, as iWAT and skeletal muscle displayed robust STAT3 phosphorylation, whereas eWAT showed preferential AMPK activation with comparatively significant STAT3 reduction (**Figure 6**).

We hypothesize that this reflects differences in IL-6 receptor expression levels and co-receptor availability across depots, a possibility supported by reported heterogeneity in the distribution of gp130 and soluble IL-6Rα across adipose depots [70], though direct mechanistic interrogation awaits future investigation. In line with this, exploratory ATAC-seq analysis of peritoneal macrophages revealed increased chromatin accessibility at loci associated with the cAMP–CREB and IL-6 signaling axes, including Creb1, Crebbp, Prkaca, Stat3, Jak2, and Il6ra. It should be noted that each ATAC-seq data point represents a pool of three biological replicates, which, while necessary to achieve sufficient chromatin input, may introduce within-group variation and should be considered when interpreting the magnitude of individual accessibility differences. Nevertheless, the overall pattern of chromatin opening at IL-6 pathway loci is consistent with the functional and cytokine data (**Figure 7**). Increased accessibility at the Il6ra locus in particular suggests enhanced transcriptional potential for IL-6 receptor expression, which is relevant given that IL-6 signals through both classical membrane-bound IL-6Rα and trans-signaling via soluble IL-6Rα, modes that have been associated with distinct metabolic and inflammatory outcomes (**Figure 7**) [76, 77]. The observed increase in Il6ra accessibility, together with elevated myeloid IL-6 production, is consistent with engagement of a metabolically beneficial IL-6 signaling axis. Critically, pharmacological IL-6 receptor blockade with LMT-28 completely abolished the metabolic benefits of myeloid Gi inhibition under both RC and HFD conditions, establishing IL-6 as a necessary (**Figure 6**), non-redundant effector of the protective arm of this switch rather than a correlative response.

In summary, this study establishes macrophage Gi signaling as a weight-independent immunometabolic switch that couples opposing intracellular states to antagonistic cytokine programs, cAMP–CREB–IL-6 versus Gβγ–AKT/mTOR–JNK–IL-1β with reciprocal consequences for systemic insulin sensitivity. This framework provides a mechanistic explanation for how identical macrophage abundance can produce divergent metabolic outcomes across individuals, and why inflammatory dysfunction can persist in the absence of excess adiposity. Given that myeloid *Gnai* expression correlates with glycaemic parameters and that anakinra, a clinically approved agent, rescues Gi-driven metabolic dysfunction in vivo, these findings carry direct translational relevance. More broadly, they suggest that targeting immune cell-intrinsic G-protein signaling may offer a more proximal and selective therapeutic strategy than cytokine blockade alone, capable of restoring metabolic homeostasis by correcting dysregulated inflammatory output at its intracellular origin.

## Methods

### Generation of Mutant Mice

To generate transgenic mice with selective GiD (DREADDs that stimulate Gi signaling upon activation) expression in myeloid cells, LysM-Cre mice were crossed with R26-LSL-Gi-DREADD mice [32]. LysM-Cre mice express Cre recombinase under the control of the mouse lysozyme 2 promoter that is active specifically in the myeloid cell lineage (JAX Stock No. 004781). R26-LSL-Gi-DREADD mice were obtained from Jackson Laboratories (Stock No. 026219). Cre expression in myeloid cells efficiently excises the floxed STOP cassette, resulting in selective GiD expression in myeloid cells [31]. For clarity, these mice are referred to as LysM-GiD, and littermates carrying one allele of R26-LSL-Gi-DREADD but lacking Cre were used as controls.

To selectively inhibit Gi signaling in myeloid cells, LysM-Cre mice were crossed with R26-LSL-PTX(S1) mice [35]. The R26-LSL-PTX(S1) line (MMRRC_030678-UCD) carries a Cre-inducible allele encoding the catalytically active S1 subunit of pertussis toxin. Cre-mediated deletion of the STOP cassette enabled myeloid-specific expression of PTX(S1), thereby disrupting Gi-coupled GPCR signaling exclusively in myeloid cells. These mice are referred to as LysM-PTX, and littermates lacking Cre served as controls. For G-protein expression analyses, C57BL/6NJ mice (JAX Stock No. 005304) were used.

All studies were performed using adult male littermate mice unless otherwise indicated. Animals were housed at 23 °C on standard chow (15% kcal from fat; 3.1 kcal/g) with ad libitum access to food and water under a 12-h light/dark cycle. To induce diet-induced obesity, male mice aged 6–8 weeks were switched to a high-fat diet (60% kcal from fat; Research Diets) and maintained on this diet for at least eight weeks prior to metabolic testing. Body weight and food intake were monitored weekly. All procedures adhered to institutional and national ethical guidelines, and protocols were approved by the Institutional Animal Ethics Committee (IAEC), Indian Institute of Technology Kanpur, India.

### In Vivo Metabolic Tests

Glucose homeostasis was assessed using a series of standardized metabolic tolerance tests. For glucose tolerance tests (GTT), mice were fasted overnight (14–16 h) and administered glucose intraperitoneally (1 or 2 g/kg) [78]. Blood glucose was measured from the tail vein at defined time points using glucometer (Glucometer Elite Sensor; Bayer) to evaluate whole-body glucose clearance. Peripheral insulin sensitivity was assessed by insulin tolerance tests (ITT) [78]. Mice were fasted for 4–6 h, injected intraperitoneally with recombinant human insulin (0.75 or 1 U/kg i.p.; Humulin, Eli Lilly), and serial blood glucose measurements were obtained to monitor the rate of glucose lowering. To examine hepatic gluconeogenic capacity, pyruvate tolerance tests (PTT) were performed following an overnight fast. Sodium pyruvate (2 g/kg) was administered intraperitoneally, and glucose excursions were monitored over time, allowing us to evaluate the impact of myeloid Gi-signaling blockade on liver glucose production [78]. Glucose-stimulated insulin secretion (GSIS) was assessed in mice fasted for 4–6 h. Animals received an intraperitoneal bolus of glucose (1 or 2 g/kg), and blood samples were collected at designated time points post-injection [78]. Plasma insulin levels were measured using a mouse insulin ELISA kit (Crystal Chem Inc./ ALPCO STELLUX) following the manufacturer’s instructions.

### Acute and chronic DCZ treatment in LysM-GiD mice

For acute activation, LysM-GiD mice and their littermate controls were fasted overnight and administered DCZ (30 µg/kg, i.p.) [79]. Blood glucose levels were measured from the tail vein at the indicated time points (−30, 0, 15, 30, 60, 90, and 120 min). For all subsequent metabolic tolerance tests, fasting protocols were identical to those described above. DCZ (30 µg/kg, i.p.) was administered 30 min prior to glucose, insulin, or pyruvate injection, and blood glucose concentrations were recorded at regular intervals throughout the test.

For chronic chemogenetic activation, control and LysM-GiD mice were maintained on a high-fat diet for 12 weeks and then treated with DCZ (30 µg/kg, i.p.) once daily for seven consecutive days. On day 8, metabolic assessments were performed following the established protocols.

### Plasma analysis

Blood was collected from control and LysM-GiD groups in heparinized tubes at the indicated time points following treatment. Samples were centrifuged at 12,000 × g for 10 min at 4 °C, and the resulting plasma was used for biochemical analyses. Plasma glycerol and triglyceride levels were measured using assay kits from Sigma-Aldrich, while free fatty acids (FFAs) were quantified using kits from Fujifilm Wako Diagnostics.

Circulating cytokines, including IL-6, IL-1β and TNF-α were assessed using ELISA kits (BioLegend/Bio-Rad) according to the manufacturer’s protocols.

### Acute and chronic Anakinra treatment in LysM-GiD and control mice

*Acute treatment protocol:* Control and LysM-GiD mice maintained on an HFD for 8 weeks were divided into two groups, with one receiving Anakinra and the other receiving PBS [80]. For acute IL-1β inhibition, mice received an initial intraperitoneal injection of Anakinra (2.5 mg/kg) 12–14 hours before the metabolic experiment, followed by a second dose (2.5 mg/kg, i.p.) administered 2 hours prior to the metabolic test [80]. DCZ (30 µg/kg, i.p.) was injected 30 minutes before initiating the GTT or ITT. At time 0, mice received either glucose (2 g/kg, i.p.) or insulin, depending on the assay. Blood glucose levels were measured from the tail vein at baseline and at 15, 30, 60, 90, and 120 minutes following the glucose or insulin injection.

*Chronic treatment protocol:* For chronic chemogenetic activation and cytokine blockade, control and LysM-GiD mice were maintained on an HFD for 12 weeks. DCZ (30 µg/kg, i.p.) was administered every 3 days throughout the treatment period. Mice receiving chronic Anakinra were injected with 5 mg/kg (i.p.) every 3 days in the days leading up to metabolic testing [80]. Immediately before the 12–14-hour overnight fast preceding metabolic assessments, mice received an elevated Anakinra dose (10 mg/kg, i.p.). A final dose of Anakinra (5 mg/kg, i.p.) was delivered 2 hours before the glucose or insulin challenge. Metabolic tests were performed as described above.

### LMT-28 administration and metabolic testing

Control and LysM-PTX mice were randomly assigned to receive either vehicle (DMSO diluted in corn oil) or the IL-6 receptor antagonist LMT-28, generating four experimental groups. LMT-28 was dissolved in DMSO, further diluted in corn oil according to the manufacturer’s protocol, and administered intraperitoneally at a dose of 5 mg/kg on alternate days for a total of three injections [81]. Following the final dose, GTT was performed according to established protocols. For GTT, mice were fasted overnight and injected intraperitoneally with glucose (1–2 g/kg). Blood glucose levels were measured from the tail vein at defined time points using a glucometer.

### Bone marrow-derived macrophages isolation and culture

Bone marrow cells were harvested from the tibia and femur of control and LysM-GiD mice by flushing the bones with Dulbecco’s Modified Eagle Medium (DMEM) using a 10-mL syringe. The resulting cell suspension was centrifuged at 2,000 rpm for 5 minutes, and the pellet was resuspended in ACK lysis buffer and incubated for 5 minutes to eliminate red blood cells. Then the cell suspension was filtered through a 70-µm cell strainer and centrifuged again at 2,000 rpm for 5 minutes [82]. Cells were resuspended in DMEM supplemented with 20 ng/mL M-CSF and plated in 6-well plates. After 24 hours, the medium was replaced with fresh DMEM containing 40 ng/mL M-CSF to remove non-adherent cells. Cultures were maintained for 7 days to allow differentiation into bone marrow–derived macrophages, which were subsequently used for downstream experiments.

### Western-blot studies

Bone marrow–derived macrophages from LysM-GiD and control mice were cultured in 6-well plates and stimulated with DCZ (100 nM) for 30 minutes. Cells were then lysed in radioimmunoprecipitation assay (RIPA) buffer supplemented with cOmplete EDTA-free protease and phosphatase inhibitor cocktail (Sigma-Aldrich) [78]. Lysates were centrifuged at 18,000 × g for 15 minutes at 4 °C, and the resulting supernatants were transferred to fresh 1.5-ml tubes. Protein concentrations were quantified using the BCA protein assay kit (Pierce) [78]. Protein concentration was normalized to 2 mg/ml, denatured at 95 °C, and resolved on 4–12% SDS–polyacrylamide. Proteins were transferred to PVDF membranes, blocked in 5% BSA at room temperature, and then incubated overnight at 4 °C with primary antibodies [78]. The following day, membranes were washed thoroughly and incubated with HRP-conjugated anti-rabbit secondary antibodies. Protein bands were visualized using ECL substrate (Bio-Rad) on the Azure600 Imaging System (Azure Biosystems). Immunoreactive bands were quantified using ImageJ software.

For inhibitor studies, bone marrow–derived macrophages were pretreated with pathway-specific inhibitors before DCZ stimulation. Cells were incubated with rapamycin (10 nM) for 12 hours to block mTORC1 signaling[83], wortmannin (10 nM) for 16 hours to inhibit PI3K activity [84], or gallein (10 µM) for 1 hour to inhibit Gβγ signaling[85]. Corresponding vehicle controls were included for all conditions. Following pretreatment, cells were stimulated with DCZ (100 nM) for 30 minutes and processed for Western blot analysis as described above.

For tissues immune-blotting, Inguinal white adipose tissue (IWAT), epididymal white adipose tissue (EWAT), quadriceps (SKM-Q), and gastrocnemius (SKM-G) muscles were rapidly dissected, rinsed briefly in ice-cold PBS, snap-frozen in liquid nitrogen and stored at −80 °C for further processing [86]. Frozen tissue was homogenized using FastPrep®-24 homogenizer (MP Biomedicals) in radioimmunoprecipitation assay (RIPA) buffer supplemented with cOmplete EDTA-free protease and phosphatase inhibitor cocktail (Sigma-Aldrich) [78]. Homogenates were cleared by centrifugation at 12,000 × g for 15 min at 4 °C; the aqueous supernatant was collected and protein concentration determined by BCA protein assay kit (Pierce) [78]. Equal amounts of protein were then prepared in Laemmli buffer and subjected to Western blot analysis as described above.

### Bulk RNA sequencing

Bone marrow–derived macrophages from LysM-GiD mice were serum-starved for 1 hour prior to stimulation. Cells were then treated with DCZ (100 nM) for 1, 2, or 4 hours, with untreated cells (0 hours) processed in parallel as controls. Following treatment, cells were washed with PBS, and total RNA was extracted using the ReliaPrep™ RNA miniprep system (Promega) following the manufacturer’s instructions. RNA concentration, purity, and integrity were assessed and only samples that met quality control criteria were advanced for sequencing. Sequencing was performed by Nucleome Informatics Private Limited, Hyderabad, India.

### Bulk RNA seq Data analysis

Raw paired-end FASTQ files generated from bulk RNA sequencing were subjected to quality assessment using FastQC (reference). Adapter contamination and low-quality bases were removed using Trimmomatic (https://trimmomatic.com/), retaining only high-quality reads for downstream analysis. Quality-filtered reads were aligned to the reference genome (e.g., mm10) using Hisat2 [87]. Aligned reads in BAM format were quantified at the gene level using featureCounts [88]. Reads were assigned to genomic features based on the reference GTF annotation, and only properly paired reads were counted. The resulting gene-level count matrix was used as input for differential expression analysis. Differential gene expression analysis was performed using DESeq2 in R with the default pipeline [89]. Raw gene counts were to account for differences in sequencing depth. Genes with an adjusted p-value < 0.05 and an absolute log2 fold change exceeding the specified threshold of 0.5 were considered significantly differentially expressed. Volcano plots were visualized using the Enhanced volcano package, and the heatmaps were visualized using the pheatmap package in R. To perform the pathway analysis, gseGO package was used.

### scRNA data analysis

Publicly available mouse adipose tissue scRNA datasets (GSE128518, GSE160729, GSE176171, GSE183288) were retrieved from Gene Expression Omnibus. After merging all the datasets, adipose tissue cells were integrated batchwise using ‘scDREAMER’ [36]. Following the identification of major cell types, Immune cell population was extracted and reintegrated using ‘scDREAMER’. Immune cell types were annotated based on established marker genes, after which macrophages were subsetted and reintegrated for focused analysis of Gi gene expression. Downstream single-cell analyses, including Normalization, variable marker finding, scaling was performed in R using ‘Seurat’ package [90]. For human adipose tissue analysis, processed RDS objects from publicly available datasets (GSE176067 and GSE176171) were obtained from the White Adipose Atlas GitHub repository (https://gitlab.com/rosen-lab/white-adipose-atlas). Human macrophages were subsetted from these datasets and analysed for Gi gene expression. Visualization tools, including UMAPs, bar plots, dot plots, and heatmaps, were generated using the SeuratExtend package.

### Differentiation of 3T3F442A and C2C12 cells

Mouse 3T3-F442A preadipocytes were maintained in DMEM supplemented with 10% bovine calf serum (BCS) and 1% penicillin–streptomycin at 37 °C in a humidified 5% CO₂ incubator [91]. Once the cells were fully confluent, adipogenic differentiation was initiated by treating cells for 48 hours with induction cocktail containing 0.5 µM insulin, 250 µM IBMX, 2 µM troglitazone, 0.5 µM dexamethasone, and 60 µM indomethacin [91]. After induction, the medium was replaced with DMEM containing 10% BCS, 1% penicillin–streptomycin, and 0.5 µM insulin, and cells were maintained for an additional 48 hours to complete differentiation [91]. Mature adipocytes were subsequently used for downstream analyses.

Mouse C2C12 myoblasts were maintained on DMEM supplemented with 10% FBS and 1% penicillin–streptomycin at 37 °C in a humidified 5% CO₂ incubator. For differentiation, cells were grown to 80–90% confluence and then switched to DMEM containing 2% horse serum. Cultures were allowed to differentiate for 5–7 days, with medium refreshed every 48 hours. Differentiated myotubes were collected for downstream analyses [92].

### Co-culture of 3T3-F442A and C2C12 with macrophage-conditioned medium

Control and LysM-GiD BMDMs were generated and cultured in 6-well plates as described above. Cells were serum-starved for one hour and treated with DCZ (100 nM) for an additional hour. Following stimulation, macrophage-conditioned media were collected and transferred to mature 3T3-F442A adipocytes or differentiated C2C12 myotubes, which had been serum-starved for one hour before treatment. After a one hour incubation with macrophage-conditioned medium, adipocytes and myotubes were stimulated with 10 nM insulin for 10 minutes or 30 minutes, respectively, and subsequently processed for downstream analyses [91].

### Measurement of mouse plasma cytokine levels

Blood was collected from the mouse tail vein into K2-EDTA tubes and quickly centrifuged at 4°C to obtain plasma. Plasma cytokine levels were measured using the procartaplex cytokine/chemokine panel from Invitrogen according to the manufacturer’s instructions [78]. The concentrations of cytokines were determined using the bio-plex MAGPIX multiplex reader as specified by the manufacturer.

### PBMC isolation and culture

Human peripheral blood was collected from healthy donors in accordance with guidelines approved by the Institutional Ethics Committee, IIT Kanpur. Approximately 20 mL of venous blood was drawn into heparinized vacutainers (BD Vacutainer, Sodium Heparin, 158 USP units) and diluted 1:1 with phosphate-buffered saline (PBS) [93]. Peripheral blood mononuclear cells (PBMCs) were isolated by density-gradient centrifugation using Histopaque PLUS (Sigma-Aldrich) at 1,200 × g for 10 minutes at room temperature with no brake applied [93]. The PBMC layer was carefully recovered, washed twice with PBS (300 × g for 10 minutes), and resuspended in RPMI-1640 medium (Gibco) supplemented with 20 ng/mL M-CSF [93]. Cells were cultured for three days, after which fresh RPMI containing 20 ng/mL M-CSF was added. Cultures were maintained under these conditions until further use.

### Real-time qRT-PCR gene expression analysis

Mouse tissues collected from experimental (LysM-GiD and LysM-PTX) and control mice were rapidly excised and snap-frozen. Total RNA was isolated using the RNeasy Mini Kit (Qiagen) with on-column DNase treatment, following the manufacturer’s instructions. cDNA was synthesized using the PrimeScript™ First Strand cDNA Synthesis Kit, and quantitative real-time PCR was performed using SYBR Green chemistry (Takara). Gene expression levels were normalized to β-actin or 18S rRNA. For in vitro gene expression analyses, control and LysM-GiD BMDMs isolated from mice maintained on HFD for 4–6 weeks were cultured in 6-well plates. Cells were serum-starved for 1 hour and treated with DCZ (100 nM) for 1 hour. Total RNA was extracted using the ReliaPrep™ RNA Miniprep System, and cDNA synthesis and qPCR were performed as described above.

### H&E staining

Inguinal (iWAT) and epididymal (eWAT) white adipose tissues were collected from control and LysM-GiD mice following chronic DCZ injections, fixed in 10% formalin at room temperature, and processed for paraffin embedding. Tissue blocks were sectioned at 10 μm thickness and mounted on glass slides. Sections were deparaffinized, rehydrated through graded alcohols, and stained with hematoxylin and eosin (H&E). Brightfield images were acquired using a NIKON light microscope under identical settings across all samples.

### Measurement of intracellular cAMP in peritoneal macrophages

Intracellular cAMP levels were measured using the HTRF cAMP Gi kit (Revvity) according to the manufacturer’s instructions with minor modifications. Peritoneal macrophages were isolated from control and LysM-PTX mice and resuspended in complete DMEM supplemented with IBMX to prevent phosphodiesterase-mediated cAMP degradation. Cells were allowed to adhere for 2 h in non–tissue culture–treated plates at 37°C in a humidified incubator containing 5% CO₂. Following adhesion, non-adherent cells were removed, and macrophages were harvested, counted, and replated in duplicate at a density of 8,000 cells per well in assay plates. Cells were then stimulated with forskolin at the indicated concentrations (0, 0.1, 1, or 10 μM) for 15 min at 37°C. After stimulation, cells were lysed and cAMP levels were quantified using the HTRF cAMP Gi detection reagents, and fluorescence resonance energy transfer signals were measured using a BMG plate reader. cAMP concentrations were calculated from standard curves generated in parallel and expressed as absolute intracellular cAMP levels.

### Determination of IL-6 levels in macrophages from LysM-PTX mice

Peritoneal macrophages were isolated from control and LysM-PTX mice by peritoneal lavage using complete DMEM supplemented with 50 µM 3-isobutyl-1-methylxanthine (IBMX) to preserve intracellular signaling states. Cells were pelleted at 500 × g for 5 minutes, resuspended in fresh complete medium containing 50 µM IBMX, and plated onto 35-mm non–tissue-culture–treated dishes. Macrophages were allowed to adhere for 2 hours at 37 °C, after which non-adherent cells were removed by gentle PBS washing. Adherent cells were then treated with or without 10 µM FRS in the continued presence of 50 µM IBMX for 1 hour before processing for RNA isolation.

For IL-6 secretion assays, peritoneal macrophages isolated as described above were resuspended in complete DMEM containing 50 µM IBMX and plated in 96-well flat-bottom plates at 5 × 10⁴ cells per well. Cells were maintained in IBMX-supplemented medium and treated with vehicle, 0.1 µM FRS, or 1 µM FRS. Following a 24-hour incubation, culture supernatants were collected and clarified by centrifugation (500 × g, 5 minutes). IL-6 concentrations in the supernatants were quantified by ELISA according to the manufacturer’s instructions (Biolegend).

### High content microscopy

Cells were seeded in 96-well plates and transfected with plasmids as described in the figure legends. After overnight transfection, cells were treated with DCZ as indicated in the figures. Once treatment was complete, cells were fixed with 4% paraformaldehyde for 5 minutes. The cells were then permeabilized with 0.1% Triton X-100 and blocked with 3% BSA for 45 minutes, followed by overnight incubation with primary antibody at 4°C. Cells were washed three times with PBS, then incubated with Alexa Fluor-conjugated secondary antibody for 1 hour at room temperature. High-content microscopy with automated image acquisition and quantification was performed using a Cellomics HCS scanner and iDEV software (Thermo) in 96-well plates. For analysis, at least 6 wells per plate were used, and a minimum of 500 cells were analyzed per well for each sample.

### Co-Immunoprecipitation assay

Cells were seeded in 100mm cell culture dishes and, where specified, transfected with 10 µg of plasmid. Cells were then lysed in NP-40 buffer containing a protease inhibitor cocktail (Roche, cat# 11697498001) and PMSF (Sigma, cat# 93482) and incubated on ice for 45 minutes. Subsequently, samples were centrifuged at 12,000g for 15 minutes at 4°C, and the supernatant containing cell lysates was collected. Proteins were quantified using a Bradford assay. 5-10% of the lysates were reserved as input controls. The remaining protein lysates (approximately 1.5 mg) were incubated with 5 µg of antibody overnight at 4°C, followed by incubation with Dynabeads Protein G (Invitrogen) for 4 hours at 4°C. Beads were washed three times with PBS and then boiled with SDS-PAGE buffer for immunoblotting to analyze interacting proteins.

### Assay for Transposase-Accessible Chromatin (ATAC-seq) library preparation

Peritoneal macrophages (PMs) were isolated from control and LysM-PTX as described above. Approximately 2 × 10⁶ cells were used for nuclei isolation. Nuclei were extracted from adherent PMs using a modified nuclei isolation protocol adapted from established single-nucleus ATAC-seq methods (Dhall J.K et al, 2023). Briefly, cells were gently scraped, collected, and pelleted at 500 × g for 5 min at 4°C. Cell pellets were resuspended in cold lysis buffer (10 mM Tris-HCl, pH 7.4; 10 mM NaCl; 3 mM MgCl₂; 0.1% NP-40) to selectively permeabilize the plasma membrane while preserving nuclear integrity. Following incubation on ice for 15 min, nuclei were washed with wash buffer (10 mM Tris-HCl, pH 7.4; 10 mM NaCl; 3 mM MgCl₂; 0.1% Tween-20) and passed through a 40-µm cell strainer to remove cellular debris. Isolated nuclei were pelleted at 500 × g for 5 min at 4°C and resuspended in diluted nuclei buffer for counting. For ATAC-seq library preparation, 75,000 nuclei per biological sample were used for transposition. Nuclei were pelleted at 500 × g for 5 min at 4°C and resuspended in the tagmentation reaction mix containing Tn5 transposase and associated buffers according to the ATAC-seq library preparation kit manufacturer’s instructions. Transposition was performed at 37°C for 45 min with gentle mixing. Following tagmentation, DNA was purified using AMPure XP beads as specified in the protocol and PCR-amplified with indexed primers to generate sequencing libraries. The number of PCR cycles was determined by qPCR pre-amplification to minimize over-amplification. Library quality was assessed by Qubit fluorometric quantification and fragment size distribution analysis using a Bioanalyzer. Libraries exhibiting a characteristic nucleosomal ladder pattern and appropriate fragment size profiles were pooled equimolarly and submitted for paired-end high-throughput sequencing on an Illumina platform following standard manufacturer recommendations.

### ATAC-seq data analysis

ATAC-seq libraries were generated from peritoneal macrophages isolated from LysM-PTX-S1 (LysM-Cre) WT mice. Due to limited cell yield, cells from three mice were pooled to generate each biological replicate, resulting in two pooled replicates per condition. Raw sequencing reads were processed using a Snakemake-based workflow. Adapter trimming was performed using Trim Galore (v0.6.10; Cutadapt v4.x) with parameters ‘--nextera ––quality 20 ––length 20’. Trimmed reads were aligned to the mouse reference genome (GRCm39; GENCODE vM33) using Bowtie2 (v2.5.3) with parameters ‘--very-sensitive ––no-mixed ––no-discordant –X 1000’. Reads with mapping quality <20, reads mapping to mitochondrial DNA (chrM), and reads overlapping ENCODE blacklist regions were removed using SAMtools (v1.19) and BEDtools (v2.31.1). PCR duplicates were removed using Picard (v3.1.1).

To accurately represent transposase insertion events, read alignments were adjusted for the Tn5 integration offset (+4 bp for positive strand and −5 bp for negative strand) using deepTools (v3.5.4; alignmentSieve). Peaks were called using MACS3 (v3.0.4) in paired-end mode (‘BAMPÈ, ‘-q 0.05’, ‘--nomodel’, ‘--shift –100’, ‘--extsize 200’, ‘--nolambdà). A consensus peak set was defined by retaining peaks reproducibly detected within each condition (present in both pooled biological replicates) and merging these sets across conditions. Read counts within consensus peaks were quantified using summarizeOverlaps (GenomicAlignments; Union mode, strand-agnostic, paired-end). Differential chromatin accessibility was assessed using DESeq2 (v1.42) with a design formula of ‘∼Condition’, and log2 fold changes were shrunk using the ashr method. Peaks with false discovery rate (FDR) < 0.05 and absolute log2 fold change ≥ 0.5 were considered differentially accessible. Genome-wide signal tracks were generated using RPGC normalization (effective genome size = 2,494,787,188 bp; bin size = 10 bp). Transcription factor motif enrichment analysis was performed using HOMER (v4.11) using mm39 genome sequence, 200 bp windows, and the consensus peak set as background. Gene set enrichment analysis was performed using fgsea (v1.28; nPermSimple = 10,000) on DESeq2 Wald statistics ranked by nearest-gene assignment.

## Statistical analysis

Data are presented as mean ± standard error of the mean (s.e.m.). Statistical significance was assessed using unpaired two-tailed Student’s t-tests or one-way ANOVA, as appropriate, using GraphPad Prism. A p-value < 0.05 was considered statistically significant.

## Acknowledgements

We thank Dr. Sabarinathan Radhakrishnan and members of his laboratory for their expert assistance with ATAC-seq library preparation and data analysis. We are grateful to Prof. Bryan Roth (University of North Carolina, Chapel Hill, USA) for generously providing the Rosa26-LSL-Gi-DREADD mouse line. The B6;129P2-Gt(ROSA)26Sor^tm1(ptxA)Cgh/Mmucd strain (RRID: MMRRC_030678-UCD) was obtained from the Mutant Mouse Resource and Research Center (MMRRC) at the University of California, Davis, an NIH-funded repository, and was originally donated by Dr. Shaun Coughlin (University of California, San Francisco). We thank all current and past members of the MMCSL laboratory for stimulating discussions and critical input throughout the course of this work, and the Central Experimental Animal Facility at IIT Kanpur for support with in vivo studies.

This work was primarily supported by the DBT/Wellcome Trust India Alliance (IA/I/21/1/505613) and the Department of Biotechnology, Government of India (DBT; B1/PR44526/MED/30/2376/2021). Research in the SPP laboratory is additionally supported by the Department of Science and Technology–Science and Engineering Research Board (DST-SERB; CRG/2021/004502), the Indian Council of Medical Research (ICMR; 5/4/8-18/Obs/SPP/2022-NCD-II), and an Initiation Grant from IIT Kanpur. S.K. is supported by a CSIR Junior Research Fellowship (09/0092(12663)/2021-EMR-I), and H.Y. is supported by the Prime Minister’s Research Fellowship (PMRF). AI-assisted tools, including Grammarly, were used for language editing and manuscript refinement.

## Author Contribution

Experimental design was carried out by S.K., H.Y., and S.P.P. All *in vivo* studies were primarily performed by S.K.,H.Y. and S.B.. Co-IP studies were done by S.R. Su.K.. Plasma analyses were conducted by H.Y., S.K., A.S., S.S., A.K., and S.P.. Immunostaining was performed by S.K. and R.F. ATAC-Seq library and data analysis was performed by H.Y. Single-cell RNA-seq and bulk RNA seq data analysis was carried out by M.S., S.S., and H.Z. The project was conceptualized by S.P.P. The manuscript was written by S.K., H.Y., and S.P.P., with input from all authors. All authors reviewed and approved the final version of the manuscript.

## Supplementary Figures

**Figure S1: Extended analysis of myeloid Gαi expression and metabolic parameters.** (A–C) UMAP visualization and quantification of immune cell populations following integration of multiple murine adipose tissue scRNA-seq datasets (GSE128518, GSE160729, GSE176171, GSE183288) using the scDREAMER framework. Integrated datasets were subsetted to immune cells and annotated based on canonical marker genes. Macrophage abundance is increased in HFD-fed mice compared to RC controls. B) UMAP representation of human adipose tissue single-cell transcriptomic data (preprocessed and annotated). (E, F) Canonical marker gene expression used for annotation of immune cell populations in mice (E) and humans (F). (G, H) Immune cell-type abundance across conditions in mice (RC vs HFD) (G) and humans (lean vs obese) (H), demonstrating expansion of macrophage populations under metabolic stress. (I) Feature plots showing expression of *Gnai*/*GNAI* family genes across macrophage populations in mice (RC vs HFD) and humans (lean vs obese), indicating enhanced Gαi signaling in metabolically stressed states. (J) Relative expression of *Gnai* isoforms in murine bone marrow–derived macrophages (BMDMs), demonstrating *Gnai2* as the predominant isoform. (K) Relative expression of *GNAI* isoforms in human peripheral blood mononuclear cells (PBMCs), confirming conserved enrichment of *GNAI2* across species.

**Figure S2: Metabolic profiling of control and LysM-PTX mice under basal conditions**. (A–E) Plasma metabolic parameters were measured in control and LysM-PTX mice maintained on a regular chow diet. Blood glucose (A), plasma insulin (B), glycerol (C), triglyceride (TG) levels (D), and non-esterified fatty acids (NEFA) (E) were assessed in fasting and fed states. (F–I) Corresponding metabolic measurements in mice maintained on high-fat diet conditions, including blood glucose (F), plasma insulin (G), glycerol (H), and NEFA levels (I). Data are presented as mean ± s.e.m. Statistical significance was determined using a two-tailed Student’s t-test.

**Figure S3: Whole-body energy metabolism is not significantly altered by macrophage Gi signaling inhibition.** (A–H) Indirect calorimetry analysis of control and LysM-PTX mice maintained on a regular chow diet. Oxygen consumption (VO₂), energy expenditure (EE), and respiratory exchange ratio (RER) were measured over the indicated time periods at 23 °C and 30 °C. (I–P) Corresponding metabolic measurements in mice maintained on high-fat diet conditions, including VO₂, EE, and RER. Data are presented as mean ± s.e.m. Statistical significance was determined using a two-tailed Student’s t-test.

**Figure S4: Validation and functional characterization of GiD expression in macrophages.** (A) qPCR analysis of GiD expression in control and LysM-GiD bone marrow–derived macrophages (BMDMs). Gene expression was normalized to 18S rRNA. (B) Immunoblot analysis confirming expression of HA-tagged GiD protein in control and LysM-GiD BMDMs. β-actin was used as a loading control. (C) Tissue distribution of GiD expression in brown adipose tissue (BAT), epididymal white adipose tissue (eWAT), inguinal white adipose tissue (iWAT), liver, pancreas, glycolytic skeletal muscle (SKM-g), and oxidative skeletal muscle (SKM-q) from control and LysM-GiD mice. Expression levels were normalized to β-actin. (D) Functional assessment of GiD signaling using an HTRF cAMP assay in BMDMs. Cells were stimulated with forskolin and DCZ, and intracellular cAMP levels were quantified (nM). (E) Functional validation of GiD signaling in the presence of pertussis toxin (PTX) using the HTRF cAMP assay to assess Gi-dependent inhibition of cAMP production. Data are presented as mean ± s.e.m. Statistical significance was determined using a two-tailed Student’s *t*-test.

**Figure S5: Metabolic characterization of control and LysM-GiD mice maintained on regular chow following DCZ treatment.** (A) The body weight of control and LysM-GiD mice was monitored from 6 to 15 weeks of age. (B) Random-fed and fasting blood glucose levels were measured in control and LysM-GiD mice. (C) DCZ challenge test showing blood glucose levels following acute DCZ administration (30 µg/kg, i.p.) in control and LysM-GiD mice (N= 8 per group). (D) Glucose challenge test (GTT; glucose 2 g/kg, i.p.) performed following DCZ injection, with corresponding area under the curve (AUC-GTT) analysis (N= 8 per group). (E) Pyruvate challenge test (PTT; pyruvate 2 g/kg, i.p.) performed following DCZ injection, with corresponding area under the curve (AUC-PTT)(N= 8 per group). (F) Insulin challenge test (ITT; insulin 0.75 U/kg, i.p.) following DCZ injection, with quantification of area under the curve (AUC-ITT) (N= 8 per group). (G) Plasma triglyceride (TG) levels measured at indicated time points following DCZ administration (N=7 mice per group).(H) Plasma glycerol levels measured at indicated time points following DCZ administration (N=7 mice per group). Data are presented as mean ± s.e.m., C & I: two-tailed Student’s t-test.

**Figure S6: Lipid metabolic parameters and adipose tissue morphology following acute and chronic myeloid Gi signaling activation.** (A) Plasma non-esterified fatty acid (NEFA) levels measured following acute DCZ administration (30 µg/kg, i.p.) (B-C) Fast and fed plasma NEFA (B) and glycerol (C) levels measured from chronically activated Gi signaling in HFD control and lysM-GiD mice. (D) Representative hematoxylin and eosin (H&E) staining of inguinal white adipose tissue (iWAT) and epididymal white adipose tissue (eWAT) from Control and LysM-GiD mice, followed by chronic activation of Gi Signaling. Data are presented as mean ± s.e.m., two-tailed Student’s t-test.

**Figure S7: GiD activation in myeloid cells does not significantly alter whole-body energy expenditure.** (A) Time-course of oxygen consumption (VO₂) measured by indirect calorimetry in control and LysM-GiD mice following DCZ injection. (B,C) Quantification of VO₂ during the 12-h light phase (B) and 12-h dark phase (C). (D) Time-course of carbon dioxide production (VCO₂) in control and LysM-GiD mice following DCZ injection. (E,F) Quantification of VCO₂ during the light phase (E) and dark phase (F). (G) Time-course of respiratory exchange ratio (RER) in control and LysM-GiD mice following DCZ administration. (H,I) Quantification of RER during the light phase (H) and dark phase (I). (J) Time-course of energy expenditure (EE) measured by indirect calorimetry in control and LysM-GiD mice after DCZ injection. (K,L) Quantification of EE during the light phase (K) and dark phase (L). Data are presented as mean ± s.e.m. Statistical significance was determined using a two-tailed

**Figure S8: Circulating Chemokine Profile in LysM-GiD Mice Following DCZ Treatment.** (A-D) Multiplex cytokine analysis of plasma from control and LysM-GiD mice treated with vehicle or DCZ, showing circulating levels of MIP1α (A), MIP2α (B), and CCL11 (C), and CXCL10 (D).

**Figure S9: Time-dependent transcriptional changes following GiD activation.** (A–C) Volcano plots showing differential gene expression following DCZ-mediated GiD activation at 1 h (A), 2 h (B), and 4 h (C) compared with control conditions. The x-axis represents log₂ fold change, and the y-axis represents –log₁₀ (adjusted P value). Each dot represents an individual gene. Upregulated genes are shown in red, downregulated genes in blue, and non-significant genes in grey.

**Figure S10: Heatmap of 1-hour and 2-hour treated upregulated and downregulated genes.** Control and lysM-GiD BMDM cells were treated with DCZ for 1 and 2 hour and their differentially enriched genes were analysed.

**Figure S11**: (A) Gene Ontology (GO) enrichment analysis of differentially expressed genes following chemogenetic activation of Gi signaling at 2 h and 4 h post-treatment. (B–F) Quantitative PCR (qPCR) analysis of selected genes associated with inflammatory signaling, metabolic regulation, transcriptional control, and chemokine expression in control and LysM-GiD BMDMs following treatment. (G) qPCR analysis of gene expression in epididymal white adipose tissue (eWAT) and brown adipose tissue (BAT) from HFD-fed control and LysM-GiD mice after acute activation of Gi signaling. Data are presented as mean ± s.e.m. Statistical significance was determined using a two-tailed Student’s *t*-test.

**Figure S12: Gi signaling regulates the AKT–mTOR and JNK signaling pathways in macrophages.** (A) Immunoblot analysis of phosphorylated PI3K (P-PI3K) and total PI3K (T-PI3K) in Control and LysM-GiD macrophages following DCZ stimulation. β-actin was used as a loading control. (B) Immunoblot analysis of signaling pathway activation in control and LysM-GiD macrophages following DCZ treatment in the presence or absence of pertussis toxin (PTX). Quantification of the phosphorylated proteins normalized to their respective total protein levels is shown below. (C) Immunoblot analysis of P-PI3K and T-PI3K in Control and LysM-GiD macrophages treated with DCZ in the presence or absence of the PI3K inhibitor wortmannin or the mTOR inhibitor rapamycin. β-actin serves as a loading control. (D) Immunoblot analysis of P-PI3K and T-PI3K in Control and LysM-GiD macrophages treated with DCZ in the presence or absence of the Gβγ inhibitor gallein. β-actin was used as a loading control. (E) Co-immunoprecipitation (Co-IP) and immunoblot analysis showing the interaction of FLAG-tagged Gβ–associated complexes with components of the mTOR signaling pathway, including mTOR, and PI3K (p110) under rapamycin treatment and GiD activation for 30 mins. Data are presented as mean ± s.e.m. Statistical significance was determined using a two-tailed Student’s *t*-test.

**Figure S13: Pertussis toxin–mediated inhibition of Gi signaling modulates macrophage gene expression.** (A–F) Quantitative PCR analysis of macrophage polarization and inflammatory genes in control and LysM-PTX macrophages treated with forskolin (FRS). Expression of *Arg1*, *Atf4*, *Klf4*,*Il4*, *Nfκb*, *IL1α* was measured and normalized to 18s rRNA. (G) Plasma profiling of IL1β and TNFα concentration collected from WT and LysM-PTX mice kept on HFD. Data are presented as mean ± s.e.m. Statistical significance was determined using a two-tailed Student’s t-test.

**Figure S14: Genomic distribution and quality assessment of chromatin accessibility changes upon Gi inhibition.** (A) MA plot showing differential chromatin accessibility between LysM-PTX and WT macrophages, with regions exhibiting increased (red) or decreased (blue) accessibility. (B) Genomic distribution of regions showing increased or decreased accessibility, indicating enrichment of gained regions at promoter-proximal sites and decreased regions at distal intergenic loci. (C) Pathway enrichment analysis of genes associated with regions showing increased accessibility, highlighting signaling pathways related to cAMP–CREB, IL-6/STAT3, and metabolic regulation. (D) Motif enrichment analysis of regions with increased accessibility, showing enrichment of transcription factor binding motifs for CREB/ATF, NF-κB, and KLF family members. (E) Genome browser tracks showing chromatin accessibility at representative loci associated with the cAMP–CREB axis (Crebbp, Crtc1, Atf2). (F) Genome browser tracks showing chromatin accessibility at loci associated with regulatory and anti-inflammatory genes (Klf2, Nfkbia, Relb). Each condition was analyzed using two independent biological replicates, each comprising pooled samples from N = 3 mice.

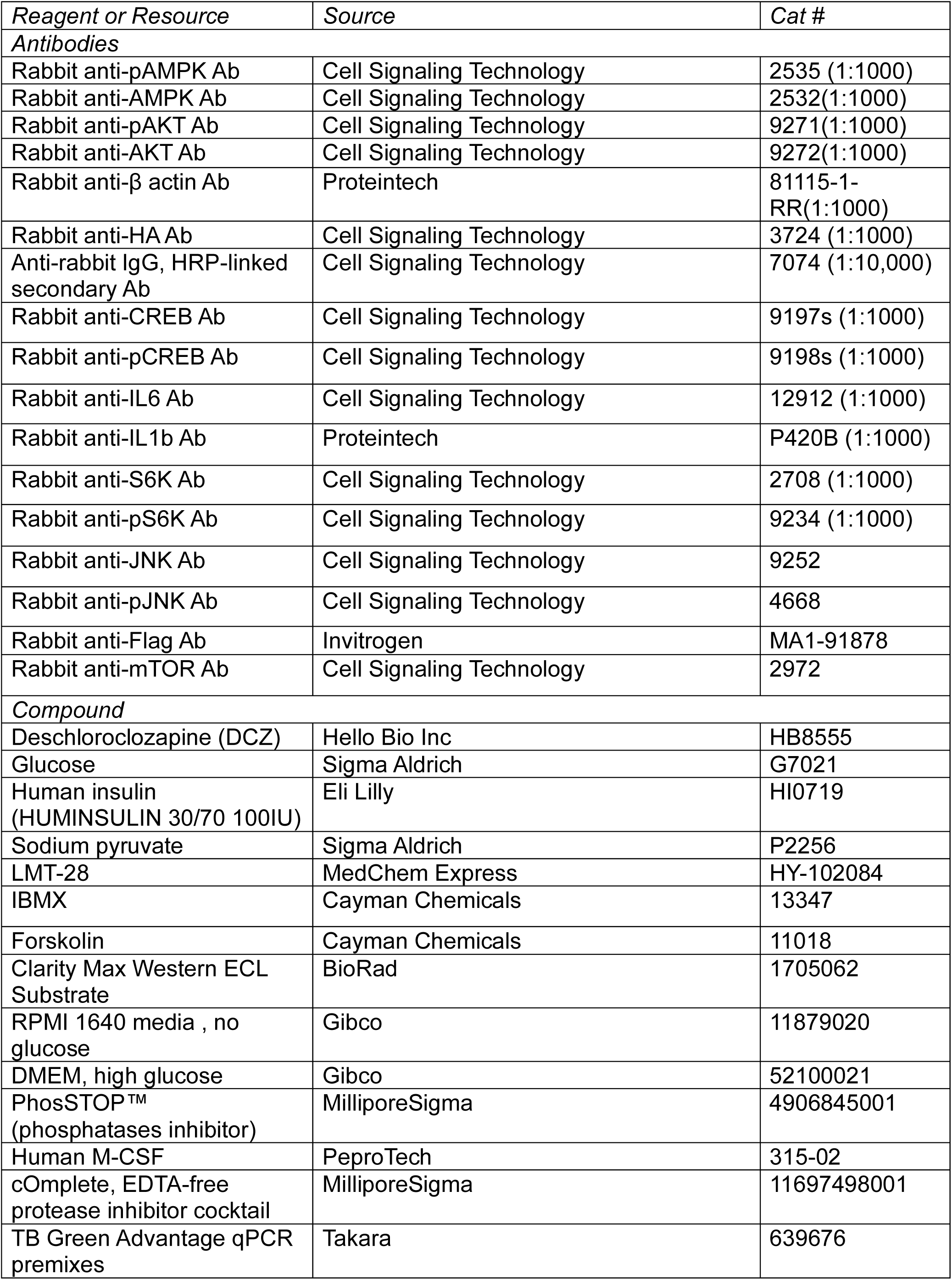
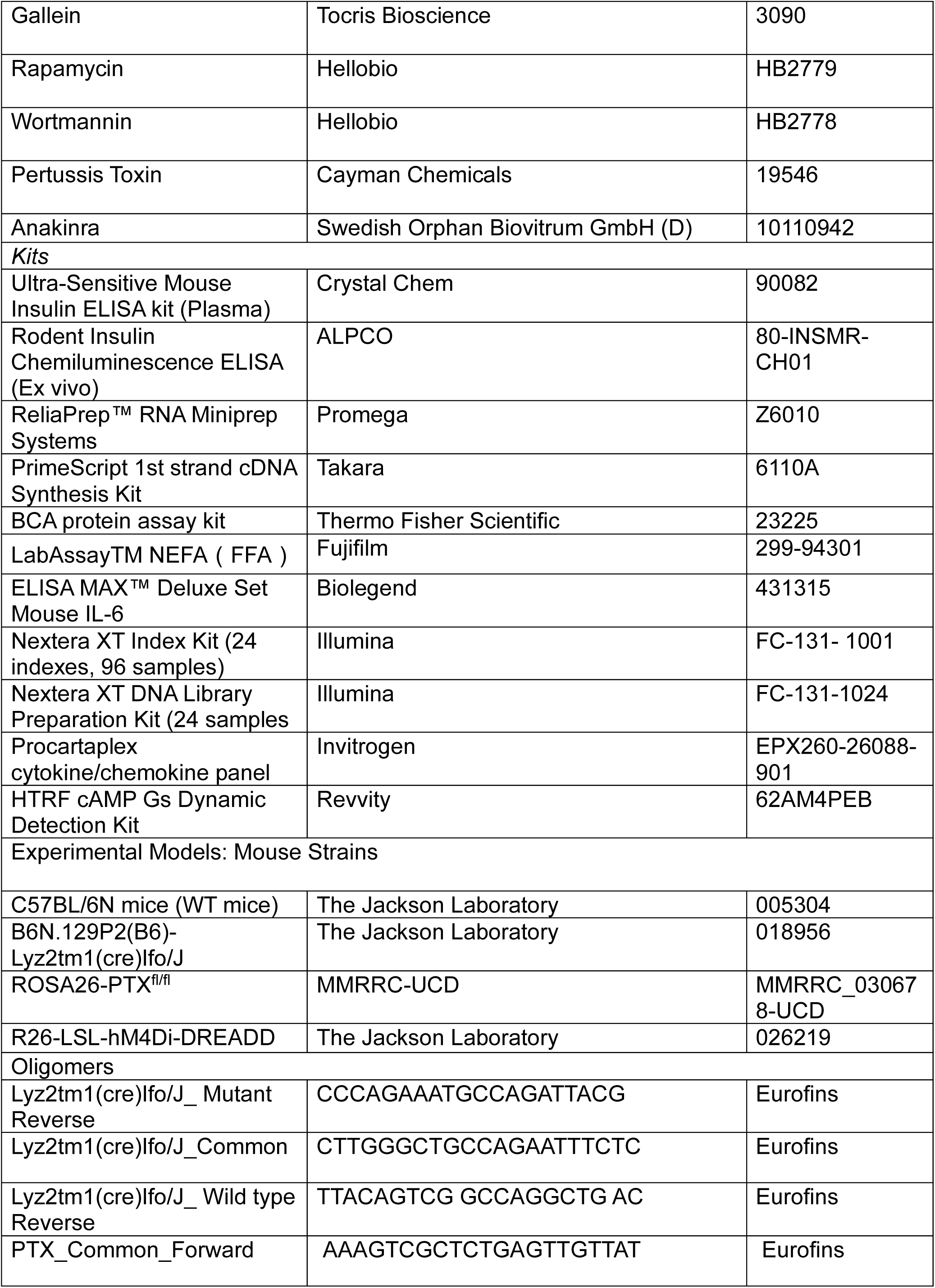
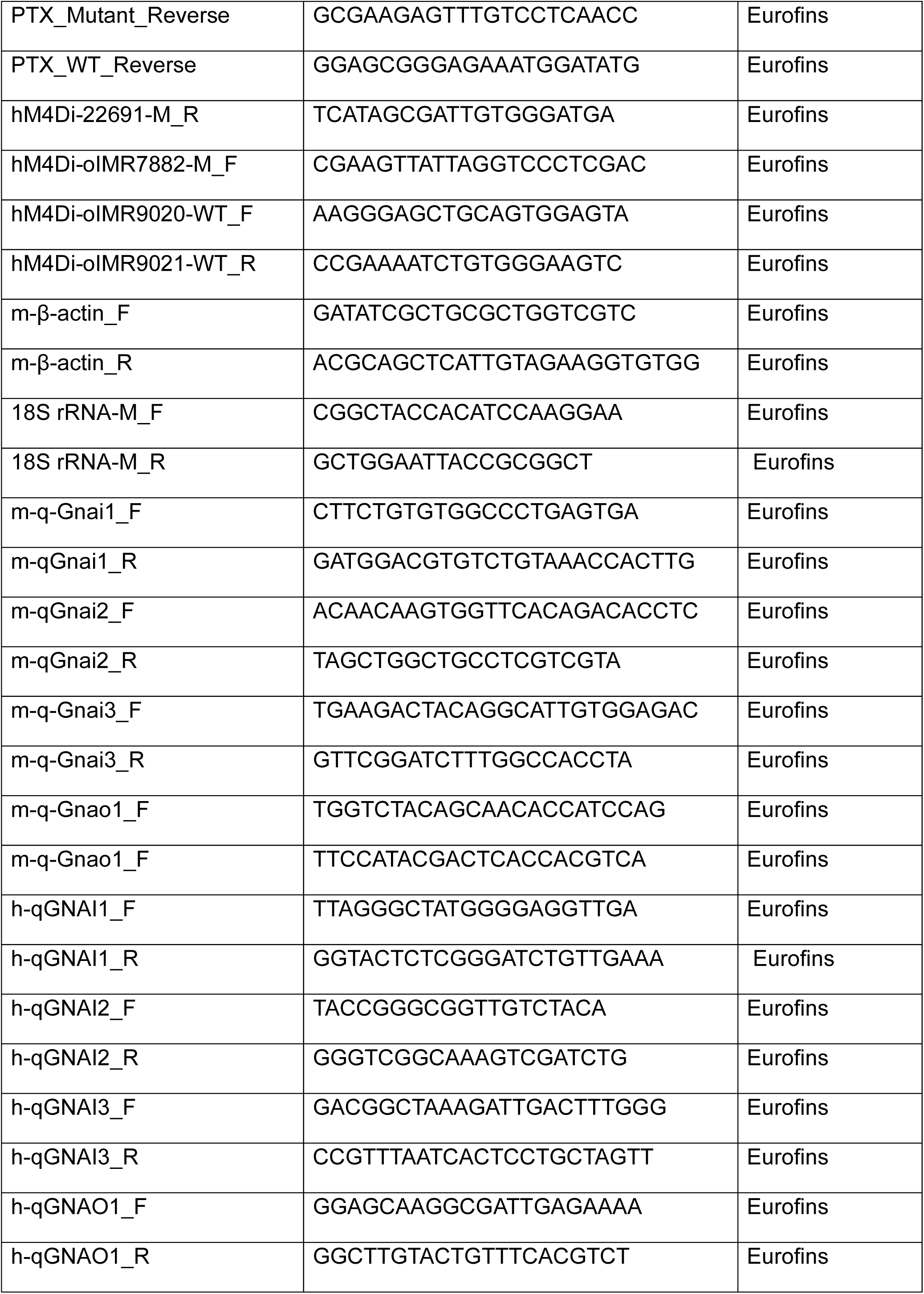
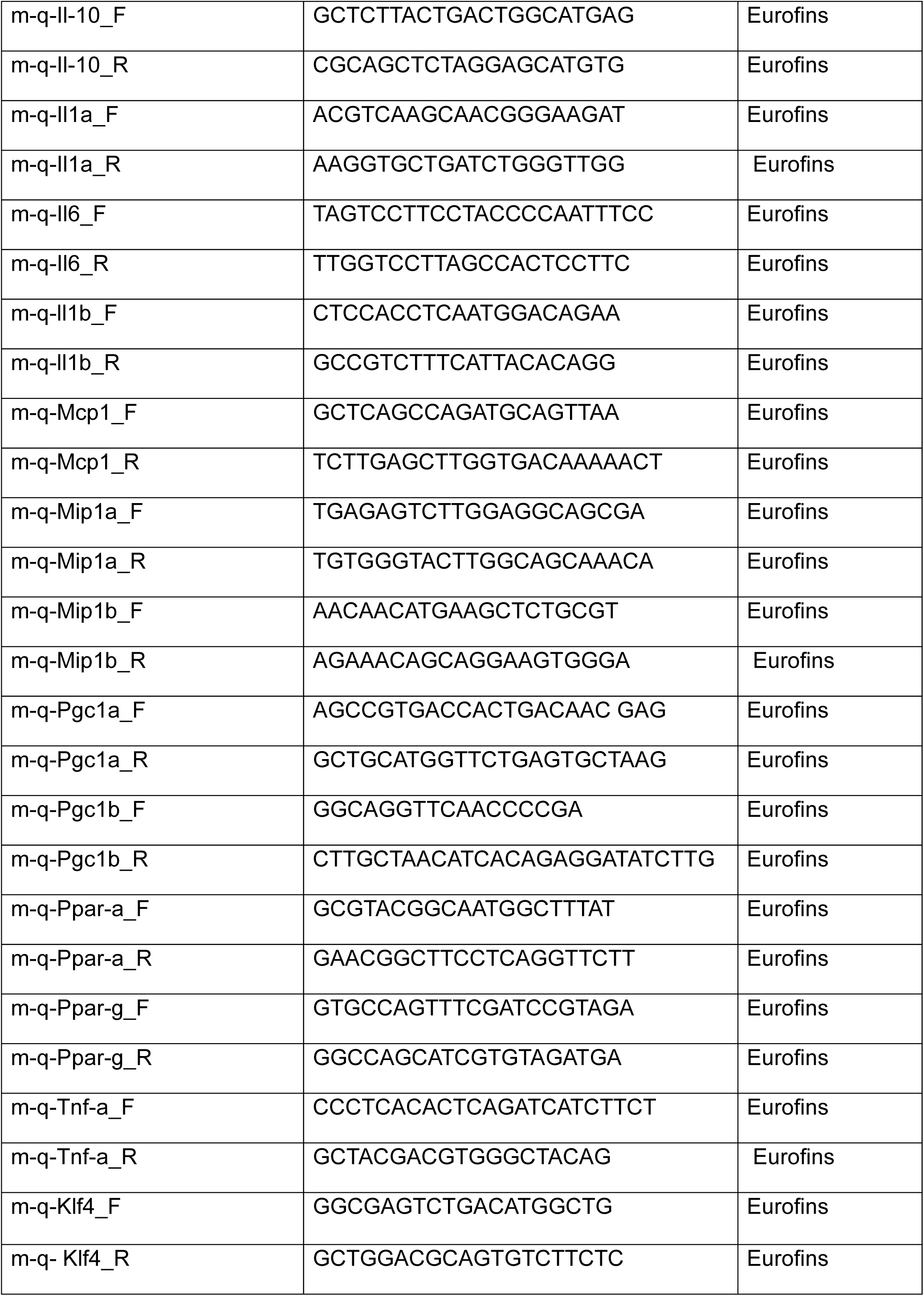
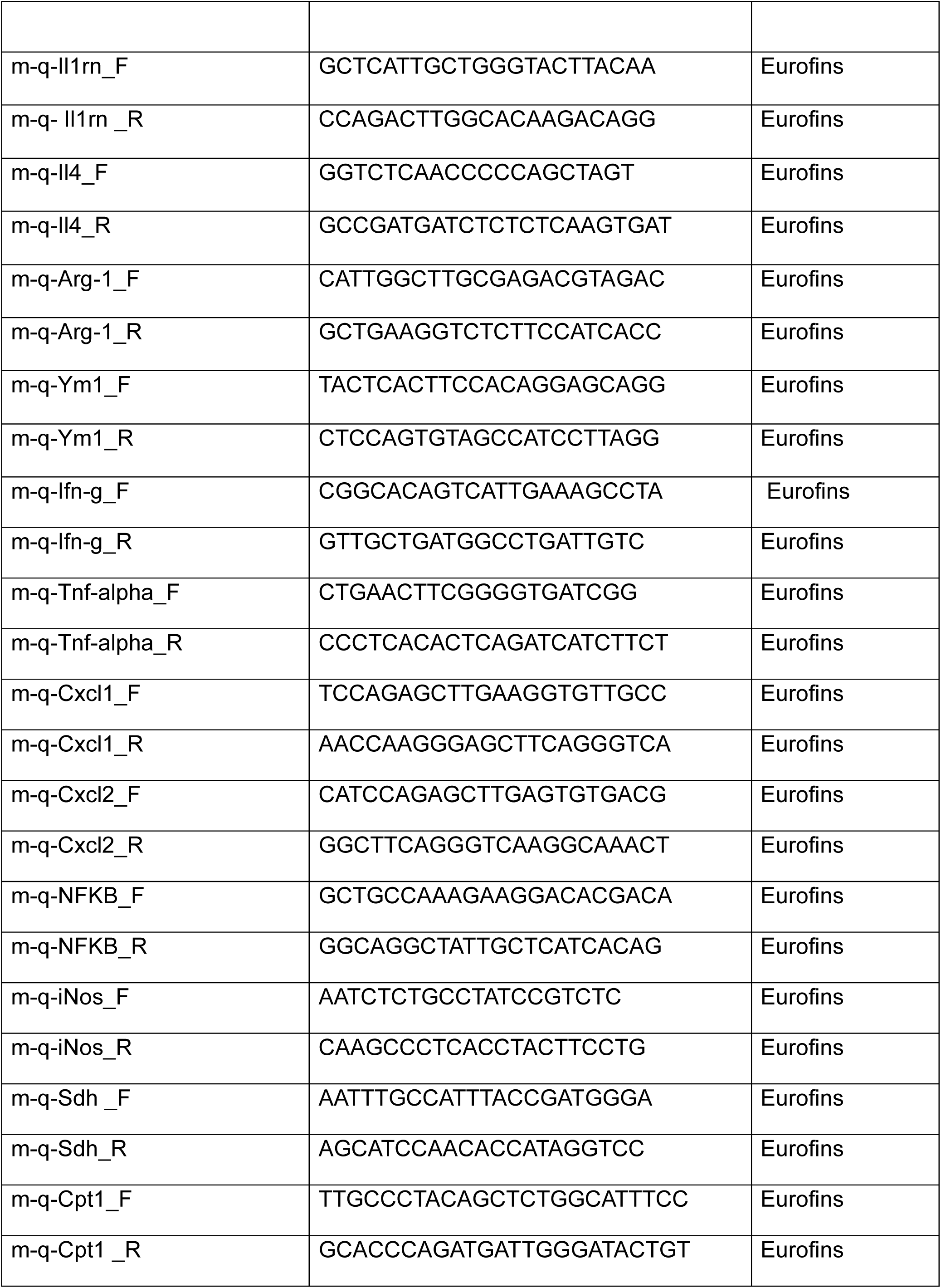
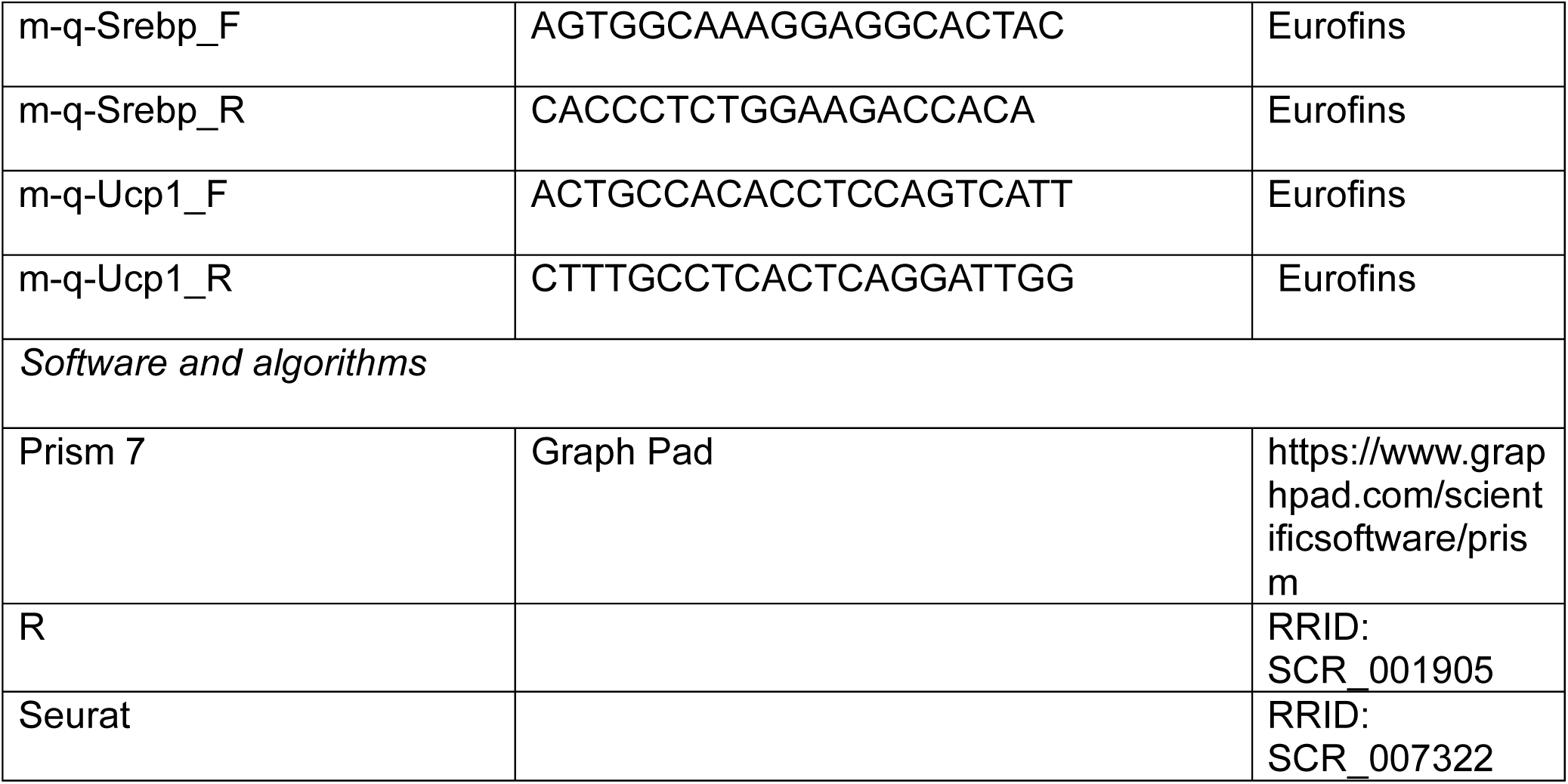
Resource Table: 1.

## References

1. Furman, D., et al., Chronic inflammation in the etiology of disease across the life span. Nat Med, 2019. 25(12): p. 1822–1832.

2. Rohm, T.V., et al., Inflammation in obesity, diabetes, and related disorders. Immunity, 2022. 55(1): p. 31–55.

3. Dandona, P., A. Aljada, and A. Bandyopadhyay, Inflammation: the link between insulin resistance, obesity and diabetes. Trends Immunol, 2004. 25(1): p. 4–7.

4. Smith, G.I., B. Mittendorfer, and S. Klein, Metabolically healthy obesity: facts and fantasies. J Clin Invest, 2019. 129(10): p. 3978–3989.

5. van Vliet-Ostaptchouk, J.V., et al., The prevalence of metabolic syndrome and metabolically healthy obesity in Europe: a collaborative analysis of ten large cohort studies. BMC Endocr Disord, 2014. 14: p. 9.

6. Caputo, T., F. Gilardi, and B. Desvergne, From chronic overnutrition to metaflammation and insulin resistance: adipose tissue and liver contributions. FEBS letters, 2017. 591(19): p. 3061–3088.

7. Wu, H. and C.M. Ballantyne, Metabolic Inflammation and Insulin Resistance in Obesity. Circulation Research, 2020. 126(11): p. 1549–1564.

8. Lu, J., et al., Adipose Tissue-Resident Immune Cells in Obesity and Type 2 Diabetes. Front Immunol, 2019. 10: p. 1173.

9. Gregor, M.F. and G.S. Hotamisligil, Inflammatory mechanisms in obesity. Annu Rev Immunol, 2011. 29: p. 415–45.

10. Lackey, D.E. and J.M. Olefsky, Regulation of metabolism by the innate immune system. Nature Reviews Endocrinology, 2016. 12(1): p. 15–28.

11. Hotamisligil, G.S., Inflammation, metaflammation and immunometabolic disorders. Nature, 2017. 542(7640): p. 177–185.

12. Weisberg, S.P., et al., Obesity is associated with macrophage accumulation in adipose tissue. J Clin Invest, 2003. 112(12): p. 1796–808.

13. Xu, H., et al., Chronic inflammation in fat plays a crucial role in the development of obesity-related insulin resistance. The Journal of clinical investigation, 2003. 112(12): p. 1821–1830.

14. Wellen, K.E. and G.S. Hotamisligil, Obesity-induced inflammatory changes in adipose tissue. The Journal of clinical investigation, 2003. 112(12): p. 1785–1788.

15. Ruggiero, A.D., C.C. Key, and K. Kavanagh, Adipose Tissue Macrophage Polarization in Healthy and Unhealthy Obesity. Front Nutr, 2021. 8: p. 625331.

16. Hata, M., et al., Past history of obesity triggers persistent epigenetic changes in innate immunity and exacerbates neuroinflammation. Science, 2023. 379(6627): p. 45–62.

17. Lumeng, C.N., et al., Increased Inflammatory Properties of Adipose Tissue Macrophages Recruited During Diet-Induced Obesity. Diabetes, 2007. 56(1): p. 16–23.

18. Strissel, K.J., et al., Adipocyte Death, Adipose Tissue Remodeling, and Obesity Complications. Diabetes, 2007. 56(12): p. 2910–2918.

19. Guria, S., et al., Adipose tissue macrophages and their role in obesity-associated insulin resistance: an overview of the complex dynamics at play. Biosci Rep, 2023. 43(3).

20. Tao, Y., Q. Jiang, and Q. Wang, Adipose tissue macrophages in remote modulation of hepatic glucose production. Front Immunol, 2022. 13: p. 998947.

21. Shoelson, S.E., J. Lee, and A.B. Goldfine, Inflammation and insulin resistance. The Journal of clinical investigation, 2006. 116(7): p. 1793–1801.

22. Alipourfard, I., N. Datukishvili, and D. Mikeladze, TNF-α Downregulation Modifies Insulin Receptor Substrate 1 (IRS-1) in Metabolic Signaling of Diabetic Insulin-Resistant Hepatocytes. Mediators Inflamm, 2019. 2019: p. 3560819.

23. Lin, H.H. and M. Stacey, G protein-coupled receptors in macrophages. Myeloid Cells in Health and Disease: A Synthesis, 2017: p. 485–505.

24. Weisberg, S.P., et al., CCR2 modulates inflammatory and metabolic effects of high-fat feeding. The Journal of clinical investigation, 2006. 116(1): p. 115–124.

25. Obstfeld, A.E., et al., CC chemokine receptor 2 (CCR2) regulates the hepatic recruitment of myeloid cells that promote obesity-induced hepatic steatosis. Diabetes, 2010. 59(4): p. 916–925.

26. Yokomizo, T., Leukotriene B4 receptors: novel roles in immunological regulations. Advances in enzyme regulation, 2011. 51(1): p. 59–64.

27. Li, P., et al., LTB4 promotes insulin resistance in obese mice by acting on macrophages, hepatocytes and myocytes. Nat Med, 2015. 21(3): p. 239–247.

28. Chen, X., et al., FFAR4-mediated IL-6 release from islet macrophages promotes insulin secretion and is compromised in type-2 diabetes. Nat Commun, 2025. 16(1): p. 3422.

29. Lipscomb, M., et al., Resolvin D2 limits atherosclerosis progression via myeloid cell-GPR18. FASEB J, 2024. 38(6): p. e23555.

30. Lattin, J., et al., G-protein-coupled receptor expression, function, and signaling in macrophages. J Leukoc Biol, 2007. 82(1): p. 16–32.

31. Zhu, H., et al., *Cre-dependent DREADD (designer receptors exclusively activated by designer drugs) mice*. genesis, 2016. 54(8): p. 439–446.

32. Armbruster, B.N., et al., Evolving the lock to fit the key to create a family of G protein-coupled receptors potently activated by an inert ligand. Proc Natl Acad Sci U S A, 2007. 104(12): p. 5163–8.

33. Wess, J., Use of Designer G Protein-Coupled Receptors to Dissect Metabolic Pathways. Trends Endocrinol Metab, 2016. 27(9): p. 600–603.

34. Rossi, M., et al., Hepatic Gi signaling regulates whole-body glucose homeostasis. J Clin Invest, 2018. 128(2): p. 746–759.

35. Regard, J.B., et al., Probing cell type-specific functions of Gi in vivo identifies GPCR regulators of insulin secretion. J Clin Invest, 2007. 117(12): p. 4034–43.

36. Shree, A., M.K. Pavan, and H. Zafar, scDREAMER for atlas-level integration of single-cell datasets using deep generative model paired with adversarial classifier. Nature Communications, 2023. 14(1): p. 7781.

37. Emont, M.P., et al., A single-cell atlas of human and mouse white adipose tissue. Nature, 2022. 603(7903): p. 926–933.

38. Walker, N.M., et al., *c-Jun N-terminal kinase (JNK)-mediated induction of mSin1 expression and mTORC2 activation in mesenchymal cells during fibrosis*. J Biol Chem, 2018. 293(44): p. 17229–17239.

39. Kitanaka, T., et al., JNK activation is essential for activation of MEK/ERK signaling in IL-1beta-induced COX-2 expression in synovial fibroblasts. Sci Rep, 2017. 7: p. 39914.

40. Chen, C.-L., et al., Molecular basis for Gβγ-mediated activation of phosphoinositide 3-kinase γ. Nature Structural & Molecular Biology, 2024. 31(8): p. 1198–1207.

41. Wiedemann, S.J., et al., The cephalic phase of insulin release is modulated by IL-1β. Cell metabolism, 2022. 34(7): p. 991–1003. e6.

42. Montminy, M., Transcriptional regulation by cyclic AMP. Annu Rev Biochem, 1997. 66: p. 807–22.

43. Wen, A.Y., K.M. Sakamoto, and L.S. Miller, The role of the transcription factor CREB in immune function. J Immunol, 2010. 185(11): p. 6413–9.

44. Sarvas, J.L., N. Khaper, and S.J. Lees, The IL-6 Paradox: Context Dependent Interplay of SOCS3 and AMPK. J Diabetes Metab, 2013. **Suppl** 13.

45. Heinrich, P.C., et al., Interleukin-6-type cytokine signalling through the gp130/Jak/STAT pathway. Biochem J, 1998. 334 (**Pt 2**)(Pt 2): p. 297–314.

46. Kelly, M., et al., Activation of AMP-activated protein kinase by interleukin-6 in rat skeletal muscle: association with changes in cAMP, energy state, and endogenous fuel mobilization. Diabetes, 2009. 58(9): p. 1953–60.

47. Priceman, S.J., et al., Regulation of adipose tissue T cell subsets by Stat3 is crucial for diet-induced obesity and insulin resistance. Proc Natl Acad Sci U S A, 2013. 110(32): p. 13079–84.

48. Huang, T., et al., Adipocyte-derived kynurenine promotes obesity and insulin resistance by activating the AhR/STAT3/IL-6 signaling. Nat Commun, 2022. 13(1): p. 3489.

49. Sims, E.A., Are there persons who are obese, but metabolically healthy? Metabolism, 2001. 50(12): p. 1499–504.

50. Samocha-Bonet, D., et al., Insulin-sensitive obesity in humans – a ‘favorable fat’ phenotype? Trends Endocrinol Metab, 2012. 23(3): p. 116–24.

51. Kloting, N., et al., Insulin-sensitive obesity. Am J Physiol Endocrinol Metab, 2010. 299(3): p. E506–15.

52. Bluher, M., Metabolically Healthy Obesity. Endocr Rev, 2020. 41(3).

53. Duque, A.P., et al., Emerging concepts in metabolically healthy obesity. Am J Cardiovasc Dis, 2020. 10(2): p. 48–61.

54. Ying, W., et al., The role of macrophages in obesity-associated islet inflammation and beta-cell abnormalities. Nat Rev Endocrinol, 2020. 16(2): p. 81–90.

55. Lumeng, C.N., J.L. Bodzin, and A.R. Saltiel, Obesity induces a phenotypic switch in adipose tissue macrophage polarization. J Clin Invest, 2007. 117(1): p. 175–84.

56. Trauelsen, M., et al., Extracellular succinate hyperpolarizes M2 macrophages through SUCNR1/GPR91-mediated Gq signaling. Cell Reports, 2021. 35(11).

57. Lanahan, S.M., M.P. Wymann, and C.L. Lucas, The role of PI3Kγ in the immune system: new insights and translational implications. Nature Reviews Immunology, 2022. 22(11): p. 687–700.

58. Hirsch, E., et al., Central role for G protein-coupled phosphoinositide 3-kinase gamma in inflammation. Science, 2000. 287(5455): p. 1049–53.

59. Stephens, L.R., et al., The G beta gamma sensitivity of a PI3K is dependent upon a tightly associated adaptor, p101. Cell, 1997. 89(1): p. 105–14.

60. Aguirre, V., et al., The c-Jun NH(2)-terminal kinase promotes insulin resistance during association with insulin receptor substrate-1 and phosphorylation of Ser(307). J Biol Chem, 2000. 275(12): p. 9047–54.

61. Solinas, G., et al., JNK1 in hematopoietically derived cells contributes to diet-induced inflammation and insulin resistance without affecting obesity. Cell Metab, 2007. 6(5): p. 386–97.

62. Jager, J., et al., Interleukin-1beta-induced insulin resistance in adipocytes through down-regulation of insulin receptor substrate-1 expression. Endocrinology, 2007. 148(1): p. 241–51.

63. Wen, H., et al., Fatty acid-induced NLRP3-ASC inflammasome activation interferes with insulin signaling. Nat Immunol, 2011. 12(5): p. 408–15.

64. Maedler, K., et al., Glucose-induced beta cell production of IL-1beta contributes to glucotoxicity in human pancreatic islets. J Clin Invest, 2002. 110(6): p. 851–60.

65. Dror, E., et al., Postprandial macrophage-derived IL-1beta stimulates insulin, and both synergistically promote glucose disposal and inflammation. Nat Immunol, 2017. 18(3): p. 283–292.

66. Ridker, P.M., et al., Antiinflammatory Therapy with Canakinumab for Atherosclerotic Disease. N Engl J Med, 2017. 377(12): p. 1119–1131.

67. Samiea, A., et al., Interleukin-10 contributes to PGE2 signalling through upregulation of EP4 via SHIP1 and STAT3. PLoS One, 2020. 15(4): p. e0230427.

68. Tavares, L.P., et al., Blame the signaling: Role of cAMP for the resolution of inflammation. Pharmacol Res, 2020. 159: p. 105030.

69. Klover, P.J., et al., Chronic exposure to interleukin-6 causes hepatic insulin resistance in mice. Diabetes, 2003. 52(11): p. 2784–9.

70. Mauer, J., et al., Signaling by IL-6 promotes alternative activation of macrophages to limit endotoxemia and obesity-associated resistance to insulin. Nat Immunol, 2014. 15(5): p. 423–30.

71. Han, M.S., et al., Regulation of adipose tissue inflammation by interleukin 6. Proc Natl Acad Sci U S A, 2020. 117(6): p. 2751–2760.

72. Wu, L., et al., AMP-Activated Protein Kinase (AMPK) Regulates Energy Metabolism through Modulating Thermogenesis in Adipose Tissue. Frontiers in Physiology, 2018. **Volume** 9 **-** 2018.

73. Mottillo, Emilio P., et al., Lack of Adipocyte AMPK Exacerbates Insulin Resistance and Hepatic Steatosis through Brown and Beige Adipose Tissue Function. Cell Metabolism, 2016. 24(1): p. 118–129.

74. O’Neill, H.M., AMPK and Exercise: Glucose Uptake and Insulin Sensitivity. Diabetes Metab J, 2013. 37(1): p. 1–21.

75. Steneberg, P., et al., PAN-AMPK activator O304 improves glucose homeostasis and microvascular perfusion in mice and type 2 diabetes patients. JCI Insight, 2018. 3(12).

76. Rose-John, S., IL-6 trans-signaling via the soluble IL-6 receptor: importance for the pro-inflammatory activities of IL-6. Int J Biol Sci, 2012. 8(9): p. 1237–47.

77. Rose-John, S., Interleukin-6 signalling in health and disease. F1000Res, 2020. 9.

78. Wang, L., et al., Adipocyte G(i) signaling is essential for maintaining whole-body glucose homeostasis and insulin sensitivity. Nat Commun, 2020. 11(1): p. 2995.

79. Oteng, A.B., et al., Activation of Gs signaling in mouse enteroendocrine K cells greatly improves obesity– and diabetes-related metabolic deficits. J Clin Invest, 2024. 134(24).

80. Wiedemann, S.J., et al., The cephalic phase of insulin release is modulated by IL-1beta. Cell Metab, 2022. 34(7): p. 991–1003 e6.

81. Romera-Hernandez, M., et al., Yap1-driven intestinal repair is controlled by group 3 innate lymphoid cells. Cell Reports, 2020. 30(1): p. 37–45. e3.

82. Toda, G., et al., Preparation and culture of bone marrow-derived macrophages from mice for functional analysis. STAR Protoc, 2021. 2(1): p. 100246.

83. Sarbassov, D.D., et al., Prolonged Rapamycin Treatment Inhibits mTORC2 Assembly and Akt/PKB. Molecular Cell, 2006. 22(2): p. 159–168.

84. Liu, Y., et al., Wortmannin, a Widely Used Phosphoinositide 3-Kinase Inhibitor, also Potently Inhibits Mammalian Polo-like Kinase. Chemistry & Biology, 2005. 12(1): p. 99–107.

85. Lehmann, D.M., A.M. Seneviratne, and A.V. Smrcka, Small molecule disruption of G protein beta gamma subunit signaling inhibits neutrophil chemotaxis and inflammation. Mol Pharmacol, 2008. 73(2): p. 410–8.

86. Negoita, F., et al., CaMKK2 is not involved in contraction-stimulated AMPK activation and glucose uptake in skeletal muscle. Molecular Metabolism, 2023. 75: p. 101761.

87. Kim, D., et al., Graph-based genome alignment and genotyping with HISAT2 and HISAT-genotype. Nat Biotechnol, 2019. 37(8): p. 907–915.

88. Liao, Y., G.K. Smyth, and W. Shi, featureCounts: an efficient general purpose program for assigning sequence reads to genomic features. Bioinformatics, 2014. 30(7): p. 923–30.

89. Love, M.I., W. Huber, and S. Anders, Moderated estimation of fold change and dispersion for RNA-seq data with DESeq2. Genome Biol, 2014. 15(12): p. 550.

90. Stuart, T., et al., Comprehensive Integration of Single-Cell Data. Cell, 2019. 177(7): p. 1888–1902 e21.

91. Pydi, S.P., et al., Adipocyte β-arrestin-2 is essential for maintaining whole body glucose and energy homeostasis. Nature Communications, 2019. 10(1): p. 2936.

92. Liu, Y., et al., miR-324-5p inhibits C2C12 cell differentiation and promotes intramuscular lipid deposition through lncDUM and PM20D1. Molecular Therapy Nucleic Acids, 2020. 22: p. 722–732.

93. Cui, C., et al., Isolation of polymorphonuclear neutrophils and monocytes from a single sample of human peripheral blood. STAR Protoc, 2021. 2(4): p. 100845.

