## Supplementary Figures for "Myeloid Gi signaling acts as a weight-independent immunometabolic switch controlling systemic insulin sensitivity"

### Fig. S1

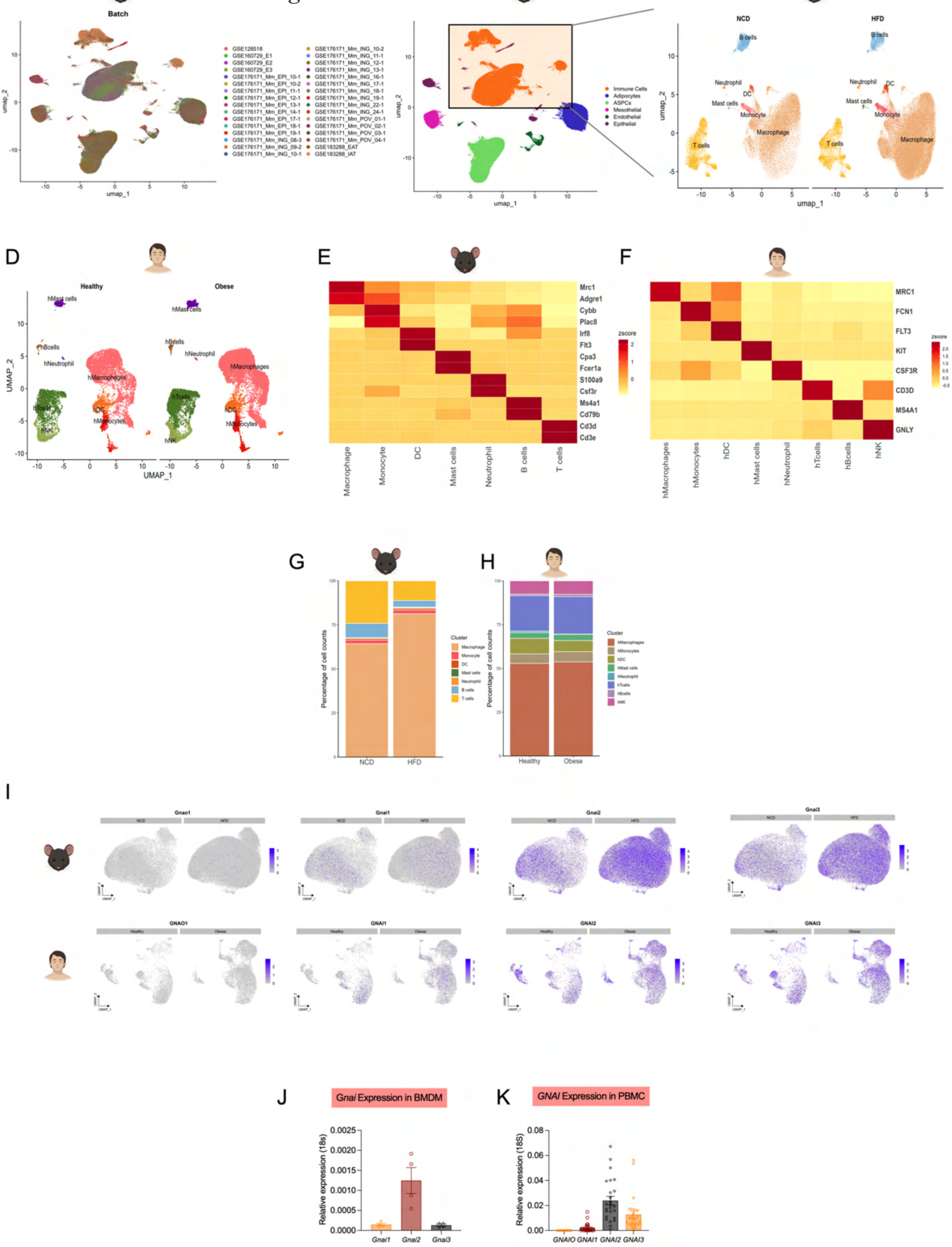

A
Batch
B
Immune Cells
Adipocytes
ASPCs
Mesothelial
Endothelial
Epithelial
C
NCD
HFD
B cells
Neutrophil
DC
Mast cells
Monocyte
Macrophage
T cells
D
Healthy
Obese
hMast cells
hBcells
hNeutrophil
hMacrophages
hTcells
hDC
hMonocytes
hNK
E
Mrc1
Adgre1
Cybb
Plac8
Irf8
Flt3
Cpa3
Fcer1a
S100a9
Csf3r
Ms4a1
Cd79b
Cd3d
Cd3e
zscore
Macrophage
Monocyte
DC
Mast cells
Neutrophil
B cells
T cells
F
MRC1
FCN1
FLT3
KIT
CSF3R
CD3D
MS4A1
GNLY
hMacrophages
hMonocytes
hDC
hMast cells
hNeutrophil
hTcells
hBcells
hNK
G
Percentage of cell counts
NCD
HFD
Cluster
H
Healthy
Obese
I
Gnao1
Gnai1
Gnai2
Gnai3
GNAO1
GNAI1
GNAI2
GNAI3
J
*Gnai* Expression in BMDM
Relative expression (18s)
*Gnai1*
*Gnai2*
*Gnai3*
K
*GNAI* Expression in PBMC
Relative expression (18S)
*GNAIO*
*GNAI1*
*GNAI2*
*GNAI3*

#### Fig. S2

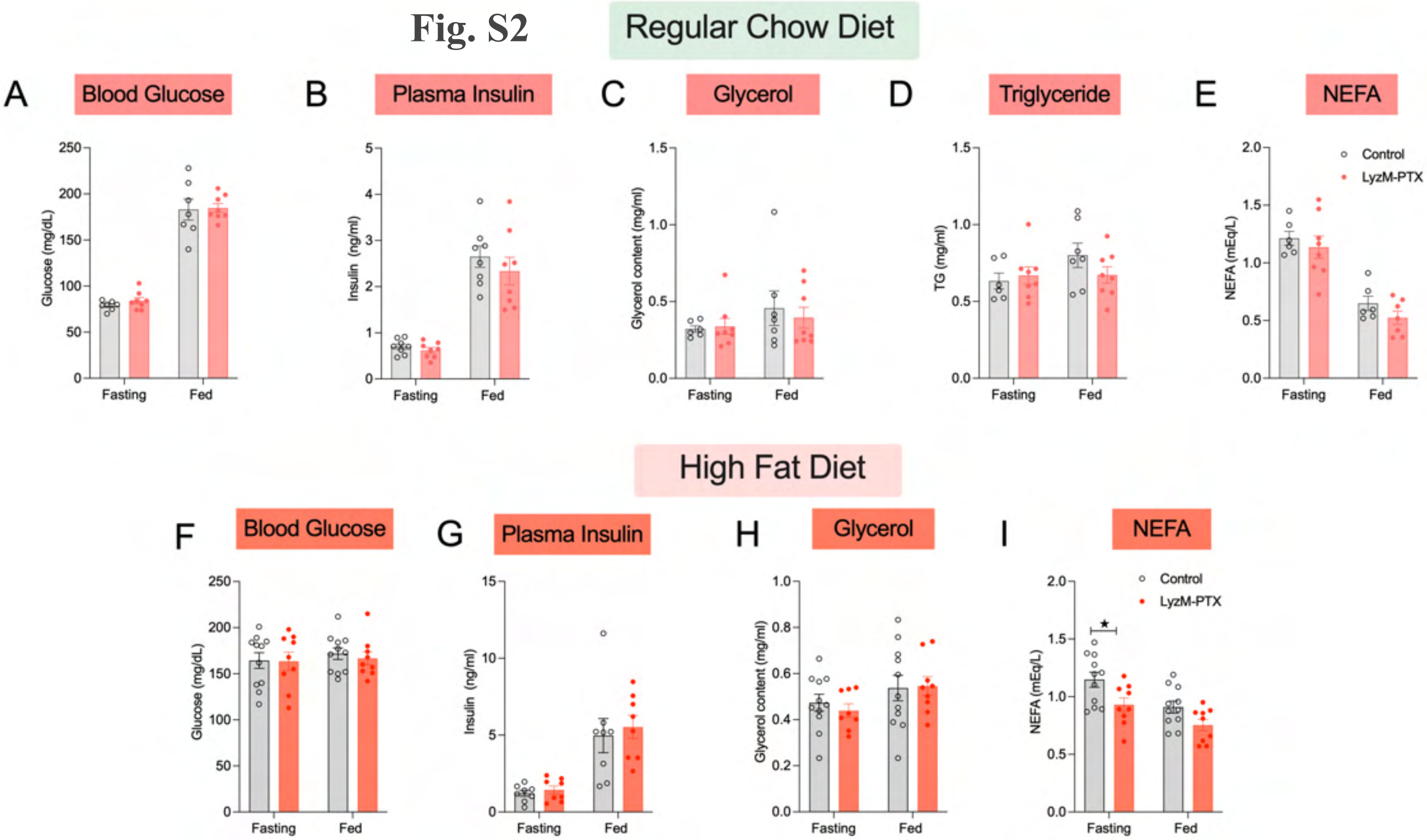

Regular Chow Diet
A Blood Glucose
Glucose (mg/dL)
250
200
150
100
50
0
Fasting
Fed
B Plasma Insulin
Insulin (ng/ml)
5
4
3
2
1
0
Fasting
Fed
C Glycerol
Glycerol content (mg/ml)
1.5
1.0
0.5
0.0
Fasting
Fed
D Triglyceride
TG (mg/ml)
1.5
1.0
0.5
0.0
Fasting
Fed
E NEFA
NEFA (mEq/L)
2.0
1.5
1.0
0.5
0.0
Fasting
Fed
Control
LyzM-PTX
High Fat Diet
F Blood Glucose
Glucose (mg/dL)
250
200
150
100
50
0
Fasting
Fed
G Plasma Insulin
Insulin (ng/ml)
15
10
5
0
Fasting
Fed
H Glycerol
Glycerol content (mg/ml)
1.0
0.8
0.6
0.4
0.2
0.0
Fasting
Fed
I NEFA
NEFA (mEq/L)
2.0
1.5
1.0
0.5
0.0
Fasting
Fed
Control
LyzM-PTX
★

Fig. S3

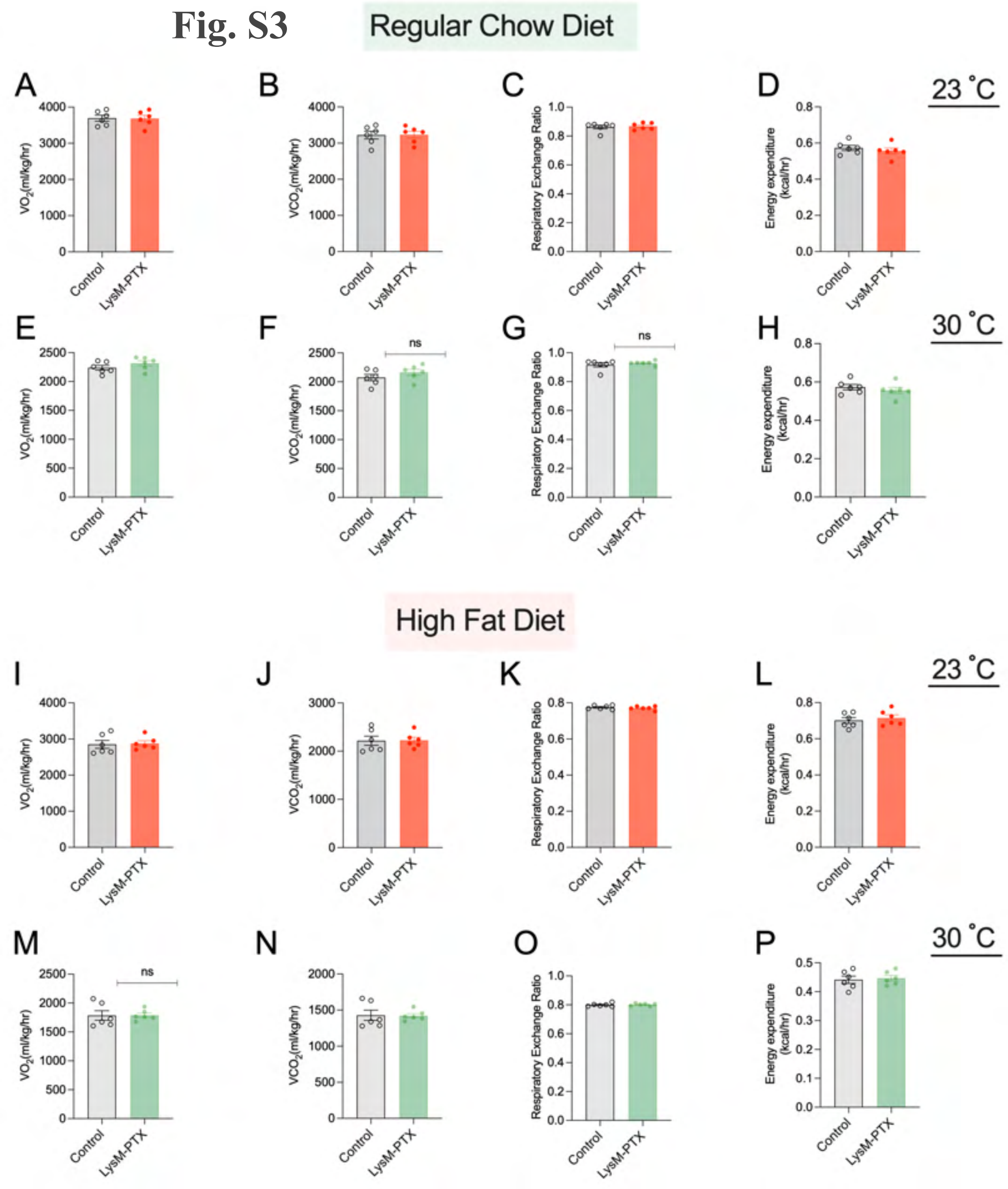
Regular Chow Diet
23 ˚C
30 ˚C
High Fat Diet
23 ˚C
30 ˚C
A
B
C
D
E
F
G
H
I
J
K
L
M
N
O
P
VO2(ml/kg/hr)
VCO2(ml/kg/hr)
Respiratory Exchange Ratio
Energy expenditure (kcal/hr)
Control
LysM-PTX
ns

### Fig. S4

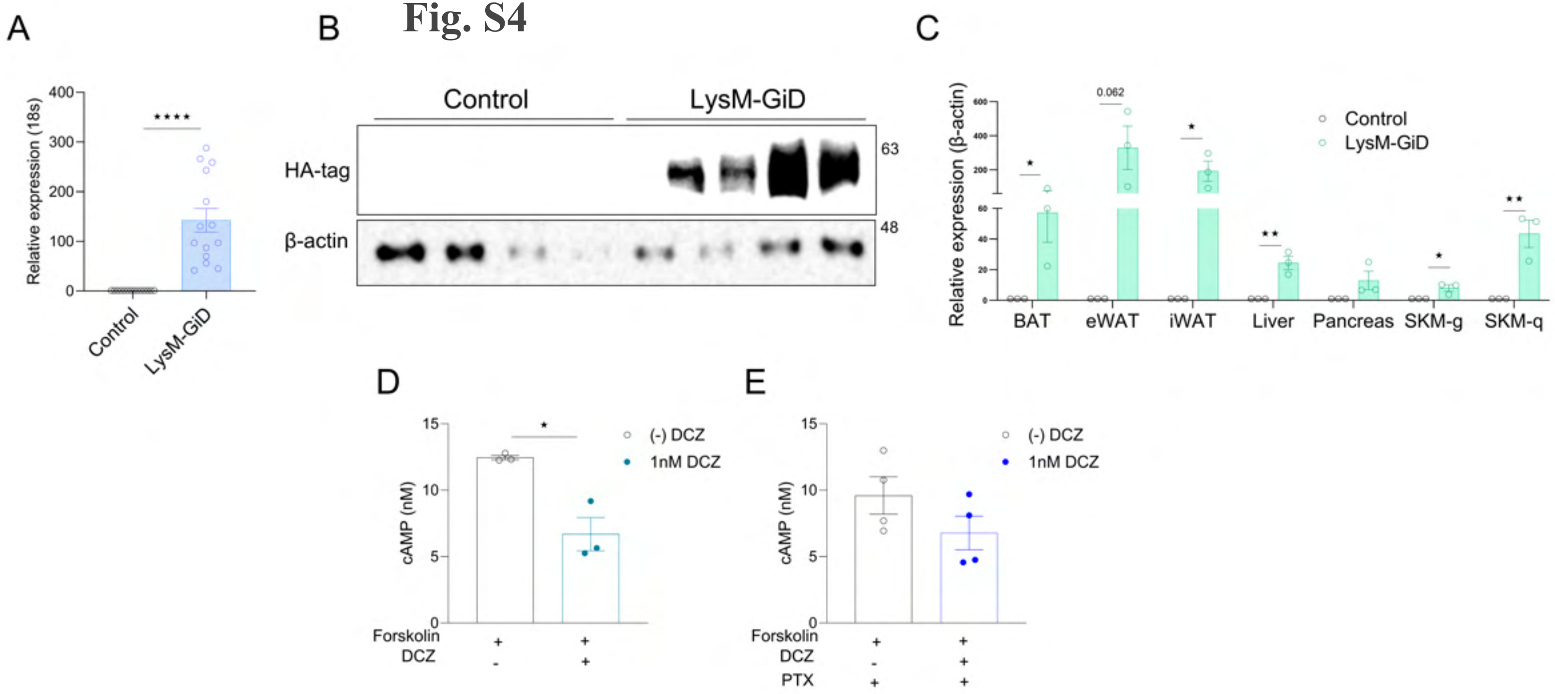

A
B
C
D
E
Relative expression (18s)
****
Control
LysM-GiD
HA-tag
β-actin
63
48
Relative expression (β-actin)
0.062
BAT
eWAT
iWAT
Liver
Pancreas
SKM-g
SKM-q
cAMP (nM)
(-) DCZ
1nM DCZ
Forskolin
DCZ
PTX

Fig. S5

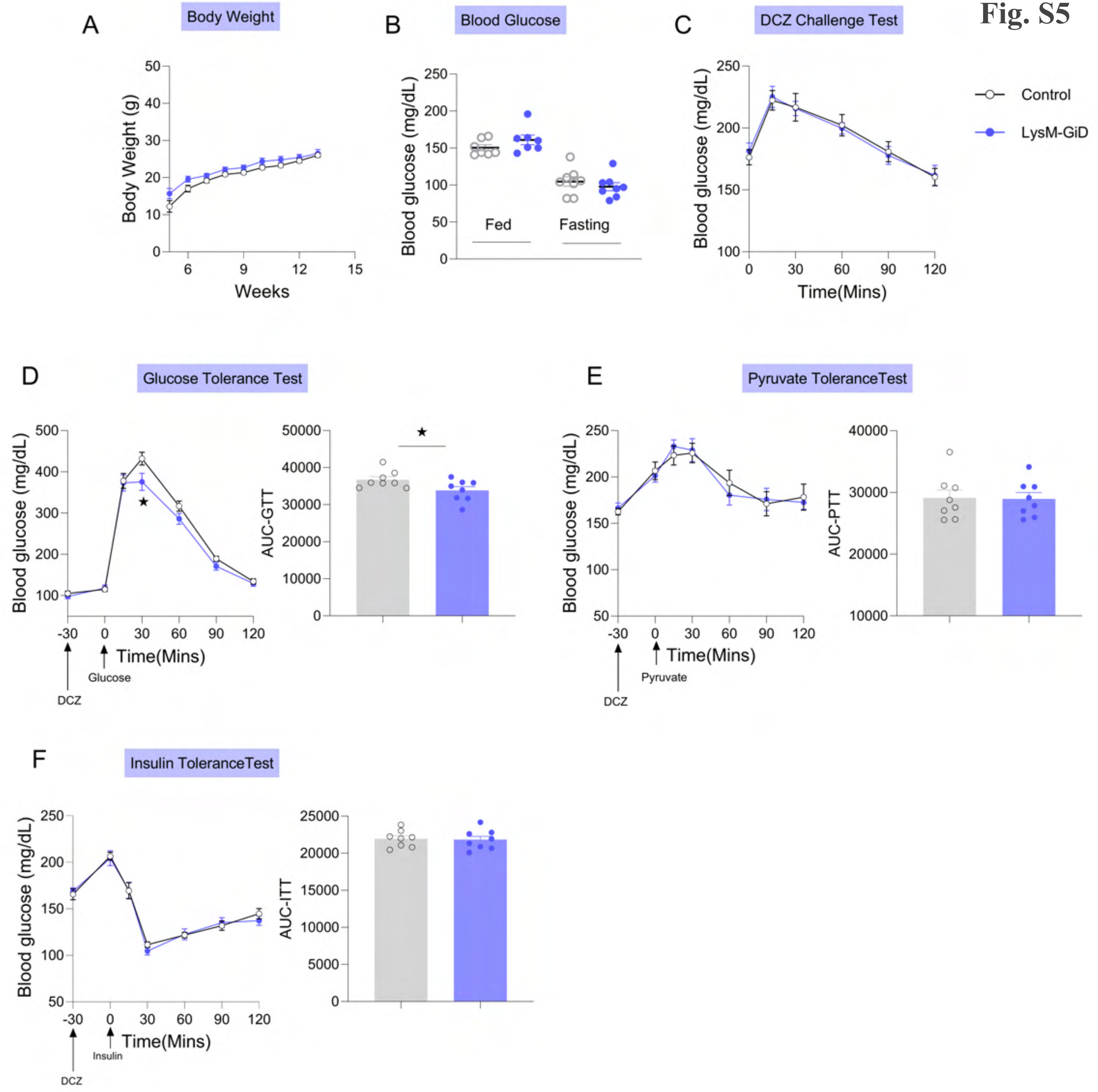

A
Body Weight
Body Weight (g)
Weeks
B
Blood Glucose
Blood glucose (mg/dL)
Fed
Fasting
C
DCZ Challenge Test
Blood glucose (mg/dL)
Time(Mins)
Control
LysM-GiD
D
Glucose Tolerance Test
Blood glucose (mg/dL)
Time(Mins)
Glucose
DCZ
AUC-GTT
E
Pyruvate ToleranceTest
Blood glucose (mg/dL)
Time(Mins)
Pyruvate
DCZ
AUC-PTT
F
Insulin ToleranceTest
Blood glucose (mg/dL)
Time(Mins)
Insulin
DCZ
AUC-ITT

### Fig. S6

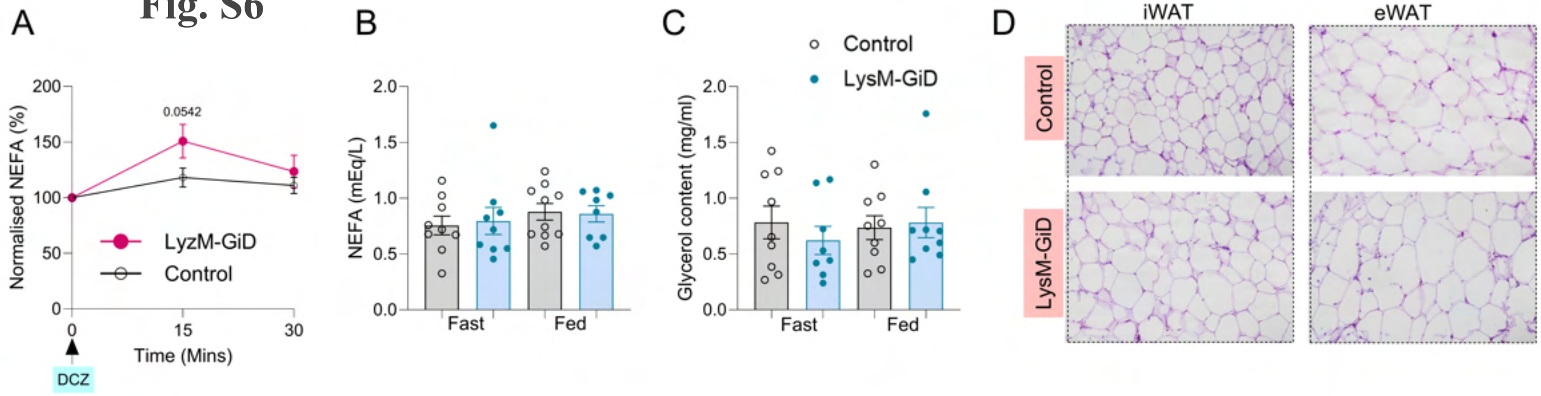
A
Normalised NEFA (%)
200
150
100
50
0
0.0542
LyzM-GiD
Control
0
15
30
Time (Mins)
DCZ
B
NEFA (mEq/L)
2.0
1.5
1.0
0.5
0.0
Fast
Fed
C
Control
LysM-GiD
Glycerol content (mg/ml)
2.0
1.5
1.0
0.5
0.0
Fast
Fed
D
iWAT
eWAT
Control
LysM-GiD

**Fig. S7**

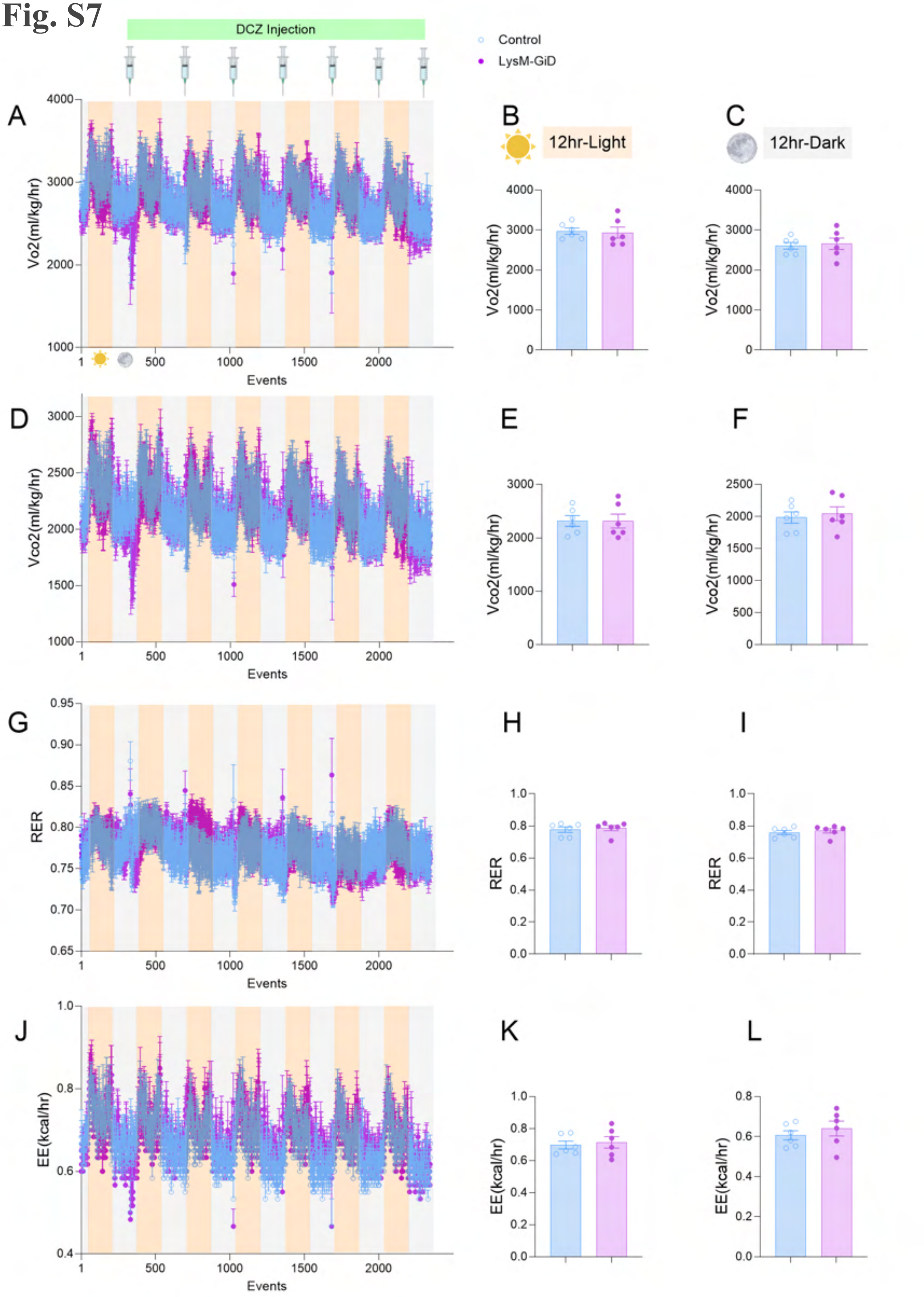
DCZ Injection
Control
LysM-GiD
A
Vo2(ml/kg/hr)
4000
3000
2000
1000
1
500
1000
1500
2000
Events
B
12hr-Light
C
12hr-Dark
D
Vco2(ml/kg/hr)
E
F
G
RER
H
I
J
EE(kcal/hr)
K
L

### Fig. S8

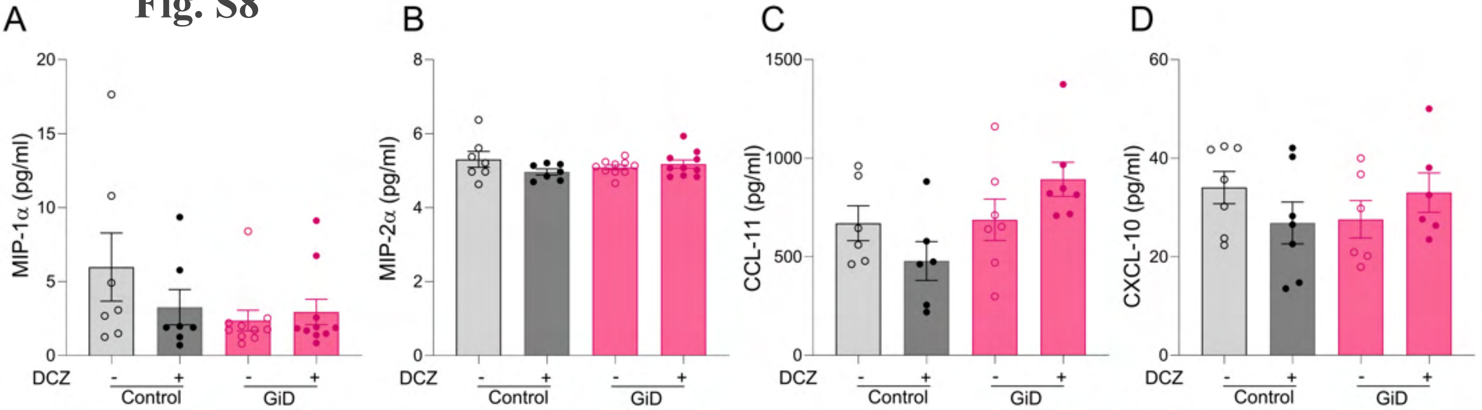
A
B
C
D
MIP-1α (pg/ml)
MIP-2α (pg/ml)
CCL-11 (pg/ml)
CXCL-10 (pg/ml)
0
5
10
15
20
2
4
6
8
500
1000
1500
40
60
DCZ
-
+
Control
GiD

**Fig. S9**

A

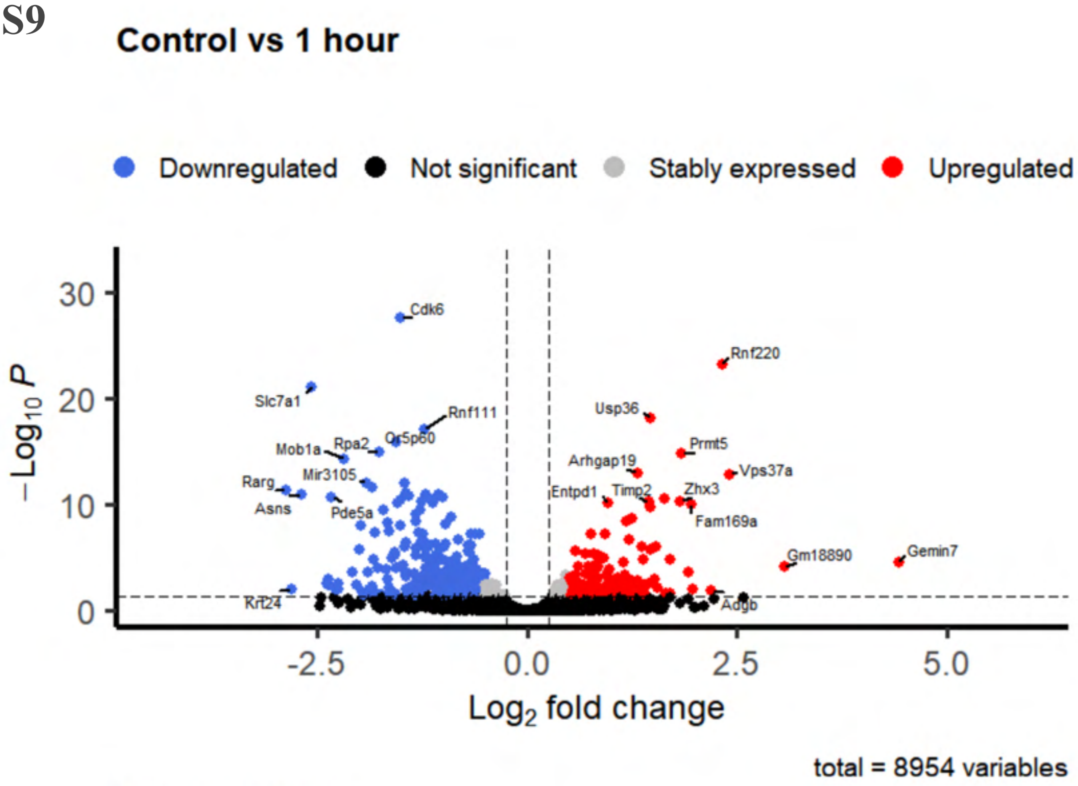

Control vs 1 hour
Downregulated
Not significant
Stably expressed
Upregulated
−Log10 P
Log2 fold change
total = 8954 variables
Cdk6
Rnf220
Slc7a1
Usp36
Rnf111
Mob1a
Rpa2
Or5p60
Prmt5
Arhgap19
Vps37a
Mir3105
Rarg
Asns
Pde5a
Entpd1
Timp2
Zhx3
Fam169a
Gm18890
Gemin7
Krt24
Adgb


B

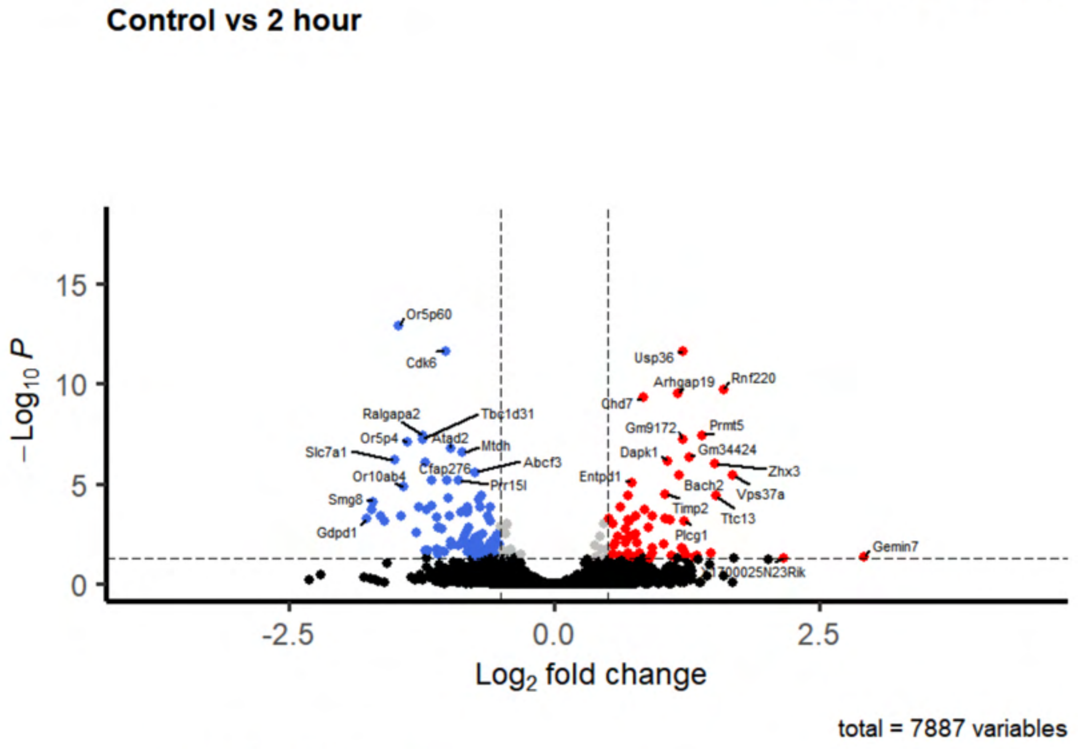

Control vs 2 hour
−Log10 P
Log2 fold change
total = 7887 variables
Or5p60
Cdk6
Usp36
Arhgap19
Rnf220
Chd7
Ralgapa2
Tbc1d31
Or5p4
Atad2
Mtdh
Slc7a1
Gm9172
Prmt5
Dapk1
Gm34424
Abcf3
Cfap276
Zhx3
Or10ab4
Entpd1
Prr15l
Bach2
Vps37a
Smg8
Timp2
Ttc13
Gdpd1
Plcg1
Gemin7
1700025N23Rik


C

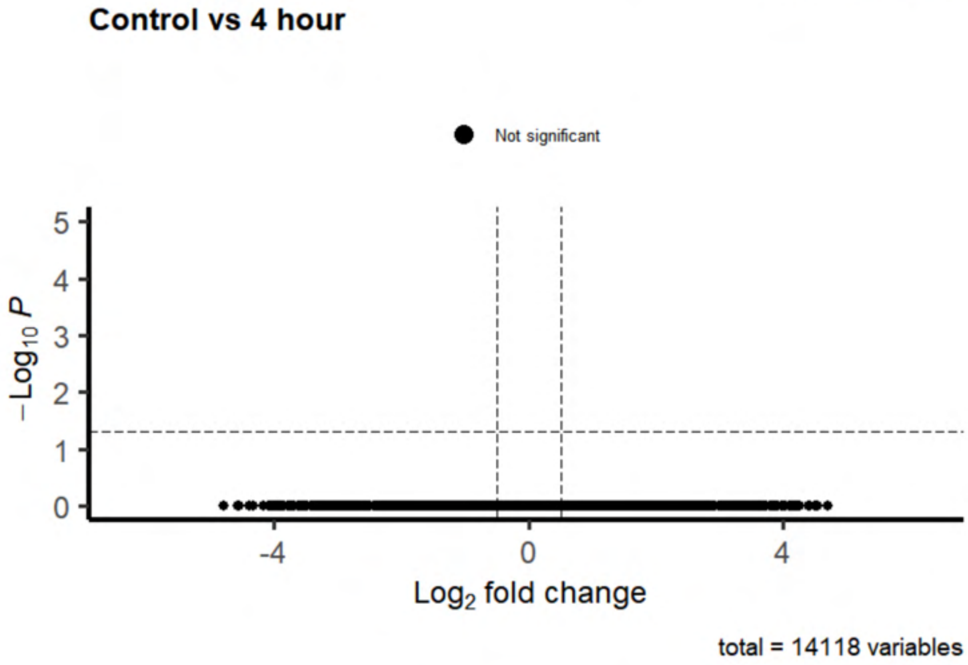

Control vs 4 hour
Not significant
−Log10 P
Log2 fold change
total = 14118 variables

### Fig. S10

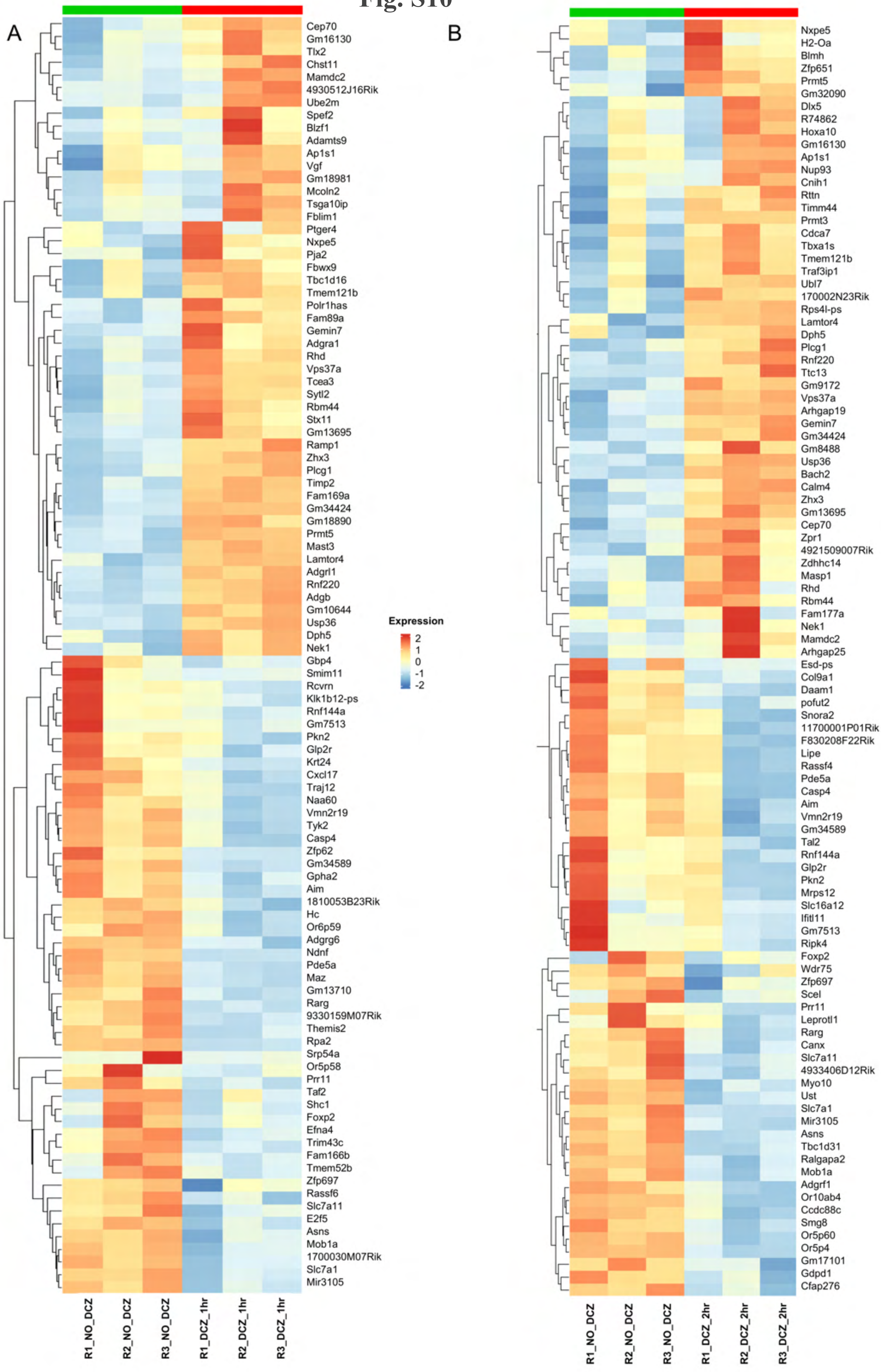
A
Cep70
Gm16130
Tlx2
Chst11
Mamdc2
4930512J16Rik
Ube2m
Spef2
Blzf1
Adamts9
Ap1s1
Vgf
Gm18981
Mcoln2
Tsga10ip
Fblim1
Ptger4
Nxpe5
Pja2
Fbwx9
Tbc1d16
Tmem121b
Polr1has
Fam89a
Gemin7
Adgra1
Rhd
Vps37a
Tcea3
Sytl2
Rbm44
Stx11
Gm13695
Ramp1
Zhx3
Plcg1
Timp2
Fam169a
Gm34424
Gm18890
Prmt5
Mast3
Lamtor4
Adgrl1
Rnf220
Adgb
Gm10644
Usp36
Dph5
Nek1
Gbp4
Smim11
Rcvrn
Klk1b12-ps
Rnf144a
Gm7513
Pkn2
Glp2r
Krt24
Cxcl17
Traj12
Naa60
Vmn2r19
Tyk2
Casp4
Zfp62
Gm34589
Gpha2
Aim
1810053B23Rik
Hc
Or6p59
Adgrg6
Ndnf
Pde5a
Maz
Gm13710
Rarg
9330159M07Rik
Themis2
Rpa2
Srp54a
Or5p58
Prr11
Taf2
Shc1
Foxp2
Efna4
Trim43c
Fam166b
Tmem52b
Zfp697
Rassf6
Slc7a11
E2f5
Asns
Mob1a
1700030M07Rik
Slc7a1
Mir3105
R1_NO_DCZ
R2_NO_DCZ
R3_NO_DCZ
R1_DCZ_1hr
R2_DCZ_1hr
R3_DCZ_1hr
Expression
2
1
0
-1
-2
B
Nxpe5
H2-Oa
Blmh
Zfp651
Prmt5
Gm32090
Dlx5
R74862
Hoxa10
Gm16130
Ap1s1
Nup93
Cnih1
Rttn
Timm44
Prmt3
Cdca7
Tbxa1s
Tmem121b
Traf3ip1
Ubl7
170002N23Rik
Rps4l-ps
Lamtor4
Dph5
Plcg1
Rnf220
Ttc13
Gm9172
Vps37a
Arhgap19
Gemin7
Gm34424
Gm8488
Usp36
Bach2
Calm4
Zhx3
Gm13695
Cep70
Zpr1
4921509007Rik
Zdhhc14
Masp1
Rhd
Rbm44
Fam177a
Nek1
Mamdc2
Arhgap25
Esd-ps
Col9a1
Daam1
pofut2
Snora2
11700001P01Rik
F830208F22Rik
Lipe
Rassf4
Pde5a
Casp4
Aim
Vmn2r19
Gm34589
Tal2
Rnf144a
Glp2r
Pkn2
Mrps12
Slc16a12
Ifitl11
Gm7513
Ripk4
Foxp2
Wdr75
Zfp697
Scel
Prr11
Leprotl1
Rarg
Canx
Slc7a11
4933406D12Rik
Myo10
Ust
Slc7a1
Mir3105
Asns
Tbc1d31
Ralgapa2
Mob1a
Adgrf1
Or10ab4
Ccdc88c
Smg8
Or5p60
Or5p4
Gm17101
Gdpd1
Cfap276
R1_NO_DCZ
R2_NO_DCZ
R3_NO_DCZ
R1_DCZ_2hr
R2_DCZ_2hr
R3_DCZ_2hr

### Fig. S11

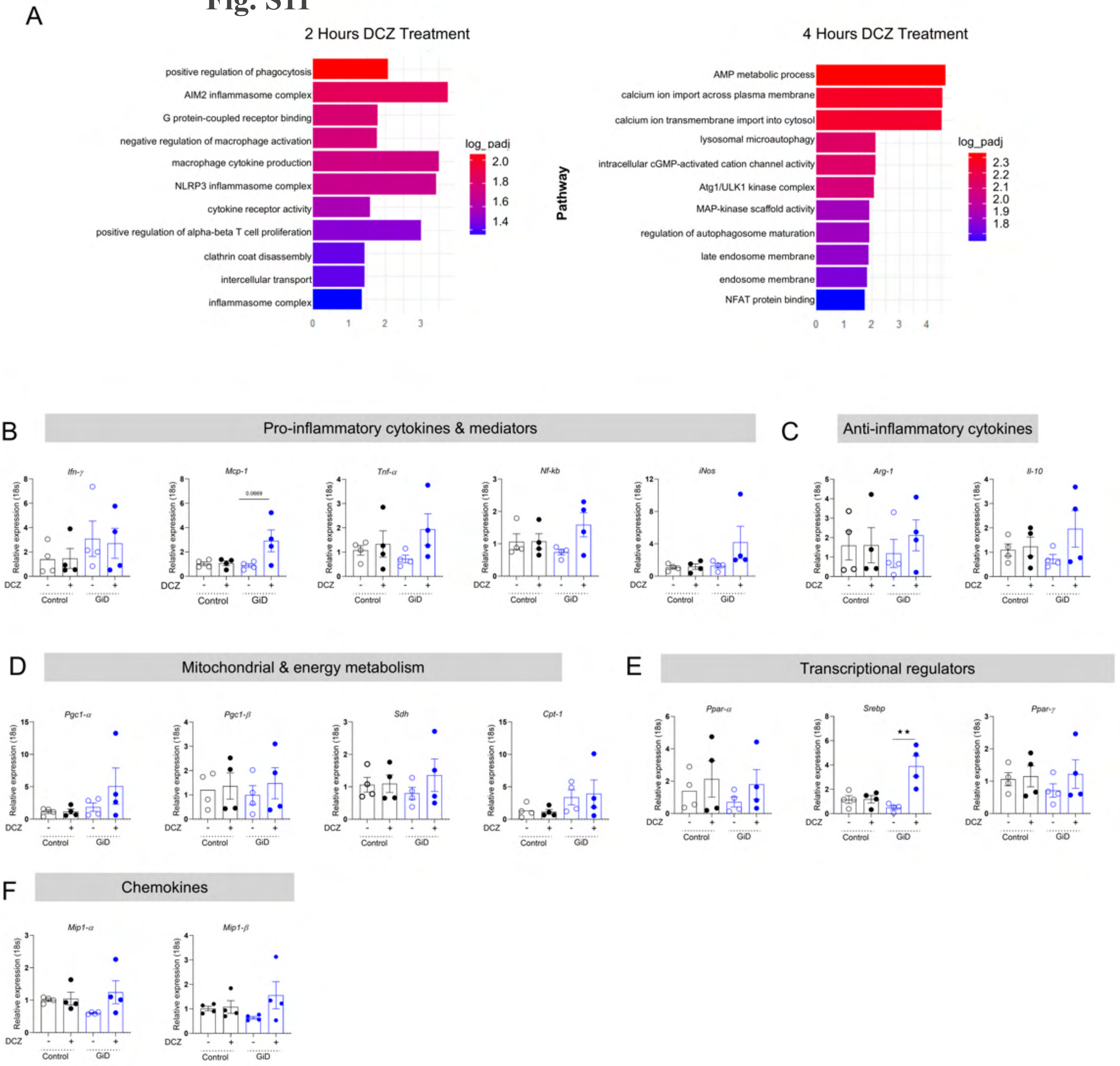
A
2 Hours DCZ Treatment
positive regulation of phagocytosis
AIM2 inflammasome complex
G protein-coupled receptor binding
negative regulation of macrophage activation
macrophage cytokine production
NLRP3 inflammasome complex
cytokine receptor activity
positive regulation of alpha-beta T cell proliferation
clathrin coat disassembly
intercellular transport
inflammasome complex
log_padj
Pathway
4 Hours DCZ Treatment
AMP metabolic process
calcium ion import across plasma membrane
calcium ion transmembrane import into cytosol
lysosomal microautophagy
intracellular cGMP-activated cation channel activity
Atg1/ULK1 kinase complex
MAP-kinase scaffold activity
regulation of autophagosome maturation
late endosome membrane
endosome membrane
NFAT protein binding
B
Pro-inflammatory cytokines & mediators
Ifn-γ
Mcp-1
0.0669
Tnf-α
Nf-kb
iNos
C
Anti-inflammatory cytokines
Arg-1
Il-10
D
Mitochondrial & energy metabolism
Pgc1-α
Pgc1-β
Sdh
Cpt-1
E
Transcriptional regulators
Ppar-α
Srebp
Ppar-γ
F
Chemokines
Mip1-α
Mip1-β
Relative expression (18s)
DCZ
Control
GiD

**Fig. S12**

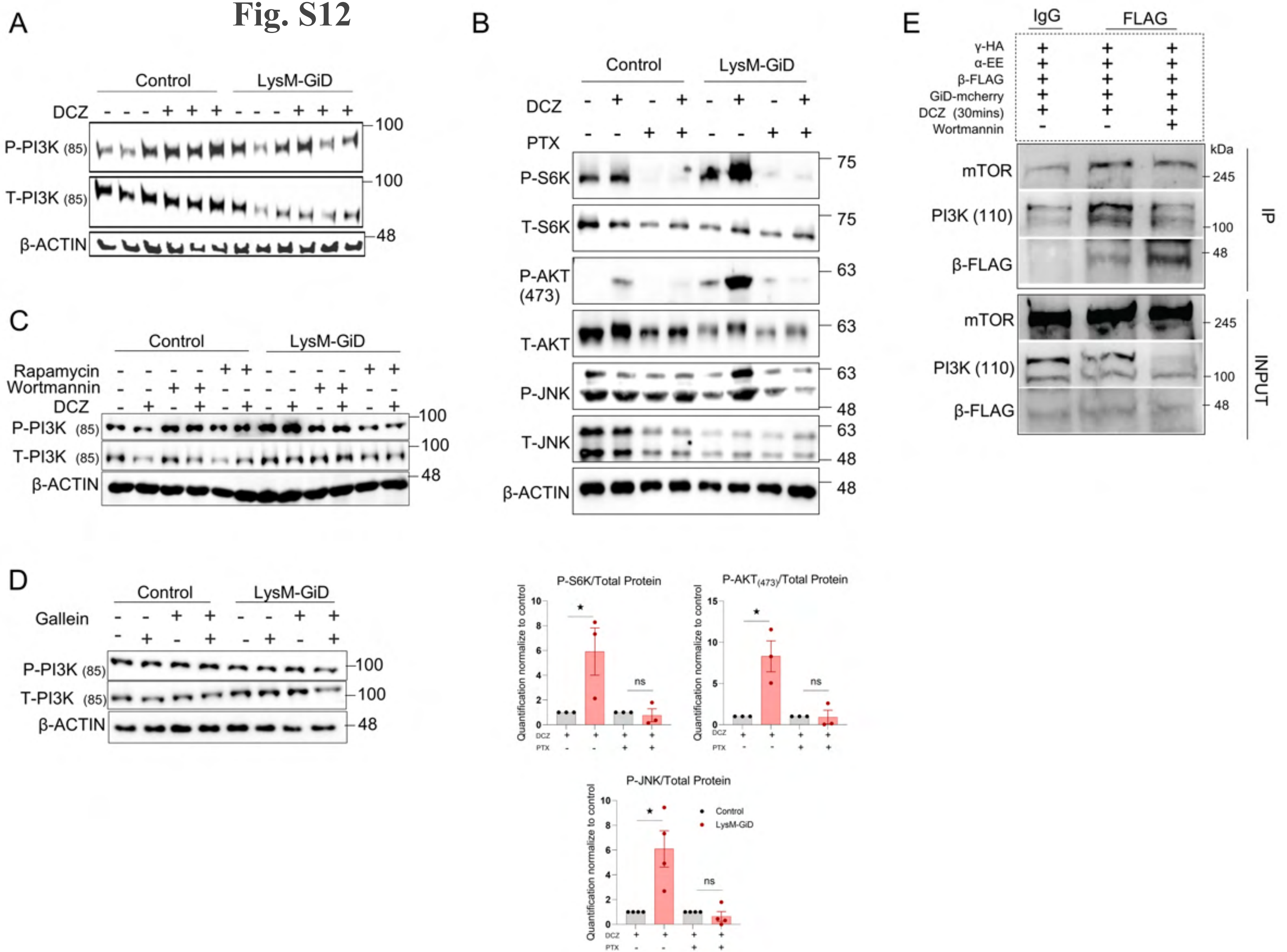
A
Control
LysM-GiD
DCZ
P-PI3K (85)
T-PI3K (85)
β-ACTIN
100
100
48
B
Control
LysM-GiD
DCZ
PTX
P-S6K
T-S6K
P-AKT (473)
T-AKT
P-JNK
T-JNK
β-ACTIN
75
75
63
63
63
48
63
48
48
C
Control
LysM-GiD
Rapamycin
Wortmannin
DCZ
P-PI3K (85)
T-PI3K (85)
β-ACTIN
100
100
48
D
Control
LysM-GiD
Gallein
P-PI3K (85)
T-PI3K (85)
β-ACTIN
100
100
48
E
IgG
FLAG
γ-HA
α-EE
β-FLAG
GiD-mcherry
DCZ (30mins)
Wortmannin
kDa
mTOR
PI3K (110)
β-FLAG
245
100
48
IP
INPUT
P-S6K/Total Protein
P-AKT(473)/Total Protein
P-JNK/Total Protein
Quantification normalize to control
DCZ
PTX
ns
Control
LysM-GiD

### Fig. S13

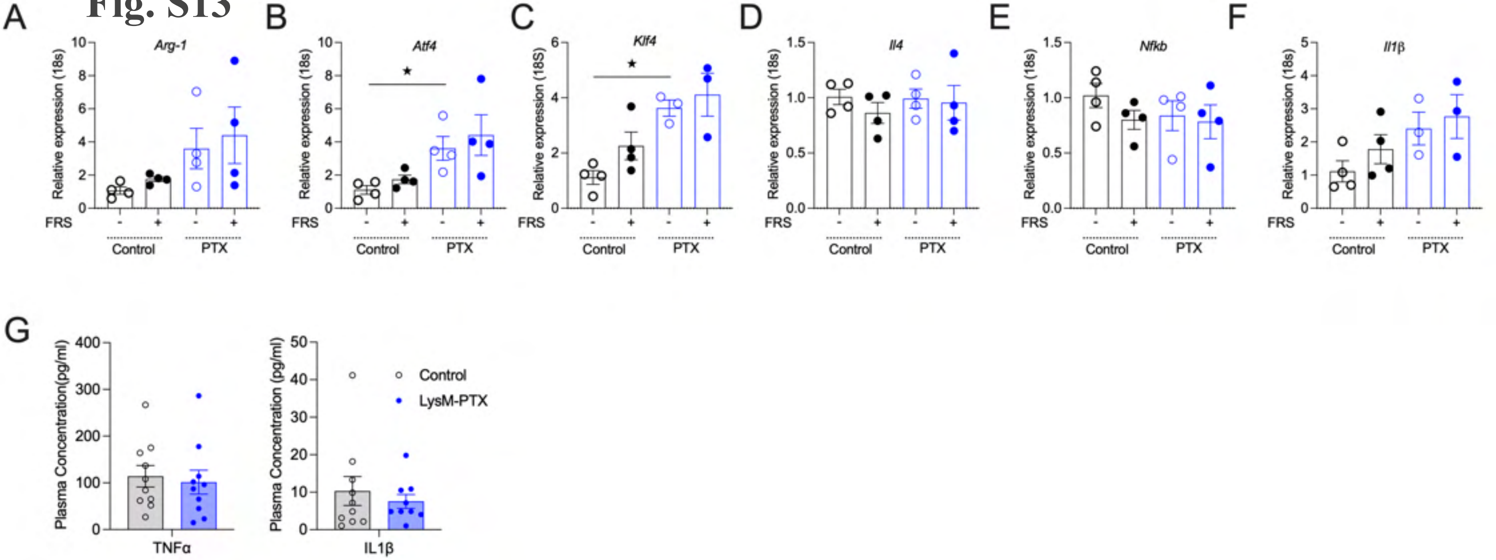
A
Arg-1
Relative expression (18s)
FRS
Control
PTX
B
Atf4
Relative expression (18s)
C
Klf4
Relative expression (18S)
D
Il4
E
Nfkb
F
Il1β
G
Plasma Concentration(pg/ml)
TNFα
Plasma Concentration (pg/ml)
IL1β
Control
LysM-PTX

### Fig. S14

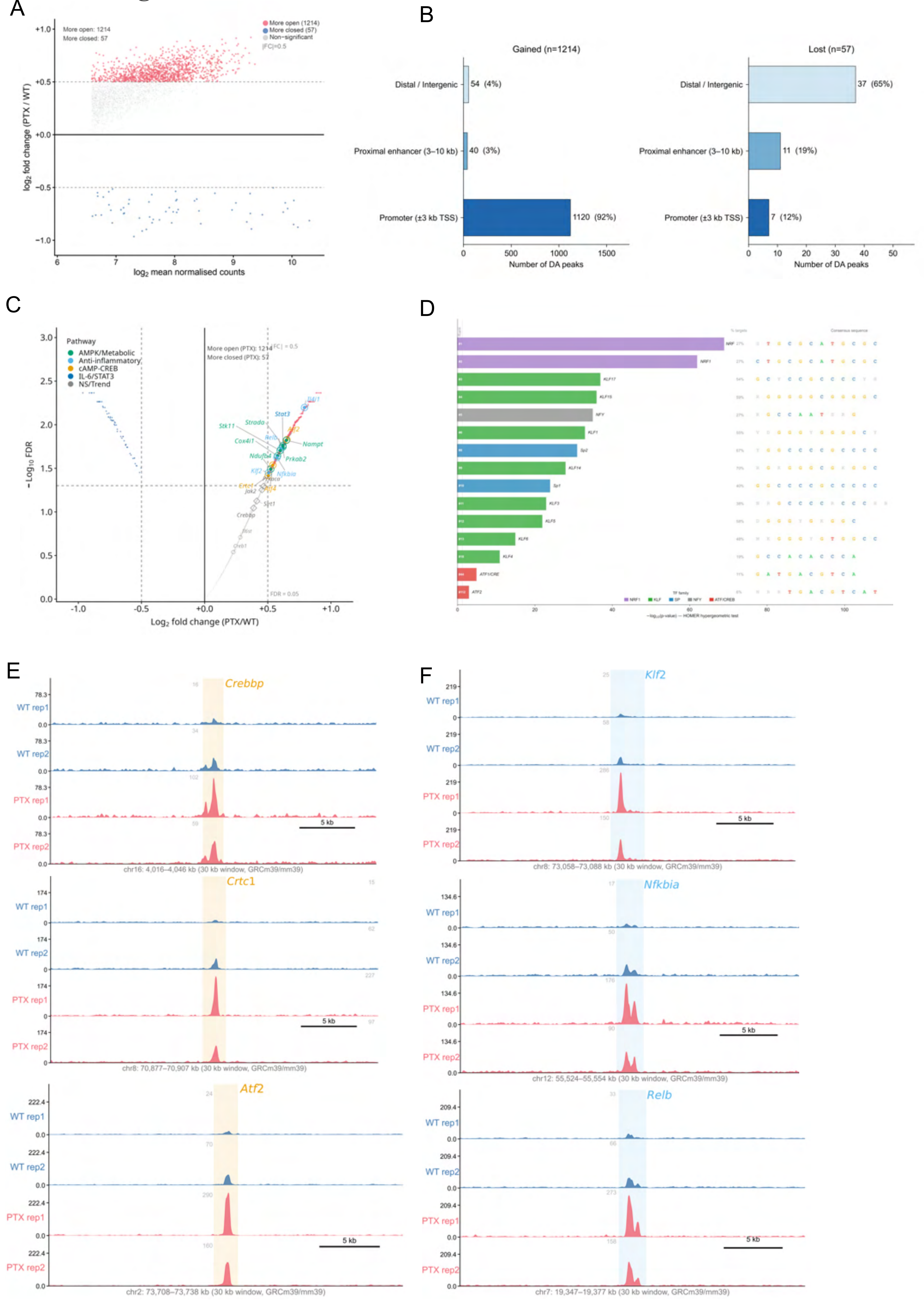
A
More open: 1214
More closed: 57
More open (1214)
More closed (57)
Non-significant
|FC|=0.5
log2 fold change (PTX / WT)
log2 mean normalised counts
B
Gained (n=1214)
Lost (n=57)
Distal / Intergenic
Proximal enhancer (3–10 kb)
Promoter (±3 kb TSS)
54 (4%)
40 (3%)
1120 (92%)
37 (65%)
11 (19%)
7 (12%)
Number of DA peaks
C
Pathway
AMPK/Metabolic
Anti-inflammatory
cAMP-CREB
IL-6/STAT3
NS/Trend
More open (PTX): 1214
More closed (PTX): 57
−Log10 FDR
Log2 fold change (PTX/WT)
FDR = 0.05
D
E
Crebbp
Crtc1
Atf2
WT rep1
WT rep2
PTX rep1
PTX rep2
5 kb
chr16: 4,016–4,046 kb (30 kb window, GRCm39/mm39)
chr8: 70,877–70,907 kb (30 kb window, GRCm39/mm39)
chr2: 73,708–73,738 kb (30 kb window, GRCm39/mm39)
F
Klf2
Nfkbia
Relb
chr8: 73,058–73,088 kb (30 kb window, GRCm39/mm39)
chr12: 55,524–55,554 kb (30 kb window, GRCm39/mm39)
chr7: 19,347–19,377 kb (30 kb window, GRCm39/mm39)
